# Adrenomedullin restores the human cortical interneurons migration defects induced by hypoxia

**DOI:** 10.1101/2023.05.01.538334

**Authors:** Alyssa Puno, Wojciech P. Michno, Li Li, Amanda Everitt, Kate McCluskey, Saw Htun, Dhriti Nagar, Jong Bin Choi, Yuqin Dai, Seyeon Park, Emily Gurwitz, A. Jeremy Willsey, Fikri Birey, Anca M. Pasca

## Abstract

Extremely preterm birth (at < 28 postconceptional weeks) leads to brain injury and represents the leading cause of childhood-onset neuropsychiatric diseases. No effective therapeutics exist to reduce the incidence and severity of brain injury of prematurity. Hypoxic events are the most important environmental factor, along with inflammation. Among other developmental processes, the second half of in utero fetal development coincides with the migration of cortical interneurons from the ganglionic eminences into the cortex; this process is thus prone to disruptions following extremely preterm birth. To date, no studies have directly investigated the migration of human cortical inhibitory neurons under hypoxic conditions. Using multi-day confocal live imaging in human forebrain assembloids (hFA) derived from human induced pluripotent stem cells (hiPSCs) and *ex vivo* developing human brain tissue, we found a substantial reduction in the migration of hypoxic interneurons. Using transcriptomics, we identified adrenomedullin (*ADM*) as the gene with the highest fold change increase in expression. Based on previous literature about the protective role of supplemental ADM for other injuries, here, we demonstrated that addition of exogenous ADM to the hypoxic media restores the migration defects of interneurons. Lastly, we showed that one of the mechanisms of protection by ADM is through the activation of the cAMP/PKA pathway and subsequent pCREB-dependent rescued expression of a subset of GABA receptors, which are known to promote migration. Overall, in this manuscript we provide the first direct evidence for hypoxia-induced deficits in the migration of human cortical interneurons and identify ADM as a possible target for therapeutic development.

**Summary:** We identified human cortical interneurons migration deficits under hypoxic conditions and rescued this phenotype using adrenomedullin.

## INTRODUCTION

Extremely preterm birth (at < 28 postconceptional weeks) is commonly associated with brain injury of prematurity, and represents the leading risk factor for childhood-onset neuropsychiatric diseases^1–3^. Clinical studies have identified postnatal hypoxic events as one of the most important environmental risk factors for brain injury of prematurity^4^. Despite important clinical progress in understanding brain injury of prematurity from a histologic and radiologic perspective, the cell type-specific biological mechanisms of injury remain largely understudied in humans.

Among other phenotypes^5^, histology studies demonstrate a substantial reduction in the number of cortical interneurons in post-mortem tissue samples from individuals previously born extremely premature^6^. This decrease is now suggested as a significant contributor to the pathophysiology of neuropsychiatric disorders associated with preterm birth^7–10^. Cortical interneurons are born and proliferate in the ganglionic eminences before migrating to the cortex, where they eventually become part of the cortical synaptic networks in a process that evolves over multiple years in humans^11^. Of all these developmental stages, migration toward the cortex is the process most active during the second half of in utero development, which, in extremely preterm infants is replaced by postnatal hospitalization in the Neonatal ICU and associated with hypoxic events due to lung immaturity and inability to transfer oxygen from the environment to the tissues.

Despite migration defects being highly suspected to be impaired in hypoxic brain injury of prematurity, the investigation of this process in preclinical models has been uniquely challenging to study, because this migration mostly happens in utero in most species. As a result, to date, no studies have yet been able to directly visualize the migration patterns of cortical interneurons upon exposure to hypoxia. Moreover, no in- depth data exist about the molecular mechanisms of injury in hypoxic migrating interneurons, thus making therapeutic discovery challenging. Previous animal models for hypoxic brain injury of prematurity have, however, replicated the human-observed histologic decrease in cortical interneurons in the dorsal forebrain of pre-born embryos as early as 4 days after exposing the pregnant dam to a hypoxic injury, and suggested that this decrease is related to a defect in the migration of interneurons that have already exited the ganglionic eminences at the time of hypoxia exposure^12^. These findings are encouraging and support the hypothesis that migration is affected by hypoxia and highlight the need for further investigation using newly emerging scientific tools.

To study the effects of hypoxia on the migration of cortical interneurons and advance our mechanistic understanding of prematurity-associated interneuronopathies, we developed two complementary human cellular models which consist of a stem cell-based *in vitro* human cellular platform, and an *ex vivo* developing human brain tissue platform. Specifically, we established a multi-day confocal live imaging setup for the visualization of migrating cortical interneurons within human forebrain assembloids (hFAs) derived from induced pluripotent stem cells (hiPSCs)^13,14^, and within *ex vivo* developing human brain tissue.

Using these new models, we identified substantial migration deficits during exposure to hypoxia, thus providing the first direct evidence for the long-proposed migration delays induced by hypoxia. These findings are important because delays in reaching the specified cortical targets within the developmentally appropriate window are known to lead to elimination of cortical interneurons through apoptosis, and the direct demonstration of this deficit strengthens the hypothesis that delayed migration contributes to the decrease in the number of cortical interneurons observed in post-mortem brain tissues from preterm individuals.

Additionally, we demonstrated that supplementation with exogenous ADM peptide restores the migration cortical interneurons during exposure to hypoxia. Adrenomedullin peptide (ADM) has been previously shown to acutely increase in various non-neurologic and neurologic hypoxic and inflammatory conditions, in both clinical and preclinical studies^15–17^. Moreover, it has been demonstrated to be a direct binding partner for HIF1α, which is a hallmark of hypoxia, but is also stabilized in inflammation^18^. Interestingly, ongoing clinical trials for the treatment of inflammatory bowel disease have shown therapeutic promise and safety upon administration of exogenous ADM at supraphysiological levels^19–22^. Our data, in conjunction with previous findings, suggest exogenous administration of ADM as a therapeutic intervention should be further investigated in vivo and for various hypoxic injuries.

## RESULTS

### Development of a human cellular model to study the migration of hypoxic cortical interneurons

The hFAs recapitulate key developmental aspects of cortical interneurons migration from the medial ganglionic eminences (MGE) into the dorsal forebrain. To study the effects of hypoxia on the migration of human cortical interneurons, we established a model for hypoxic injury using hFAs. Using standardized published protocols we previously contributed to developing^13,14^, we generated hFAs through the differentiation of hiPSCs into human cortical organoids (hCO) containing dorsal forebrain excitatory neurons^23^, and human subpallial organoids (hSO) containing cortical interneurons^14^. To visualize migrating cortical interneurons, we first fluorescently tagged cells within hSOs with a forebrain interneurons cell-type specific lentiviral reporter (Dlxi1/2b::eGFP)^24^. Subsequently, we fused the labeled hSOs with hCOs to generate hFAs. At approximately 10-14 days after fusion into hFAs, we observed substantial and active migration of Dlxi1/2b::eGFP^+^ cortical interneurons on the hCO side of hFAs (Fig 1A, Fig. S1A).

**Figure 1.**
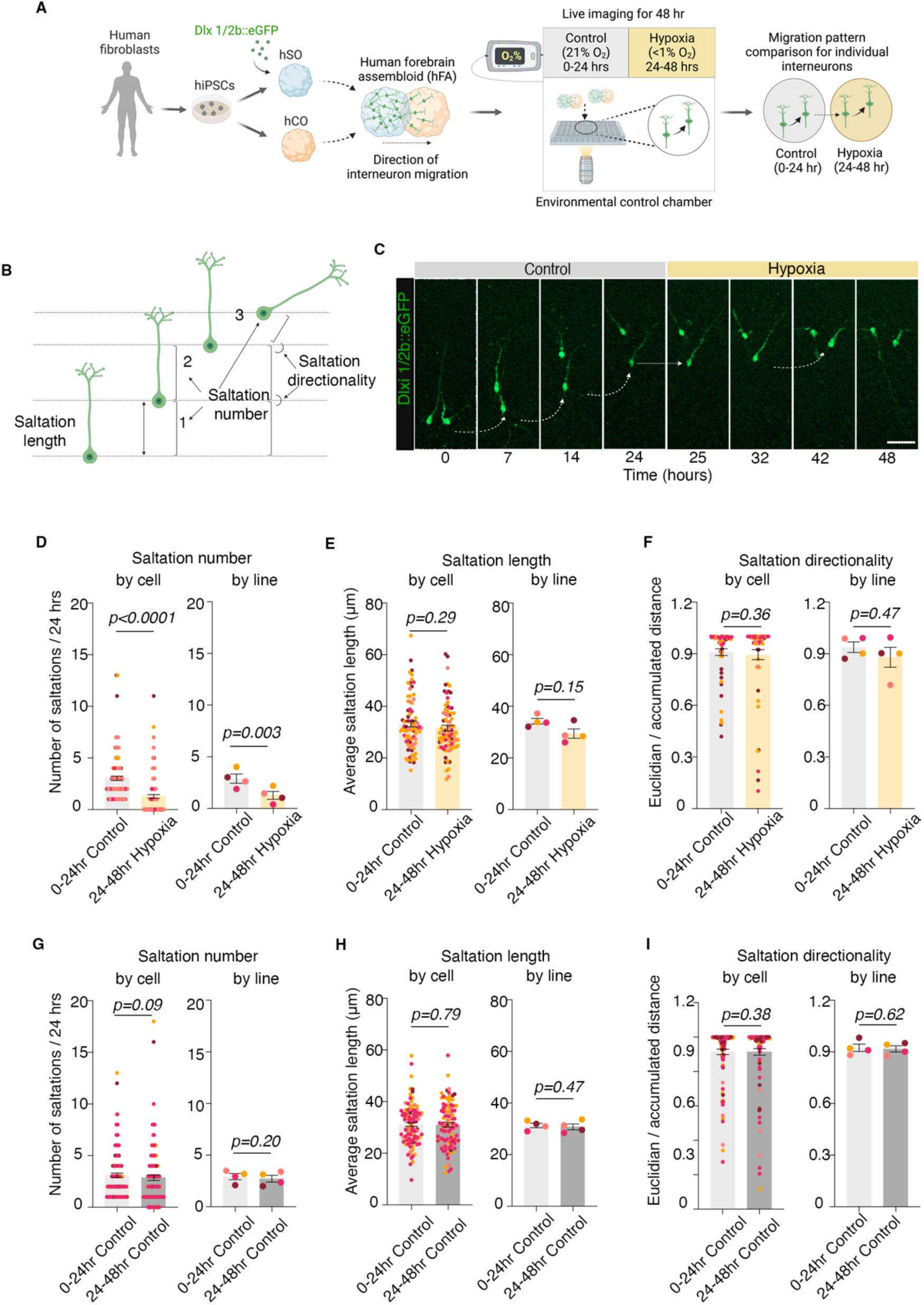
Human cellular model to study migration patterns in cortical interneurons under hypoxic stress. **A.** Schematic illustrating overall experimental design: hiPSCs were used to derive human cortical organoids (hCO) and human subpallial organoids (hSO); for direct visualization of migrating interneurons, at ∼45-55 days in culture hSO were infected with lentivirus Dlxi1/2b::eGFP and then fused with hCO into human forebrain assembloid (hFA); to study interneuron migration during exposure to hypoxia, hFA were imaged 10-14 days post infection using confocal live-imaging setup focused on the hCO part of the hFA; the movement of the same cells was followed for a total of 48 hrs, each condition for 24hrs: 0-24 hrs in control and 24-48 hrs in hypoxia; **B.** Schematic of migratory pattern of interneurons, focused on number of saltations, average saltation length and directionality; **C.** Example of migration pattern of one cortical interneuron during control and hypoxic conditions; **D.** Quantification of saltations number/24 hrs in hypoxia-exposed and non-exposed cortical interneurons by individual cells (paired Wilcoxon test, P<0.0001) and by hiPSC line (two-tailed paired *t*-test, P=0.003); **E.** Quantification of average saltation length for hypoxia-exposed and non-exposed cortical interneurons by individual cells (paired Wilcoxon test, P=0.29) and by hiPSC line (two-tailed paired *t*-test, P=0.15); **F.** Quantification of directionality of migration in hypoxia-exposed and non-exposed cortical interneurons by individual cells (paired Wilcoxon test, P=0.36), and by hiPSC cell line (two-tailed paired *t*-test, P=0.47); **G.** Quantification of saltations number/24 hrs in control conditions, in the first 24 hrs (0-24 hrs) versus the subsequent 24 hrs (24-48 hrs) of live imaging by individual cells (paired Wilcoxon test, P=0.09) and hiPSC line (two-tailed paired *t*-test, P=0.2); **H.** Quantification of average saltation length in control conditions, in the first 24 hrs (0-24hrs) versus the subsequent 24 hrs (24-48 hrs) of live imaging by individual cells (two-tailed paired *t*-test, P=0.79) and by hiPSC line (two-tailed paired *t*-test, P=0.47; **I.** Quantification of directionality of migration in control conditions, in the first 24 hrs (0-24 hrs) versus the subsequent 24 hrs (24-48 hrs) of live imaging by individual cells (paired Wilcoxon test, P=0.38) and by hiPSC line (two-tailed paired *t*-test, P=0.62); Bar charts: mean ± s.e.m; scale bar: 50 μm.

To monitor the migration patterns of cortical interneurons under control and hypoxic stress, we established a long-term live imaging microscopy setup in intact hFAs using a confocal microscope with a motorized stage (Zeiss LSM 980 with Airyscan 2) equipped with an environmentally controlled chamber (37°C, 5% CO_2_) and an oxygen level controller (Okolab Bold Line). For this, the hFAs were transferred to the confocal microscope in a glass-bottom 96-well plate with fresh cell culture media. We imaged individual Dlxi1/2b::eGFP-tagged interneurons within the hCOs for a total of 48 hrs: 24 hrs (0-24 hrs) under control conditions (37°C, 5% CO_2,_ 21% O_2_), followed by 24 hrs (24-48 hrs) in pre-equilibrated hypoxic media (37°C, 5% CO_2_, 94.5% N_2_, <1% O_2_) (Fig. 1A).

To validate the decrease in partial pressure of O_2_ (PO_2_) in the culture media in hypoxic conditions, we used an oxygen optical microsensor OXB50 (50 μm, PyroScience) attached to a fiber-optic multi-analyte meter (FireStingO_2_, PyroSciences). In line with our previous reports^25^, we observed a decrease in the media PO_2_ level from ∼150 mmHg in control conditions, to ∼25-30 mmHg in the confocal microscope environmental chamber in hypoxic conditions (Fig. S1B).

Next, we confirmed that these conditions are sufficient to activate the hypoxia response pathway in hSOs within 24 hrs of exposure, without inducing extensive death of cortical interneurons. Specifically, to validate the induction of hypoxia, we demonstrated the stabilization of HIF1α hypoxia-marker protein using western blot analyses in hSOs derived from 4 individual cell lines (two-tailed paired *t*-test, P=0.01) (Fig. S1C,D). Separately, we validated the activation of the hypoxia-response pathway using targeted transcriptional analyses for hypoxia-responsive genes, including *PFKP* (two-tailed paired *t*-test, P=0.004), *PDK1* (two-tailed paired *t*-test, P=0.001), *VEGFA* (two-tailed paired *t*-test, P=0.003) (Fig. S1E, Table S2). To check whether Dlxi1/2b::eGFP-tagged interneurons start expressing cell markers for cell death, we used two different assays: we performed FACS-based Annexin V analyses in fluorescently tagged interneurons, as well as immunostainings for cleaved caspase 3 (c-CAS3). We found no significant difference between control and hypoxic conditions using Annexin V (unpaired *t-*test, P=0.362) (Fig. S1F) nor c-CAS3 (unpaired *t-*test, P=0.125) (Fig. S1G,H).

These results demonstrate that our target hypoxic conditions (PO_2_ ∼25-30 mmHg) are sufficient to induce and activate the hypoxia response pathway, but do not induce cell death. This experimental design aligns with our goal of targeting mild hypoxic stress, partly because mild hypoxic episodes are much more common than the severe ones in preterm infants, and partly because we wanted to define, for the first time, the altered molecular pathways in hypoxic (but live) cortical interneurons.

### Human cortical interneurons display migration deficits upon exposure to hypoxia

The migration of cortical interneurons from the ganglionic eminences into the dorsal forebrain is at its peak during the second half of pregnancy in humans^26^. It is characterized by repetitive cycles of nuclear saltations (nucleokinesis events) in the direction of the leading process^27,28^. To study migration patterns of hypoxic cortical interneurons we followed the movement of the same individual Dlxi1/2b::eGFP^+^ cortical interneurons for a total of 48 hrs (control conditions: hours 0-24; hypoxia: hours 24-48). We analyzed the number of nuclear saltations, saltation length and the directionality of the migration in interneurons that had already migrated into the hCO at the time of the hypoxia exposure (Fig. 1B, C).

To quantify the number of saltations, we focused on interneurons that had at least one saltation within the first 24 hours under control conditions and remained visible throughout the entire 48 hrs of imaging. This experimental approach was specifically chosen to focus the analyses on actively migrating cells, while excluding those that may have ceased migration and begun integrating into synaptic networks at the time of imaging. To quantify saltation length and directionality, we narrowed the cell population to the subset of cells that had at least one saltation in both control and hypoxic conditions, as a minimum of 1 saltation/condition is a prerequisite for direct comparisons for these parameters.

To identify defects in the migration patterns of interneurons in hypoxia, we analyzed migrating interneurons for 24 hrs in control conditions (hours 0-24), followed by another 24 hrs in hypoxic conditions (hours 24-48). We observed a ∼58% decrease in the number of saltations of individual interneurons (paired Wilcoxon test, P<0.0001), and demonstrated that the phenotype is present in hFAs derived using hiPSCs from 4 separate individuals (two-tailed paired *t*-test, P=0.003) (Fig. 1D). Importantly, 70 out of 129 actively migrating cortical interneurons, not only decreased the number of saltations, but became completely stationary upon exposure to hypoxia for 24 hrs. However, we found no significant differences in the average length of saltation of cells when analyzed by individual cells (paired Wilcoxon test, P=0.29) and by hiPSC line (two-tailed paired *t*-test, P=0.15) (Fig. 1E). In addition, we did not observe any differences in the directionality of migration measured by the ratio of Euclidean/accumulated distance when analyzed by individual cells (paired Wilcoxon test, P=0.36) and by hiPSC line (two-tailed paired *t*-test, P=0.47) (Fig. 1F).

To ensure that the confocal chamber environment during the prolonged live imaging does not result by itself in defects in migration of the cortical interneurons, or that a possible hypoxia phenotype could be related to a physiologic saltatory pause in the second half of imaging, we performed two separate types of experiments. First, we compared the migration patterns in control conditions (37°C, 5% CO_2,_ 21% O_2_) in the first 24 hrs (hours 0-24) versus the second 24 hrs (hours 24-48). We identified no differences in the total number of saltations/24 hrs when analyzed by individual cells (paired Wilcoxon test, P=0.09) and hiPSC line (two-tailed paired *t*-test, P=0.2) (Fig. 1G); no difference in average saltation length when analyzed by individual cells (two-tailed paired *t*-test, P=0.79) and by hiPSC line (two-tailed paired *t*-test, P=0.47) (Fig. 1H); no difference in directionality of migration when analyzed by individual cells (paired Wilcoxon test, P=0.38) and by hiPSC line (two-tailed paired *t*-test, P=0.62) (Fig. 1I). Second, to ensure that the deliberate exclusion from analyses of cortical interneurons that display no saltations in control conditions do not bias the migration phenotype by excluding cells that might otherwise migrate more in the 24-48 hrs of imaging, we identified all cortical interneurons that displayed no migration in control conditions (0-24 hrs) and quantified their migration in hypoxia. Out of 113 total cells, 108 (95.6%) remained stationary (Fig. S1I), suggesting these cells are not in a highly migratory phase at the time of imaging.

Overall, the experiments in Fig. 1 and S1 suggest mild hypoxia preferentially affects the total migration length of cortical interneurons by decreasing the number of saltations, however it does not interfere with the length of individual saltations nor the directionality of the migration. The additional validation experiments demonstrate the robustness of the experimental design and strengthen the validity of saltation deficit phenotype in hypoxia. Details on the number of cells, experiments and number of hiPSC lines used for each experiment are presented in Table S1.

### Transcriptional analyses in hypoxic hSOs suggest adrenomedullin (ADM) as a key component of hypoxia-induced cellular response

To investigate the transcriptional changes induced by mild hypoxia in hSOs, we performed bulk transcriptome-wide RNA-Sequencing (RNA-Seq) at 46 days in culture (before fusion with hCOs), following exposure to hypoxia for 12 hrs or 24 hrs, as well as at 72 hrs after reoxygenation (Fig. 2A). For these experiments we induced hypoxia using the C-chamber hypoxia sub-chamber (Biospherix). We confirmed a significant decrease in PO_2_ between control and hypoxia conditions (two-tailed paired *t*-test, P<0.0001) (Fig. S2A), and showed that PO_2_ levels were similar to the ones measured in the confocal environmental chamber (Fig. S1B).

**Figure 2.**
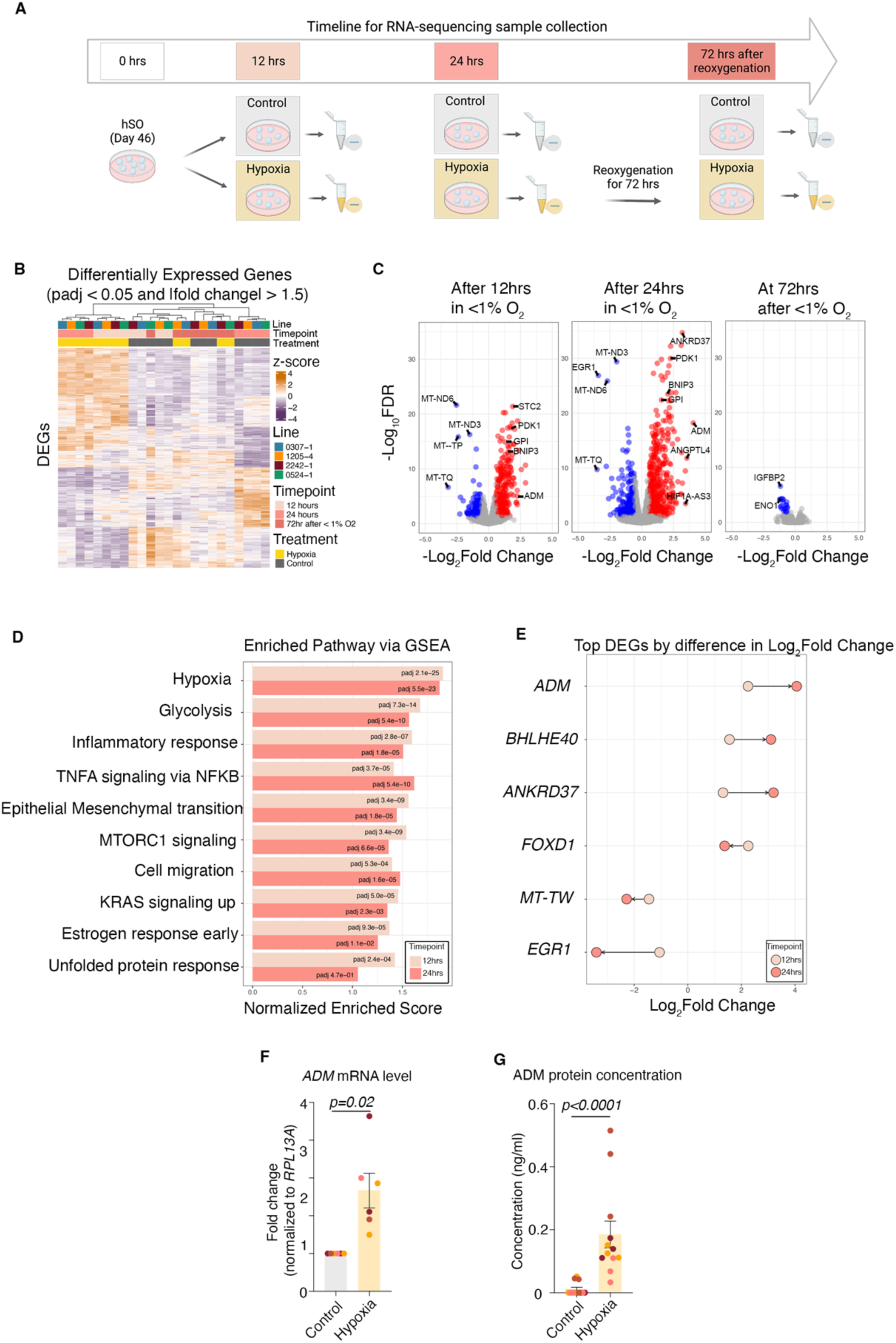
Transcriptional changes in hSO exposed to hypoxia. **A.** Schematic of hypoxia exposure of hSO and collection of samples for RNA-Sequencing; **B.** Heatmap of differentially expressed genes in RNA-Seq data showing clear transcriptional changes in hypoxia-exposed samples. Samples (*n* = 24) from hSO differentiated from 4 hiPSC lines were collected at 12 hrs and 24 hrs of exposure to hypoxia, as well as 72 hrs after reoxygenation. The union of all differentially expressed genes (*n* = 1,473) and samples (*n* = 24) are ordered by hierarchical clustering (complete linkage of Euclidean distance). Z-score normalized expression values are depicted on a continuous scale from lower values (purple) to higher values (orange). Cell lines, treatment and time point are depicted at the top and represented by different colors; **C.** Volcano plots of differentially expressed genes (DEGs) at 12 hrs, 24 hrs and 72 hrs after reoxygenation. Each dot represents a single gene. DEGs with a padj <0.05 and an absolute fold change >1.5 are shown in red (upregulated) or blue (downregulated) and unchanged genes are shown in gray; **D.** Bar plot of the top 10 shared enriched gene pathways across hypoxic conditions. Adjusted p-values are depicted as text and colors represent the length of hypoxic exposure; **E.** Dumbbell plot of the top 6 DEGs with the largest positive difference and the largest negative difference in Log_2_ fold change between 12 and 24 hrs of hypoxia exposure; the arrow indicates the direction of change from 12 to 24 hrs. **F.** Transcriptional upregulation (by qPCR) of *ADM* gene in hSO samples exposed to 24 hrs of hypoxia: *ADM* (two-tailed paired *t*-test, P=0.02); **G.** Quantitative enzyme immunoassay analysis of adrenomedullin (ADM) peptide in media from hSO exposed and non-exposed to hypoxia (unpaired Mann-Whitney test, P<0.0001); for values below the minimal detection range of <0.01 ng/mL, value was approximated to 0 (we had 9 values approximated to 0 in the control samples). Bar charts: mean ± s.e.m.; Different dot colors represent individual hiPSC lines.

Hierarchical clustering of the gene expression profiles showed clear separation between hSOs from the control (control 12hr, control 24hr, control 72hr), hypoxia-exposed (hypoxia 12hr, hypoxia 24hr), and hypoxia followed by reoxygenation samples (hypoxia 72hr after <1%) (Fig. S2b), suggesting that hypoxia exposure induces strong transcriptional profile changes in hSOs, and these largely recover following 72 hrs of reoxygenation. For this experiment, we used hSOs derived from 4 individual hiPSC lines, two XX and two XY genotypes (details by hiPSC line are presented in Table S1).

Next, we identified differentially expressed genes (DEGs) between control, hypoxia-exposed, and reoxygenated samples (padj<0.05, fold change>1.5), controlling for potential confounding variables as described in detail in the methods section. We identified 734 differentially expressed genes at 12 hrs, 985 genes at 24 hrs, and 21 genes at 72 hrs after reoxygenation (Fig. 2b, 2c, Data File S1).

Gene set enrichment analyses (GSEA) demonstrated changes in molecular pathways associated with exposure to hypoxia (including glycolysis, mTORC1, KRAS), inflammatory response pathways (represented by nonspecific immune-related cell surface receptors), estrogen response pathways and unfolded protein response. In addition, we identified changes in genes associated with general cellular migration (e.g. *AXL*, *GLIPR1*, etc) (Fig. 2D) (Data File S2).

Interestingly, we found that the adrenomedullin (*ADM*) gene had the highest fold change among DEGs at 24 hrs of hypoxia exposure (log_2_FC=4.06, padj=6.08E-19) (Fig. 2E). To validate the increase in gene expression level of *ADM* under hypoxic conditions observed in bulk RNA-sequencing, we performed qPCR analyses using hypoxia-exposed intact hSOs. We confirmed the significant transcriptional upregulation of *ADM* gene expression (two-tailed paired *t*-test, P=0.02) (Fig. 2F).

To check whether the transcriptional upregulation of *ADM* gene expression translates into changes at a protein level, we quantified ADM peptide levels using quantitative an enzyme immunoassay (EIA) (Phoenix Pharmaceuticals, EK-010-01) in cell culture media from intact hSOs exposed and non-exposed to hypoxia for 24 hrs. We observed a significant increase in the protein concentration in the hypoxia-exposed hSOs compared to control conditions (unpaired Mann-Whitney test, P<0.0001) (Fig. 2g). This experiment was performed in 3 separate biological replicates of hSOs, each derived from 4 hiPSC lines in one differentiation experiment (details by hiPSC line are presented in Table S1).

### ScRNA-sequencing in hypoxic hFAs uncovers cell type-specific expression of *ADM* and its receptors

To bring novel insight into the cell type-specific expression of *ADM* under control and hypoxic conditions in human brain cells, we performed single-cell RNA-sequencing in hSOs and hCOs, separated from hFAs that were previously exposed or non-exposed to hypoxia. To do this, we cut the hFAs under microscopic guidance (Fig. 3A). We used hFAs derived from 2 different hiPSC lines.

**Figure 3.**
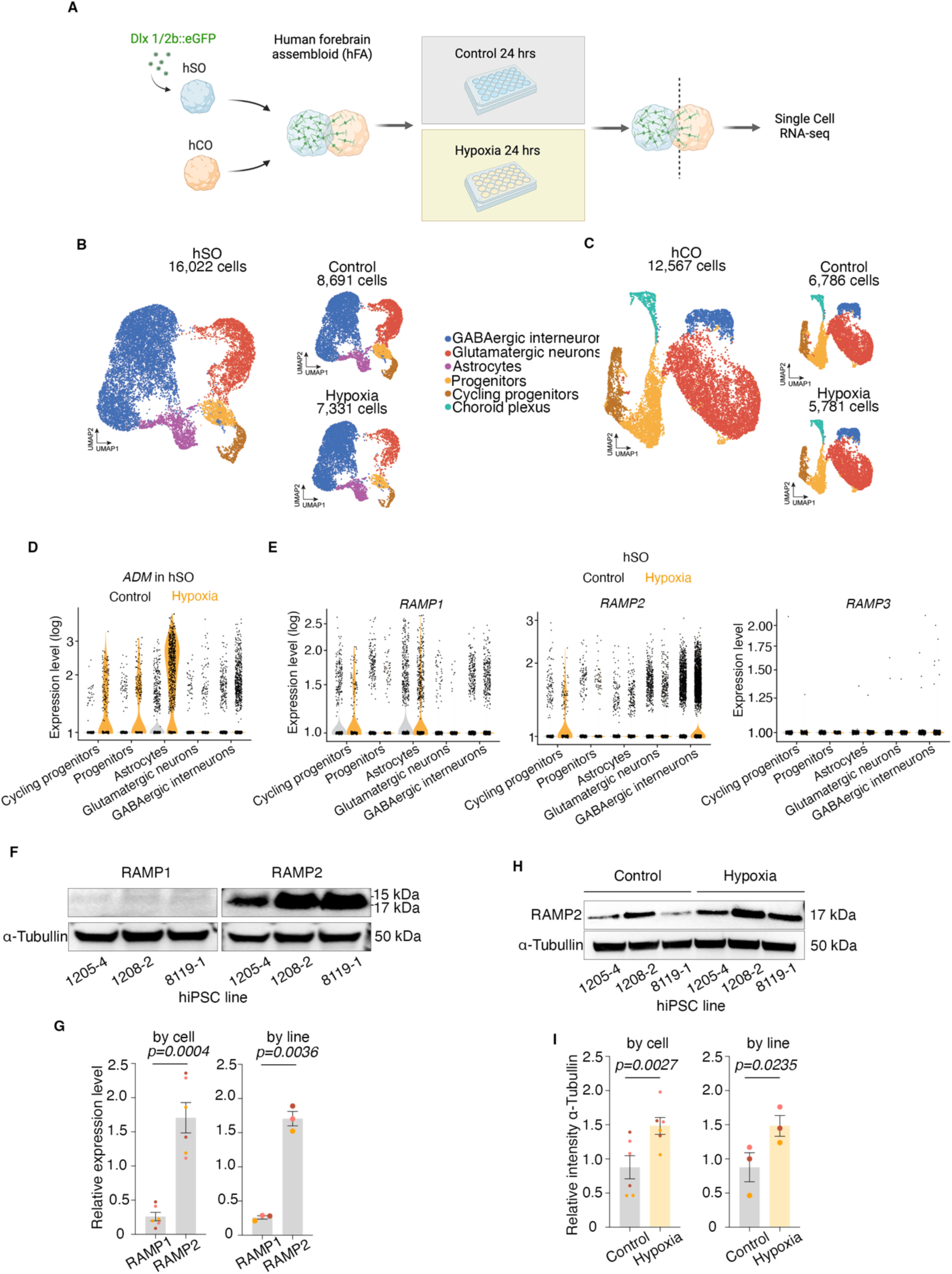
Single cell transcriptional profiling in control and hypoxia exposed hSO and hCO. Schematic of single cell RNA-seq of hCO and hSO from hFA; **B.** UMAP visualization of the resolved single cell RNA-seq data of hSO with assignment of main cell clusters, with control and hypoxia exposure shown by condition (*n[total]* = 16,022 cells, *n[control]* = 8,691 cells, *n[hypoxia]* = 7,331 cells); **C.** UMAP visualization of the resolved single cell RNA-seq data of hCO with assignment of main cell clusters, with control and hypoxia exposure shown by condition (*n[total]* = 12,567 cells, *n[control]* = 6,786 cells, *n[hypoxia]* = 5,781 cells); **D.** Single cell gene expression level (log) of adrenomedullin (*ADM*) in main cell clusters of hSO under control and hypoxia conditions; **E.** Single cell gene expression level (log) of *RAMP1* in main cell clusters of hSO under control and hypoxia conditions; Single cell gene expression level (log) of *RAMP2* in main cell clusters of hSO under control and hypoxia conditions; Single cell gene expression level (log) of *RAMP3* in main cell clusters of hSO under control and hypoxia conditions; **F. (Left)** Representative blot for RAMP1 protein expression in control conditions, **(Right)** Representative blot for RAMP2 protein expression in control conditions; normalized to ɑ-Tubulin **G.** Quantification of RAMP1 and RAMP2 protein expression by **(Left)** individual hSO sample (*two*-tailed paired test, P=0.0004) and **(Right)** by hiPSC line (*two*-tailed paired test, P=0.0036) in control conditions; **H.** Representative blots for RAMP2 protein expression changes in control and hypoxia conditions; **I.** Quantification of RAMP2 protein expression **(Left)** by individual hSO sample (*two*-tailed paired test, P=0.0027) and **(Right)** by hiPSC line (*two*-tailed paired test, P=0.0235) in control and hypoxia conditions. Bar charts: mean ± s.e.m.; Different dot colors represent individual hiPSC lines.

First, we performed Uniform Manifold Approximation and Projection (UMAP) dimensionality reduction for hSOs non-exposed (n = 8,691cells) and exposed to hypoxia (n = 7,331 cells). We identified five distinct cell clusters including ventral forebrain progenitors and early cortical interneurons (*NKX2.1*^+^, DLX2^+^, *GAD1*^+^), cycling progenitors (*TOP2A*^+^), astrocytes (*S100A10*^+^) and a small cluster of radial glia (*SOX9*^+^); as expected due to the previous fusion into hFA, we identified a cluster of cortical glutamatergic neurons (*STMN2*^+^, *NEUROD2*^+^) (Fig. 3B, Fig. S3A). For hCOs non-exposed (n = 6,786 cells) and exposed to hypoxia (n = 5,781cells), we also identified five cellular clusters consisting of dorsal forebrain progenitors (*SOX2*^+^), cycling progenitors (*TOP2A*^+^), cortical glutamatergic neurons (*STMN2*^+^, *NEUROD2*^+^), and a small cluster of choroid plexus cells (*TTR*^+^); as expected due to the migration of interneurons from the hSO side, we identified a cluster of cortical interneurons (*NKX2.1*^+^, *GAD1*^+^) (Fig. 3C, Fig. S3B). All these cell clusters from hSOs and hCOs demonstrated increased expression of hypoxia-related genes (PDK1, PFKP) (Fig. S3C, D). Moreover, we performed subcluster analyses of GABAergic interneurons on hSO, and identified that they primarily express markers for MGE and CGE including *NKX2.1*, *LHX6*, somatostatin (*SST*) or calretinin (*CALB2),* less calbindin *(CALB1)* and minimal parvalbumin (*PVALB*) and we validated these findings and performed immunocytochemistry for the following interneurons subtypes: somatostatin (*SST*), calbindin *(CALB)* and calretinin (*CALB2)* (Figure S3E, F).

Next, we assessed the expression of *ADM* in individual cell types. We found that under hypoxic conditions, the expression level of *ADM* increases in all cell types, but more so in ventral neural progenitors and astrocytes within hSOs, and in dorsal neural progenitors and choroid plexus cells in hCOs (Fig. 3D, Fig. S3G).

Lastly, we evaluated the cell type-specific expression of *RAMP1, RAMP2* and *RAMP3*, the known receptors for ADM peptide^29,30^. Since the previous literature suggests RAMP2 is the main receptor for ADM, we checked the expression of the *RAMP1-3* receptors in control conditions and found that *RAMP2* is indeed the most expressed receptor. Interestingly, we found this receptor is preferentially expressed by interneurons (hSOs) and glutamatergic neurons (hCOs) when compared to other cell types and its transcription is increased in hypoxia (Fig 3E, Fig S3H).

To validate the findings about the higher expression of RAMP2 in control conditions at a protein level, we performed Western blot analyses for RAMP1 and RAMP2 in hSOs in control conditions and found that, indeed, RAMP2 was significantly more expressed at baseline than RAMP1 by individual cell (two-tailed paired *t*-test, P=0.0004) and by hiPSC line (two-tailed paired *t*-test, P=0.0036) (Fig. 3F,G). To validate the findings about the increase in RAMP2 expression in hypoxia in hSOs, we checked for changes in the expression of RAMP2 at a protein level and identified a significant increase when analyzed by individual cell (two-tailed paired *t*-test, P=0.0027) and by hiPSC line (two-tailed paired *t*-test, P=0.0235) (Fig. 3H,I). These experiments were performed in 3 separate biological replicates of hSOs, each derived from 4 hiPSC lines in one differentiation experiment. Details by hiPSC line are presented in Table S1.

Together, these results provide the first cell type-specific characterization of the expression of ADM and its receptors in the brain in control and hypoxic conditions. Importantly, we identify complementary responses of different cell types, with neurons increasing the expression of the receptors for ADM, while astrocytes, choroid plexus and progenitors increasing the expression of ADM, suggesting a possible evolutionary adaptation mechanism of cells to protect neurons during hypoxia as an environmental stressor.

### Administration of exogenous ADM during exposure to hypoxia rescues the migration deficits of human cortical interneurons

Based on recent reports about the protective role of exogenous ADM administration in various disease processes ^20–22,31,32^, and in the light of our findings described above, we tested the rescue potential of exogenous ADM administration under hypoxic conditions.

We exposed hFAs to hypoxia in the presence of 0.5 μM human ADM (Anaspec, AS-60447) (Fig. 4A) and imaged individual migrating cortical interneurons (Fig. 4B). This dose was chosen based on previous reports from in vitro cultures^33^. We again observed a significant decrease in the number of saltations hypoxic conditions when analyzed by individual cells (paired Wilcoxon test, P<0.0001) and by hiPSC line (two-tailed paired *t*-test, P=0.02), and identified rescue of the number of saltations in hypoxia + ADM conditions when analyzed by individual cells (paired Wilcoxon test, P=0.3) and by hiPSC line (two-tailed paired *t*-test, P=0.9) (Fig. 4C).

**Figure 4.**
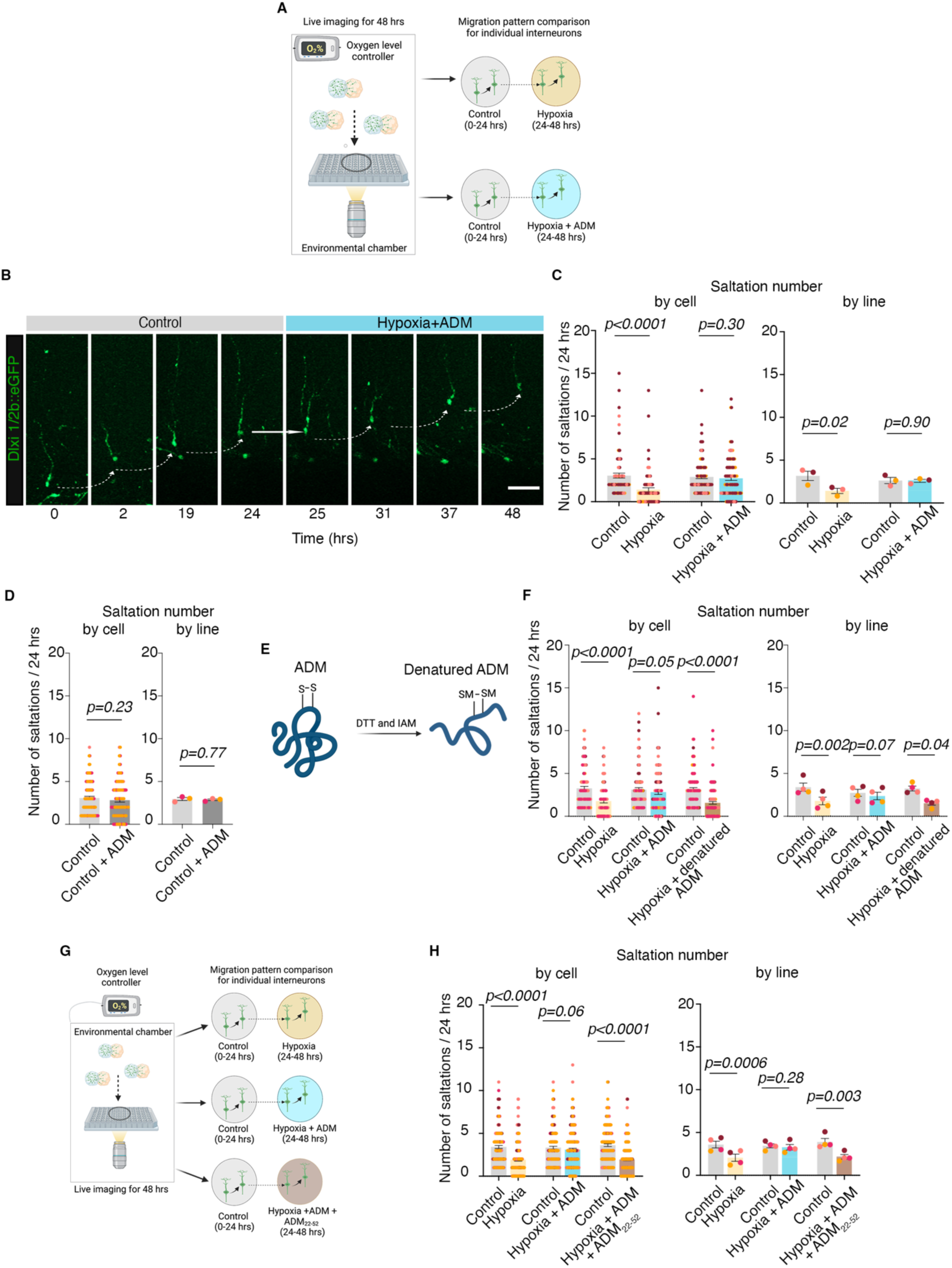
Exogenous administration of ADM peptide rescues the migration defects in hypoxia-exposed human cortical interneurons in an *ex vivo* model using human prenatal cerebral cortex at mid-gestation. **A.** Schematic of experimental design for pharmacological rescue experiments using ADM; 0.5 μM ADM was added to the media at the beginning of hypoxia exposure; **B.** Example of migration pattern for one interneuron in control versus hypoxia + ADM conditions; **C. (Left)** Quantification of saltation number/24 hrs in control versus hypoxia conditions (paired Wilcoxon test, P<0.0001) and in control versus hypoxia + ADM by individual cells (paired Wilcoxon test, P=0.3); **(Right)** Quantification of saltation number/24 hrs in control versus hypoxia conditions (two-tailed paired *t*-test, P=0.02) and in control versus hypoxia + ADM conditions by hiPSC line (two-tailed paired *t*-test, P=0.90); **D. (Left)** Quantification of saltation number/24 hrs in control versus control + ADM conditions by individual cells (paired Wilcoxon test, P=0.23); **(Right)** Quantification of saltation number/24 hrs in control versus control + ADM conditions by hiPSC line (two-tailed paired *t*-test, P=0.77); **E.** Schematic of denaturing procedure for ADM, including reduction and alkylation using dithiothreitol (*DTT*) and iodoacetamide (*IAM*), resulting in carbamidomethylated (*CAM*) cysteines at Cys16 and Cys21; **F.** (**Left)** Quantification by individual cells of saltation number/24 hrs in control versus hypoxia (paired Wilcoxon test, P<0.0001), control versus hypoxia + ADM (paired Wilcoxon test, P=0.05) and control versus hypoxia + denatured ADM (paired Wilcoxon test, P<0.0001); **(Right)** Quantification by hiPSC line of saltation number/24 hrs in control versus hypoxia (two-tailed paired *t*-test, P=0.002), control versus hypoxia + ADM (two-tailed paired *t*-test, P=0.07) and control versus hypoxia + denatured ADM (two-tailed paired *t*-test, P=0.04); **G**. Schematic of experimental design for pharmacological rescue experiments using ADM_22-52_ receptor blocker; **H. (Left)** Quantification by individual cells of saltation number/24 hrs in control versus hypoxia conditions (paired Wilcoxon test, P<0.0001), control versus hypoxia + ADM (paired Wilcoxon test, P=0.06), and control versus hypoxia + ADM + ADM_22-52_ conditions (paired Wilcoxon test, P<0.0001); **(Right)** Quantification of saltation number/24 hrs by hiPSC line in control versus hypoxia conditions (paired t-test test, P=0.0006), control versus hypoxia + ADM (paired t-test, P=0.28), and control versus hypoxia + ADM + ADM_22-52_ conditions (two-tailed paired t-test, P=0.003). Bar charts: mean±s.e.m.

To check whether exogenous ADM peptide changes the migration of cortical interneurons under control conditions, or whether this is hypoxia-specific, we analyzed the number of saltations of cortical interneurons under control conditions for a total of 48 hrs: 24 hrs under control conditions (0-24 hrs), followed by 24 hrs in control + ADM (24-48 hrs). We identified no difference in the total number of saltations/24 hrs when analyzed by individual cells (paired Wilcoxon test, P=0.23), and hiPSC line (two-tailed paired *t*-test, P=0.77) (Fig. 4D), suggesting ADM is not a main mechanism for migration under baseline conditions. These results align with the data about the low baseline expression of ADM (Fig. 2G) in control conditions, and its known increase as an acute phase reactant in hypoxia and inflammation^15–17^.

To verify that the saltation rescue is directly linked to the administration of exogenous ADM, we next performed two types of experiments.

First, we disrupted the disulfide bond between Cys16 and Cys21, which has been shown to be important for ADMs’ biological activity^34^, by denaturing and alkylating it using iodoacetamide (Fig. 4E, Fig. S4A). Next, we repeated the hypoxia experiments in the presence of 0.5 μM human denatured ADM (dADM) and demonstrated that it does not rescue the saltation deficit under hypoxic conditions, when analyzed by individual cells (paired Wilcoxon test, P<0.0001) and by hiPSC line (paired *t*-test test, P=0.04) (Fig. 4F).

Second, we performed pharmacological blocking RAMP2, the primary recognized receptor for ADM in the literature and in our data. For this, we added 10 μM adrenomedullin fragment 22-52 (ADM_22-52_) (Cayman Chemical Company 24892) to the hypoxic media already supplemented with exogenous ADM (Fig. 4G). We validated the saltation deficit in hypoxia and the rescue by exogenous ADM and found that the addition of ADM_22-52_ to the hypoxia + ADM condition was sufficient to prevent the saltation rescue when analyzed in individual cells (paired Wilcoxon test, P<0.0001) and by hiPSC line (paired *t*-test test, P=0.003) (Fig. 4H).

### Migration deficits persist in the acute phases of reoxygenation but administration of exogenous ADM during hypoxia provides partial protection

To investigate whether migration of interneurons is restored upon reoxygenation, we quantified the saltations of individual migrating cortical interneurons for a total of 72 hrs: 24 hrs in control (0-24 hrs), 24 hrs in hypoxia (24-48 hrs), followed by 24 hrs in reoxygenation (48-72 hrs). Similar to our experiments from Fig. 1 and Fig. 4, we observed a ∼53% decrease in saltations in hypoxia compared to control (Friedman test, P<0.0001), and a 65% decrease in the first 24 hrs of reoxygenation compared to control (Friedman test, P<0.0001) when analyzed by individual cells (Fig. S4B). In contrast, the presence of exogenous ADM during hypoxia exposure (not during reoxygenation) fully rescued the migration in hypoxia as previously shown in Fig. 3, and provided partial protection in the first 24 hrs after reoxygenation by decreasing the migration of interneurons by 35% compared to control when analyzed by individual cells (Friedman test, P=0.05) (Fig.S4B) and by hiPSC line (one-way ANOVA, P=0.84) (Fig S4B); again, this is in contrast to the hypoxia conditions without ADM that see a 65% decrease in saltations during reoxygenation.

While not the primary focus of the current study because reoxygenation insult has a different biological mechanism of injury^35,36^ and in-depth analyses would require a different experimental setup, these results reinforce clinical and preclinical data from animal models that reoxygenation injury is important to understand for therapeutic intervention, and that organoid models are able to mimic complex in vivo pathophysiology. Details on the number of cells, experiments and number of hiPSC lines used for each experiment are presented in Table S1.

### *Ex vivo* model using developing human brain tissue confirms the migration deficits and rescue by ADM

To verify our findings in an independent human model, we used *ex vivo* developing human brain tissue at ∼20 PCW and followed a similar experimental design used for hFA (Fig. 5A). Using previously published scRNA-sequencing data from mid-gestation human fetal tissue, we performed subcluster analyses for GABAergic interneurons^16,31^. We identified three subclusters. One small subcluster expressed doublets and was excluded from subsequent analyses (Figg. S5A)^32^. The remaining two sub-clusters were found to express MGE- and CGE-derived markers consisting of NKX2.1, LHX6, CALB2, SST, but minimal CALB1 and *PVALB* similar to the ones in hFAs (Fig. S5B).

**Figure 5.**
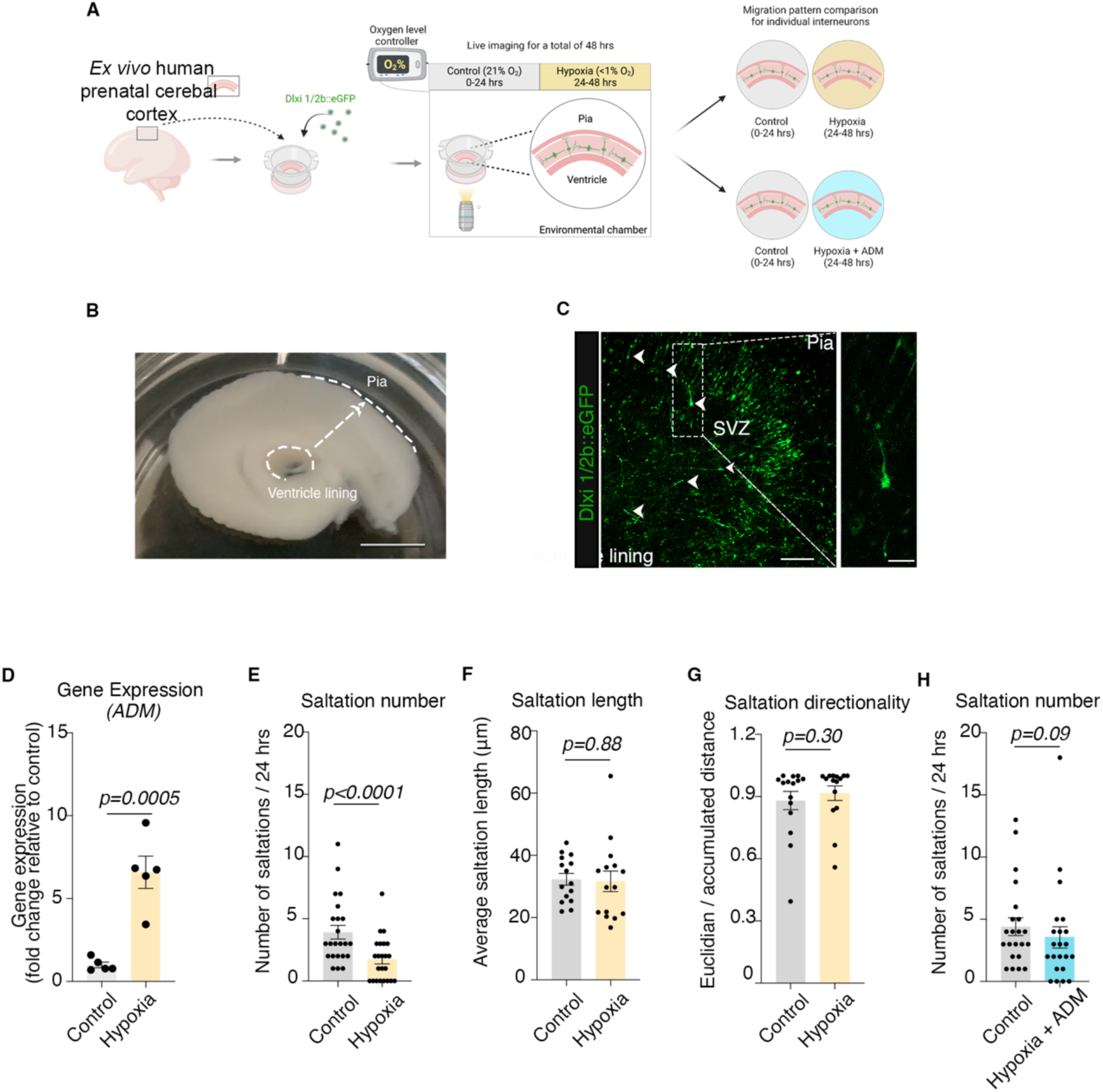
Migration defect and rescue by ADM in an *ex vivo* model using human prenatal cerebral cortex at mid-gestation. **A.** Schematic illustrating the overall experimental design: sections of *ex vivo* human cerebral cortex were collected and initially sectioned at ∼3 mm and subsequently at 400 μm thickness; sections were transferred onto cell culture membrane inserts suspended in culture media; for visualization tissue was transfected with Dlxi1/2b::eGFP lentivirus, and imaged 7-10 days post infection directly on inserts; GFP-tagged *ex vivo* human prenatal cortical interneurons were monitored for 24 hrs in control conditions and 24 hrs in hypoxic conditions in the presence or absence of 0.5 μM ADM; **B.** Example of macroscopic view of a 3mm section of fresh *ex vivo* human prenatal cerebral cortex; scale bar: 1 cm; **C.** Representative image of fluorescently-tagged cortical interneurons in a section of *ex vivo* human prenatal cerebral cortex; **D.** Transcriptional increase of *ADM* gene following 24 hrs of exposure to hypoxia of ex vivo human prenatal cerebral cortex samples (two-tailed unpaired *t*-test, P=0.0005); **E.** Quantification of saltation numbers/24 hrs in control versus hypoxia conditions (paired Wilcoxon test, P<0.0001); **F.** Quantification of average saltation length for control versus hypoxia conditions (paired Wilcoxon test, P=0.88); **G.** Quantification of directionality of migration for control versus hypoxia conditions (paired Wilcoxon test, P=0.3); **H.** Quantification of saltation numbers/24 hrs in control versus hypoxia + ADM conditions by individual cells (paired Wilcoxon test, P=0.09). Bar charts: mean±s.e.m.; scale bars: 1cm, 200 μm and 50 μm.

First, we sectioned human cerebral cortex sections into 400 μm slices (Fig. 5B) and cultured them on live imaging-compatible cell culture inserts^9^ (diameter, 23.1 mm; pore size, 0.4 μm; Falcon, 353090). To visualize migrating cortical interneurons, we fluorescently tagged cells using the same forebrain interneuron cell-type specific lentiviral reporter Dlxi1/2b::eGFP (Fig. 5C)^24^

To assess whether exposure to hypoxia is sufficient to induce a migration deficit in *ex vivo* human cortical interneurons as observed in hFA, we employed the live-imaging setup established for hFA and monitored the migration patterns of the Dlxi1/2b::eGFP^+^-tagged interneurons for 24 hrs under control conditions (37°C, 5% CO_2,_ 21% O_2_). We then transitioned the sections into pre-equilibrated hypoxic media (37°C, 5% CO_2_, 94.5% N_2_, <1% O_2_) and imaged the same cells for another 24 hrs. Similar to hypoxic hSOs, we observed a significant transcriptional increase in *ADM* gene by qPCR (two-tailed unpaired *t-*test, P=0.0005) (Fig. 5D). Following exposure to hypoxia for 24 hrs, we identified ∼55 % decrease in the number of saltations (paired Wilcoxon test, P<0.0001) (Fig. 5E), but no changes in the average saltation length (paired Wilcoxon test, P=0.88) (Fig. 5F) nor in directionality (Wilcoxon test, P=0.3) (Fig. 5G).

Lastly, we found addition of 0.5 μM ADM peptide during the hypoxia exposure was sufficient to rescue the saltation numbers (paired Wilcoxon test, P=0.09) (Fig. 5H).

### Proposed molecular mechanism that contributes to the rescue by exogenous ADM administration

First, we performed EIA analyses focused on cAMP/PKA, AKT/pAKT and ERK/pERK pathways, which have been previously shown to be modulated by ADM^37^ (Fig. 6A). We found that the concentration of cAMP was not affected by hypoxia but was significantly increased in the hypoxia + ADM conditions, both when analyzed by individual samples (one-way ANOVA test, P=0.0007), and by hiPSC line (one-way ANOVA test, P=0.002) (Fig. 6B). These results were complemented by increased PKA activity in hypoxia + ADM conditions when analyzed by individual samples (one-way ANOVA test, P<0.0001), and by hiPSC line (one-way ANOVA, P<0.0001) (Fig. 6C). In contrast, the ratio of pAKT/AKT was significantly decreased in hypoxia samples when analyzed by individual samples (one-way ANOVA, P<0.0001) and by hiPSC line (Kruskal-Wallis test, P=0.01), but was not rescued in hypoxia + ADM (Fig. 6D). These findings were similar for the ratio of pERK/ERK in individual samples (one-way ANOVA test, P=0.001) and by hiPSC line (one-way ANOVA, P=0.02) (Fig. 6E). Lastly, we quantified the ratio of pCREB/CREB which is downstream of all these pathways. We found that hypoxia induced a significant decrease in the ratio of pCREB/CREB when compared to control when analyzed by individual samples (one-way ANOVA test, P=0.003) and by hiPSC line (P=0.048). However, this ratio was restored in hypoxia + ADM conditions when analyzed by individual samples (one-way ANOVA test, P=0.27) and by hiPSC lines (one-way ANOVA, P=0.6) (Fig. 6F). These results suggest the cAMP/PKA/pCREB pathway is modulated by exogenous ADM administration in hypoxia in hSOs.

**Figure 6.**
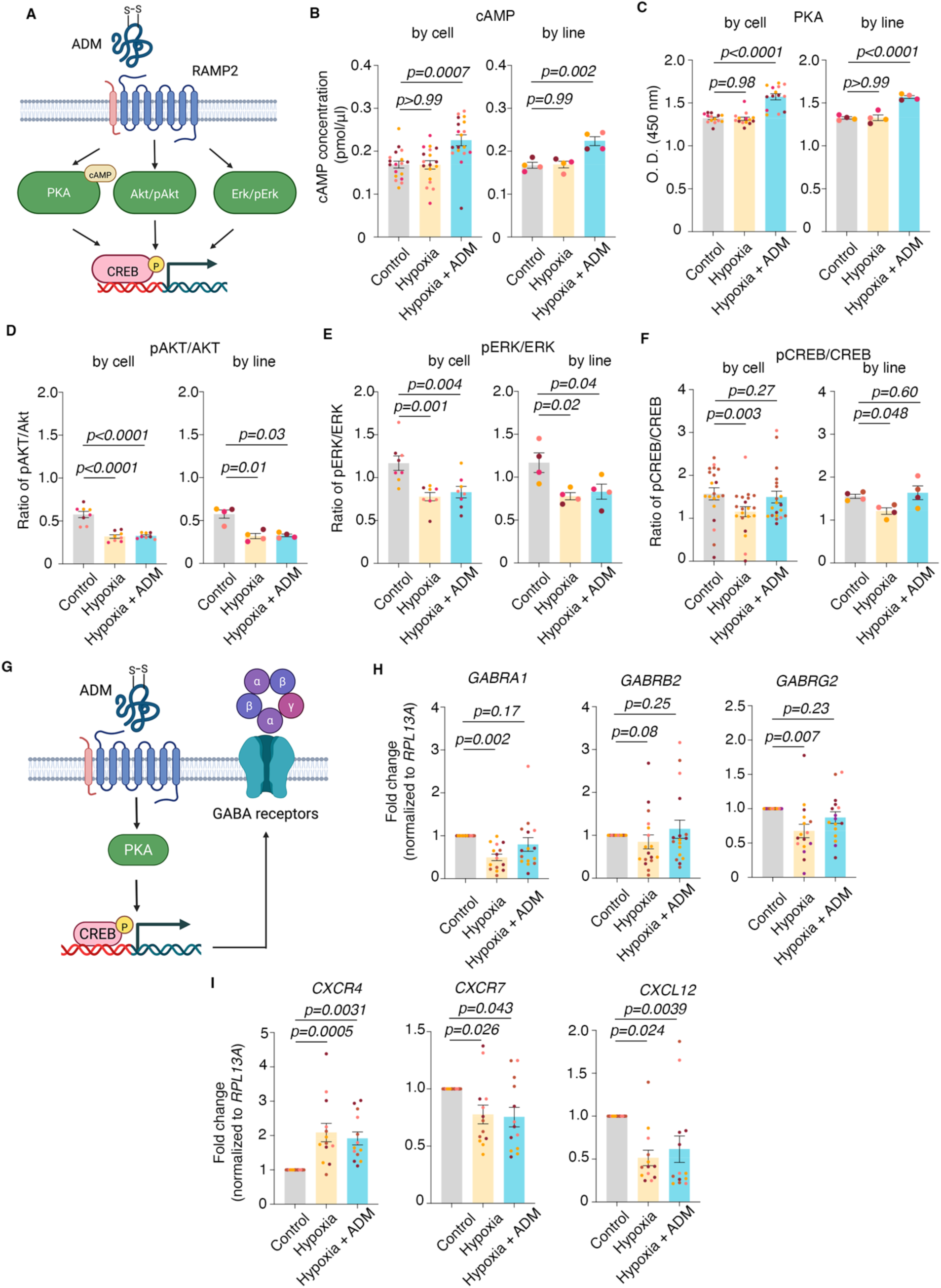
Molecular mechanism of rescue by ADM. **A.** Schematic of the previously reported main molecular pathways modulated by ADM; **B.(Left)** Quantification of cAMP concentration (pmol/μl) by individual hSOs in control versus hypoxia (one-way ANOVA, P>0.99) and, control versus hypoxia + ADM (one-way ANOVA, P=0.0007), **(Right)** Quantification of cAMP concentration (pmol/μl) by hiPSC line in control versus hypoxia (one-way ANOVA, P=0.99) and, control versus hypoxia + ADM (one-way ANOVA, P=0.002); **C. (Left)** Quantification of PKA activity (O.D. 450nm) by individual hSOs in control versus hypoxia (one-way ANOVA test, P=0.98) and control versus hypoxia + ADM (one-way ANOVA, P<0.0001); **(Right)** Quantification of PKA activity (O.D. 450nm) by hiPSC line in control versus hypoxia (one-way ANOVA, P>0.99) and control versus hypoxia + ADM (one-way ANOVA, P<0.0001); **D. (Left)** Quantification of pAKT/AKT by individual hSOs in control versus hypoxia (one-way ANOVA, P<0.0001) and control versus hypoxia + ADM (one-way ANOVA, P<0.0001); **(Right)** Quantification of pAKT/AKT by hiPSC line in control versus hypoxia (Kruskal-Wallis test, P=0.01) and control versus hypoxia + ADM (Kruskal-Wallis test, P=0.03); **E. (Left)** Quantification of pERK/ERK by individual hSOs in control versus hypoxia (one-way ANOVA test, P=0.001) and control versus hypoxia + ADM (one-way ANOVA test, P=0.004); **(Right)** Quantification of pERK/ERK by hiPSC line in control versus hypoxia (one-way ANOVA test, P=0.02) and control versus hypoxia + ADM (one-way ANOVA test, P=0.04); **F. (Left)** Quantification of pCREB/CREB by individual hSOs in control versus hypoxia (one-way ANOVA test, P=0.003) and control versus hypoxia + ADM (one-way ANOVA test, P=0.27); **(Right)** Quantification of pCREB/CREB by hiPSC line in control versus hypoxia (one-way ANOVA test, P=0.048) and control versus hypoxia + ADM (one-way ANOVA test, P=0.6); **G.** Schematic of the proposed molecular pathway activation by exogenous ADM in hSOs, including the most common pentameric structure of the GABA_A_ receptor; **H. (Left)** Quantification (by q-PCR) of GABRA1 in hSOs samples in control versus hypoxia conditions (one-way ANOVA test, P=0.002) and control versus hypoxia + ADM (one-way ANOVA test, P=0.17); **(Center)** Quantification (by q-PCR) of GABRB2 in hSOs samples in control versus hypoxia (one-way ANOVA test, P=0.08) and control versus hypoxia + ADM (one-way ANOVA test, P=0.25); **(Right)** Quantification (by q-PCR) of GABRG2 in hSO samples in control versus hypoxia conditions (one-way ANOVA test, P=0.007) and control versus hypoxia + ADM (one-way ANOVA test, P=0.23); **I. (Left)** Quantification (by q-PCR) of CXCR4 in hSOs samples in control versus hypoxia conditions (one-way ANOVA test, P=0.0005) and control versus hypoxia + ADM (one-way ANOVA test, P=0.0031); **(Center)** Quantification (by q-PCR) of CXCR7 in hSOs samples in control versus hypoxia conditions (one-way ANOVA test, P=0.026) and control versus hypoxia + ADM (one-way ANOVA test, P=0.043); **(Right)** Quantification (by q-PCR) of CXCL12 in hSOs samples in control versus hypoxia conditions (one-way ANOVA test, P=0.024) and control versus hypoxia + ADM (one-way ANOVA test, P=0.0039). Bar charts: mean±s.e.m.; Different dot colors represent individual hiPSC lines.

**Figure 7.**
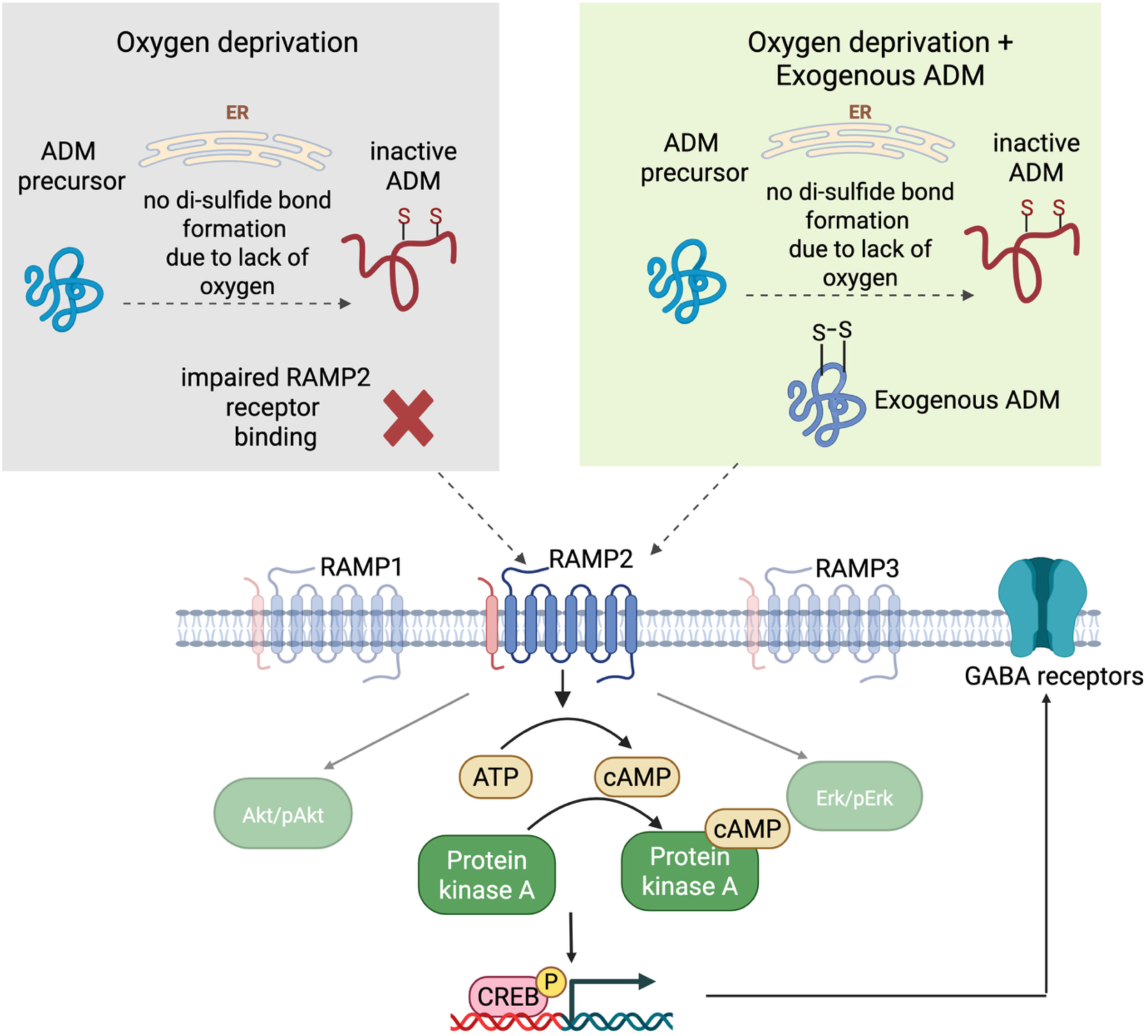
Schematic of the overall proposed mechanism of interneuron migration defect rescue by ADM upon hypoxia exposure. Based on our findings and existing data from literature, we propose endogenously produced ADM has decreased biological activity by impaired ability to form the necessary di-sulfide bond in the absence of oxygen in hypoxia. However, exogenous ADM does have biological activity as the di-sulfide bond is present, and thus it binds efficiently to its receptors, especially RAMP2. This binding initiates an activation of the cAMP/PKA/pCREB pathway, which in turn restore the expression of GABA_A_ receptors and rescue the migration.

Previous reports demonstrated that increased ratio of pCREB/CREB is directly involved in increasing the expression of GABA_A_ receptors (Fig. 6G)^38,39^, which in turn, are well-documented to facilitate interneuron migration^40^. To check whether the expression of GABA_A_ receptors is affected under hypoxic conditions and rescued by ADM, we quantified the expression of several GABA_A_ receptor subunits in the presence or absence of ADM during hypoxia exposure. We specifically focused on the most common pentameric combination, α1β2γ2, encoded by *GABRA1*, *GABRB2* and *GABRG2*^41^ (Fig. 6G). We found a significant decrease in expression in hypoxia (one-way ANOVA test; control vs hypoxia: P=0.002) and rescue by ADM for *GABRA1* (one-way ANOVA test; P=0.17); no significant change in expression in *GABRB2* (Kruskal-Wallis test; control versus hypoxia: P=0.08; control versus hypoxia + ADM: P=0.25); a significant decrease in expression in hypoxia (one-way ANOVA test; control versus hypoxia: P=0.007) and rescue by ADM for *GABRG2* (one-way ANOVA test; P=0.23) (Fig. 6H). Moreover, we analyzed other receptor subunits and found mild but still significant decrease in hypoxia and rescue by ADM for receptors *GABRB3* (Kruskal-Wallis test; control versus hypoxia: P=0.02; control versus hypoxia + ADM: P=0.05) and *GABRG3* (one-way ANOVA test; control versus hypoxia: P=0.01; control versus hypoxia + ADM: P=0.1); no change in expression under hypoxic conditions for receptors *GABRA2* (one-way ANOVA; control versus hypoxia: P=0.8; control versus hypoxia + ADM: P=0.75) (Fig. S6A).

Separately, we investigated the expression of *CXCR4*, *CXCR7* and *CXCL12*, as some of the best-known chemokines/chemokine receptors involved in interneuron migration^42,43^. Interestingly, we found a significant increase of *CXCR4* in hypoxia but no rescue by ADM (one-way ANOVA test; control versus hypoxia: P=0.0005; control versus hypoxia +ADM: P=0.0031), a significant decrease of *CXCR7* in hypoxia but no rescue by ADM (one-way ANOVA test; control versus hypoxia: P=0.026; control versus hypoxia + ADM: P=0.043), and a significant decrease of *CXCL12* in hypoxia but no rescue by ADM (one-way ANOVA test; control versus hypoxia: P=0.024; control versus hypoxia + ADM: P=0.0039) (Fig.6I).

Overall, these results suggest that rescue of GABA_A_ receptor expression is sufficient to restore migration patterns in hypoxic cortical interneurons, even in the absence of rescue of cytokine receptors.

## DISCUSSION

Delayed or reduced migration of inhibitory interneurons into the developing cortex can have profound consequences for neural circuit formation and function. Proper timing and integration of GABAergic interneurons are essential for establishing the excitatory–inhibitory (E/I) balance that regulates cortical oscillations, synaptic refinement, and neuronal synchrony^44,45^.Disruption of interneuron migration can lead to abnormal circuit assembly and hyperexcitability, ultimately impairing synaptic plasticity and cortical information processing^26,46^. Functionally, such alterations have been associated with several neurodevelopmental and neuropsychiatric conditions, including epilepsy, autism spectrum disorder (ASD), and schizophrenia^47,48^.

Disruptions in cortical interneurons migration from the ganglionic eminences toward the cortex have been long suggested as a contributing factor to the increased risk for neuropsychiatric diseases in hypoxic brain injury of prematurity, because this critical developmental process occurs predominantly during the latter half of pregnancy and is likely to be affected by preterm birth. However the migration patterns under hypoxic conditions have proved difficult to investigate in animal models, and no data exist for human cortical interneurons.

Given the accumulating evidence demonstrates the power of human cellular models for the study of neurodevelopmental diseases of genetic or environmental etiology, including tuberous sclerosis, Timothy syndrome, 22q11.2 deletion syndrome, ZikV exposure, preterm birth, etc.^39,49–52^, in this study, we established the first live imaging microscopy platform for the direct investigation of the migration patterns on human cortical interneurons in control and hypoxic conditions, thus overcoming a long-term challenge in the field. The high technical and scientific relevance of this novel platform for disease investigation is further amplified by the concomitant use of two human models consisting of hFAs derived from hiPSCs and *ex vivo* developing human cortical tissue, an approach that helps mitigate potential confounding factors associated with possible interspecies differences in how brain cells respond to hypoxia and other environmental stressors.

Using these human cellular platforms, we directly demonstrate a substantial saltation deficit of human cortical interneurons exposed to mild hypoxia. Importantly, we provide the first characterization of molecular changes in interneurons in hypoxia, identify exogenous adrenomedullin (ADM) peptide supplementation as an effective pharmacological rescue for this phenotype and bring evidence that this is, at least partly, mediated by the cAMP/PKA/pCREB pathway and rescue of the expression of specific subunits that form the α1β2γ2 pentameric GABA_A_ receptor. Moreover, in this study we provide preliminary evidence for the ongoing migration deficits in the first 24 hrs after reoxygenation, suggesting a longer-term effect of a hypoxic event, and the possible protective effect of exogenous ADM administration. Lastly, we provide evidence that the endogenous production of ADM is not sufficient for a phenotypic rescue, and we suggest this is due to the decreased biological activity of endogenously produced ADM, sue to inability to efficiently form the disulfide bond in a hypoxic environment. Since exogenous ADM administration is already showing promise in clinical trials for inflammatory bowel disease^19^, this new knowledge could serve as a starting point for the investigation of ADM as a possible future target for therapeutic interventions for hypoxic brain injuries. Lastly, the novel and interesting findings about the cell type-specific expression of ADM and its receptors on brain cells add important biologic knowledge and will inform future studies focused on the role of ADM in the brain.

Our manuscript has several limitations. First, while hFAs and *ex vivo* developing human cortical tissue are the “best next thing” after in vivo human brain, they do remain a model, like all other preclinical models. The migration of interneurons observed in hSO–hCO assembloids recapitulates several key aspects of *in vivo* cortical development, including directional migration from subpallial to pallial regions, responsiveness to chemotactic gradients, and integration into cortical-like networks. However, this model remains a simplified representation of the developing human brain. Several environmental components are absent, including vascularization, immune and glial interactions, long-range axonal inputs, and extracellular matrix complexity, all of which shape interneuron migration and maturation *in vivo*. Future studies should focus on the translational potential of ADM for brain injury of prematurity using large *in vivo* animal models (e.g. piglet, lambs, non-human primates) which have more similar brain developmental trajectories to humans, allow reliable modeling of preterm birth and multiple complex analyses (e.g. blood draws, neurological monitoring etc), which are not feasible in organoids. Second, while we show ongoing injury after reoxygenation, we do not provide in depth phenotypic cellular and molecular characterization of this injury. Since reoxygenation injury is an important, yet separate entity in the complex pathophysiology of hypoxic brain injury, future studies should adapt our current experimental design to focus on this question, which will be highly relevant for the future development of ADM and other compounds as therapeutics. Third, this study was specifically focused on studying the migration of cortical interneurons, however, other developmental processes are likely affected, including the proliferation of ventral forebrain progenitors and the functional integration of mature neurons within the neuronal cortical networks. These should be addressed in future studies. Lastly, here we focused on ADM, while excluding the detailed investigation of other molecular pathways and genes that were differentially expressed, including the inflammatory pathway, which is highly relevant for the pathophysiology of hypoxic brain injury of prematurity.

## METHODS

### Culture of hiPSCs

The hiPSC lines used in this study were validated using standardized methods we previously described^9,10,21,41^. Cultures were regularly tested for, and maintained Mycoplasma free. SNP-array experiments were performed for genomic integrity. The differentiation experiments were performed using 4 control hiPSC lines derived from fibroblasts harvested from 4 healthy individuals (two XX and two XY). Informed consent for fibroblast collection was obtained from all individuals and this process was approved by the Stanford IRB Panel.

### Generation of region-specific organoids

Brain region-specific human cortical organoids (hCO) and human subpallium organoids (hSO) were differentiated using feeder-free grown hiPSCs using validated protocols we previously reported and contributed to developing^9,10,20,21,41–45^. Briefly, hiPSCs were maintained in six-well plates coated with recombinant human vitronectin (VTN-N, Thermo Fisher Scientific, A14700) in Essential 8 medium (Thermo Fisher Scientific, A1517001). For differentiation, hiPSC colonies were lifted from the plates using Accutase (Innovate Cell Technologies, AT-104). Approximately 3 million cells were transferred to each well in AggreWell 800 (STEMCELL Technologies, 34815) and centrifuged at 100g for 3 minutes in hiPSC medium supplemented with ROCK inhibitor Y-27632 (Selleck Chemicals, S1049). After 24 hrs, the newly formed organoids were transferred into ultra-low-attachment plastic dishes (Corning, 430293) in Essential 6 medium (Thermo Fisher Scientific, A1516401) supplemented with dorsomorphin (2.5 μM, SigmaAldrich, P5499) and SB-431542 (10 μM, Tocris, 1614) and additional XAV-939 (hCO 0.5 μM, hSO: 2.5 μM Tocris, 3748). This medium was replaced daily for the first five days. On the sixth day in suspension, organoids were transferred to neural medium containing Neurobasal A (Thermo Fisher Scientific, 10888022), B-27 supplement without vitamin A (Thermo Fisher Scientific,12587010), GlutaMax (2 mM, Thermo Fisher Scientific, 35050061) and penicillin and streptomycin (100 U ml^−1^, Thermo Fisher Scientific, 15140122). For hCO the neural medium was supplemented with the growth factors EGF (20 ng ml^−1^, R&D Systems, 236-EG) and FGF2 (20 ng ml^−1^, R&D Systems, 233-FB) every day until day 15, then every-other day until day 23. For hSO, we included continued supplementation with XAV-939 (2.5 μM) on days 6-23, and SHH pathway agonist SAG (smoothened agonist; 100 nM; Thermo Fisher Scientific, 566660) on days 12-23. Between days 25-43, the neural medium was supplemented with neurotrophic factors BDNF (20 ng ml^−1^, Peprotech, 450-02) and NT3 (20 ng ml^−1^, Peprotech, 450-03), to promote differentiation of the neural progenitors into neurons in both hCO and hSO; media change was performed every other day. After day 43, hCO and hSO were maintained in neural media.

### Viral labeling of hSO and generation of human forebrain assembloids (hFA)

The viral infection of the hSO for visualization of migrating cortical interneurons was performed as previously described^9,37^. Briefly, at approximately 45-55 days in culture, hSO were transferred to a 24 well plate (Corning, 3474) containing 500μl on neural medium with 10μl of virus (Lenti-Dlxi1/2b::eGFP; construct reported and applied in reference) ^22^. After 24 hours, 500μl of neural medium was added. On day 3, all media was removed and replaced with 1ml of neurobasal medium. The next day, hSO were transferred into fresh neural media in ultra-low attachment plates. After 5-7 days, hCO and virally infected hSO were used to generate hFA. To do this, one hCO and one hSO were transferred together into a 1.5 ml microcentrifuge Eppendorf tube and maintained in direct contact for three days. More than 95% of hCO and hSO fused. Subsequently, hFA were carefully transferred into 24 well ultra-low attachment plates (Corning, 3474) and media changes were performed very gently every two to three days. Lentivirus construct (lenti-Dlxi1/2b::eGFP) was received as a gift from J. L. Rubenstein, and virus was generated by transfecting HEK293T cells with PEI Max (Polysciences, 24765); after 48 hrs the media was collected and ultracentrifuged at 17,000 rpm at 6°C for 1 hr.

### Primary human prenatal cortex processing and viral labeling

*Ex vivo* developing human cortical tissue at 20 PCW was obtained under a protocol approved by the Research Compliance Office at Stanford University. The tissue was processed within 4 hours after collection using a previously described protocol^9,37,41^. In brief, cortical tissue was embedded in 4% low melting-point agarose in PBS and cut using a Leica VT1200 Vibratome at 400 μm. The sections were transferred onto cell culture membrane inserts (diameter, 23.1 mm; pore size, 0.4 μm; Falcon, 353090) and incubated in culture media (66% BME, 25% Hanks, 5% FBS, 1% N-2, 1% penicillin, streptomycin and glutamine; all from Invitrogen) and 0.66% D-(+)-Glucose (Sigma) at 5% CO_2_, 37 °C with the Dlxi1/2b::eGFP lentivirus for 24 hrs. Sections were then transferred to fresh cell culture media and half media changes were performed every other day. After ∼5 days in culture, Dlxi1/2b::eGFP+ cells could be detected and imaging was performed within 7-10 days in culture.

### Live imaging and analyses of migration patterns of Dlxi1/2b::eGFP-tagged cortical interneurons

For imaging, hFA were transferred intact into a 96–well plate (glass-bottom, Cellvis, Catalog #: P96-0N) in 300 μl of neural media. Imaging was performed under environmentally controlled conditions (37°C, 5% CO_2,_ 21% O_2_) using a confocal microscope with a motorized stage (Zeiss LSM 980) and equipped with a cage incubator for temperature, gas (including O_2_ levels) and humidity control (Okolab Bold Line). During each recording session, we imaged up to 12 hFA at 10X magnification, at a depth of 350-400μm, and at a rate of 20 min/frame. The imaging field was focused on the hCO side of the fusion.

Samples were imaged for 24 hrs under control conditions (37°C, 5% CO_2_, 21% O_2_). To induce acute hypoxia, media was carefully removed and replaced with pre-equilibrated hypoxic media. All procedures were performed in the controlled hypoxic environment of the HypoxyLab (Oxford Optronix). The field of view was readjusted to capture the previous region of interest, and the same cells were imaged for an additional 24 hrs while maintaining a hypoxic environment (37°C, 5% CO_2_, <1% O_2_). For pharmacological rescue experiments, the pre-equilibrated hypoxic media containing the peptide Adrenomedullin (ADM) (0.5 μM, Anaspec, AS-60447) +/− 10 uM ADM_22-52_ (the RAMP2 receptor blocker (Cayman Chemical Company, cat. no. 24892)), were added to the hFA cell culture media during hypoxia exposure.

The imaging of migration of *ex vivo* human developing cortical interneurons was done with the same settings as described above, with the exception of image depth being 150-200μm. Slices were kept on the cell culture inserts during imaging.

Post-acquisition analyses of cell mobility were performed using ImageJ (v:2.00-rc-69/1.52n). Individual cells were identified and followed both in control and hypoxic conditions. Analyses were performed on all cells that had at least one saltation within the first 24 hrs of imaging (control conditions) and remained visible within the field of view until the end of the imaging period (up to 72 hrs). Paired analyses of number of saltations and saltation length were performed as described in the main text. To estimate the length of individual saltations, we manually tracked and identified the swelling of the soma of Dlxi1/2b::eGFP tagged cortical interneurons before and after saltation and measured the distance (in μm) from the initial position to the new position. The Z-stack Alignment present in Zen 3.1 Blue Edition (Zeiss) was used to correct for minor drifts during imaging. To assess changes in directionality of the movement, we extracted the x and y coordinates of each cell per frame and time using the Manual Tracking plugin (ImageJ). The Chemotaxis & Migration Tool (Ibidi) was then used to calculate the Accumulated (A) and Euclidean (E) distances traveled per cell over time. Data is presented by calculating the E/A ratio. Only cells which had at least one saltation within each of the conditions were included in this analysis.

### Hypoxia treatment of organoids and primary tissues for *in vitro* assays

Organoids or developing human cortical tissue used for in vitro hypoxia experiments were transferred to culture medium pre-equilibrated in a hypoxic C-chamber (Biospherix) or HypoxyLab (Oxford). Hypoxic C-chamber was connected to a premixed gas tank containing 5% CO_2_ and balancing N_2_. Oxygen level in the C-chamber was controlled and monitored using a Proox 110 Compact Oxygen Controller to reach <1% O_2_. HypoxyLab was connected to O_2_, N_2_, and CO_2_ gas tanks and built-in gas sensors control each gas level in real-time to achieve 30 mmHg with 5% CO_2_. After 24 hrs treatment by hypoxia, organoids or primary tissues were collected immediately for assays.

### Oxygen measurements in control and hypoxic cell culture media

The O_2_ tension of the media was measured using the oxygen optical microsensor OXB50 (50 μm, PyroScience) attached to a fiber-optic multi-analyte meter (FireStingO_2_, PyroScience), as previously described^38,45^.

### RNA-sequencing of hSO

At 46 days in culture, 3 hSO from 4 individual hiPSC lines were exposed to hypoxic conditions for 12 and 24 hrs using a hypoxia C-chamber as described above. For acute induction of hypoxia, the media was previously equilibrated overnight at <1% O_2_, 5% CO_2_, 37 °C with a resulting PO_2_ of 25-30mmHg. After 12 hrs and 24 hrs of hypoxic exposure, hSO were immediately collected and snap-frozen in dry ice for analyses. Control hSO maintained in regular incubator conditions were collected from each hiPSC line at each experimental timepoint. Additionally, extra hSO exposed to 24 hrs of hypoxia were transferred back to baseline conditions and reoxygenated for 72 hrs. At this time point, samples were collected, matched with control hSO and snap-frozen on dry ice. All samples were kept at −80°C until they were processed for RNA-sequencing.

### RNA sequencing analyses

mRNA was isolated using the RNeasy Mini kit (Qiagen) and sequence libraries were prepared by Admera Health (www.admerahealth.com) using the KAPA Hyper Prep Stranded RNA-Seq with RiboErase. Sequencing was performed on the Illumina HiSeq 400 system with paired-end 150 base pair reads. Samples averaged 40 million reads, each representing 20x genome coverage. Trimmed RNA-Seq reads were aligned to the Ensembl GRCh38 human genome reference using STAR v2.7.3 in gene annotation mode^28^. Alignment and RNA-Seq quality control metrics were generated with Picard (v 2.21.1). Principal component analysis (PCA) of the quality control matrices did not indicate any sample outliers.

Gene-level counts were filtered to remove lowly expressed genes which did not have at least 1 count per million in at least 4 out of the total 24 samples; this resulted in 17,777 remaining genes. We noticed a slight GC content bias across samples, so gene counts were further normalized by GC content, gene length, and sequencing depth using the CQN R package v1.36.0^40^. The resulting estimation parameters were provided to the negative binomial generalized linear model while performing the differential expression (DEG) analysis. This was carried out using DESeq2 v1.30.1 to identify significant genes between hypoxic conditions at each timepoint while also accounting for unwanted variation due to cell line^42^. The resulting P values were corrected for multiple comparisons using the Benjamini-Hochberg method. Genes were considered significant at FDR ≤ 0.05 and absolute fold change > 1.5 (log2FC of 0.6). In total, we identified 734 DEX genes as a consequence of the hypoxic conditions at 12 hrs, 985 genes at 24 hrs, and only 21 genes at 72 hrs after reoxygenation (Fig. 2b, 2c, Fig. S2b, S3). We generated the dendrogram in Figure 2b using expression data from all 24 samples, narrowed to the union of the genes differentially expressed between hypoxic and control conditions at 12 and 24 hrs or 72 hrs after reoxygenation (*n* = 1,473). Hierarchical clustering, using complete linkage of Euclidean distance, determined the order of the dendrogram. Z-score normalized expression values are depicted on a continuous scale from lower values (purple) to higher values (orange). The notable reduction of DEG at 72 hrs after reoxygenation reflects what we visually see in the PCA of the normalized gene expression, in that the control and hypoxia conditions cluster together at 72 hrs post reoxygenation (not shown). Indeed, hierarchical clustering using the Ward’s criterion on the first three principal components separates hypoxic samples at 12 and 24 hrs from the control 12 and 24 hrs samples as well as the reoxygenated samples (Fig. S2b).

To gain insight about putatively dysregulated biological pathways, we conducted gene set enrichment analysis that was computed using the fgsea R package v1.18.0 where all genes were ranked by their absolute log_2_ fold change^43^. We evaluated the Molecular Signatures Database “Hallmark” gene sets and supplemented the WU_cell migration set based on its presence in a preliminary gene ontology enrichment. The top 10 significantly enriched gene sets (padj < 0.05) that occur across all timepoints are shown in Figure 2d. Interested in discovering which genes had the largest change in response between 12 hrs and 24 hrs, we identified genes significant at both timepoints and ranked them by absolute difference in fold change. The top few genes increasing and decreasing in expression are shown in Fig. 2e.

### Single cell gene expression data collection and analyses

For single cell gene expression analyses, human forebrain assembloids (hFA) were infected with Lenti Dlx1/2b::eGFP as described above. Samples were incubated either under a normoxic environment (37°C, 5% CO_2_, 21% O_2_), or under the controlled hypoxic environment of the HypoxyLab (Oxford Optronix). Following the environmental exposure, hFA were dissected under a microscope to separate the hFA into hSO and hCO. To analyze only interneurons migrated within the hCO and not the ones still at the edges of the fusion, we intentionally performed the separation of hFA more toward the hCO side. For single cell gene expression analysis, organoids were dissociated as previously described^16,47^. Dissociated cells were resuspended in ice-cold PBS containing 0.02% BSA and loaded onto a Chromium Single cell 3′ chip (with an estimated recovery of 10,000 cells per channel) to generate gel beads in emulsion (GEMs). scRNA-seq libraries were prepared with the Chromium Single cell 3′ GEM, Library & Gel Bead Kit v2 (10x Genomics, PN: 120237). Libraries from different samples were pooled and sequenced on a NovaSeq S4 (Illumina) using 150 × 2 chemistry. Sample demultiplexing, barcode processing and unique molecular identifiers counting was performed using the Cell Ranger software suite (v6.1.2 with default settings excluding introns). Reads were then aligned to the human reference genome (GRCh38), filtered, and counted. Expression matrices for the samples were processed using the R (v4.1.2) package Seurat (v4.2). We excluded cells that express less than 2500 genes as well as those with a proportion of mitochondrial reads higher than 15%. Gene expression was then normalized using a global-scaling normalization method (normalization.method=‘LogNormalize’, scale.factor = 10,000), and the 2,000 most variable genes were then selected (selection.method=‘versust’). *Harmony* package^45^ was used to integrate control and hypoxia samples for downstream analysis. The top 7 and 6 principal components for hCO and hSO samples respectively were utilized to perform a UMAP projection, implemented with the *RunUMAP* function with default settings. The clusters were generated with resolutions of 0.1 and 0.3 respectively for hCO and hSO samples using the *FindNeighbors* and *FindClusters* functions. We identified clusters based on differential expression of known markers^44^ identified using the FindAllMarkers function with default settings. In some cases, using expressions of known markers, we grouped together clusters that were originally separate based on the Seurat clustering. For visualization purposes, the color palette of UMAP plots for hSO and hCO was adjusted using “magic wand” in non-contiguous mode for selection of all cells from respective clusters, for subsequent synchronization of the color for the given cell cluster between brain regions (Adobe Photoshop CS 2018). Original plots are available upon request. Markers used to verify the hypoxic response were based on *PDK1* and *PFKP.* Proportions of cells expressing *ADM* (Expression Level>0) out of all cells per cluster were calculated using^38^.

### Cryopreservation and cryosection of hSO

Whole hSOs that were infected with Dlxi1/2b::eGFP^+^ were fixed with 4% paraformaldehyde overnight. They were then washed with PBS and transferred to 30% sucrose for up to 72 hrs. The samples were transferred into an embedding medium block (Tissue-Tek OCT Compound 4583, Sakura Finetek), snap frozen and sectioned at 10- μm thick microns using a cryostat (Leica).

### Immunocytochemistry and quantification of cleaved-caspase 3

The cryosections of hSO were washed with PBS three times for 5 mins each to remove excess OCT and then blocked for 1hr in PBS + 0.5% triton X-100 + 5% BSA + 0.5% DMSO at room temperature. The slides were then incubated with cleaved-caspase 3 (anti-rabbit, 1:100, Cell Signaling Technology, 9661T) overnight at 4°C. After incubation, the slides were washed three times with PBS + 0.5% triton X-100 for 5 mins each. Secondary antibody incubation was then performed with Alexa Fluor 594 (anti-rabbit, 1:1000, Invitrogen, Cat # A-21207), anti-GFP (1:1000, GeneTex, GTX13970) for 1 hr at room temperature. After, slides were washed two times with PBS for 5 mins each and then stained with Hoescht 33258 (1:10,000, Life Technologies) for 5 mins at room temperature. Cryosections were mounted on glass coverslips using aquamount (Thermo Fisher Scientific) and were imaged using (Zeiss Axio Observer). Images were visualized and the number of cleaved-caspase 3 (c-CAS3) and Dlxi1/2b::eGFP^+^ positive cells were quantified using Fiji ImageJ Version 1.54p).

### Flow cytometry analyses for Annexin V

To assess the levels of apoptotic Dlxi1/2b::mScarlet cells, FACS was conducted on NovoCyte Penteon instrument using NovoExpress 1.5.6 software (Stanford FACS Facility). To label inhibitory interneurons in hSOs at later stages of development, hiPSC lines were engineered to express mScarlet under the dlx1/2 promoter. Briefly, LV-Dlxi1/2b-mScarlet lentiviral particles were produced according to the protocol described in this manuscript. Control iPSC lines were plated sparsely and infected with the lentivirus at 1:300 dilution in E8 medium for 24 hours, following which the media was changed to E8 medium without lentivirus. The cells were dissociated using Accutase (Innovate Cell Technologies, AT-104) and plated sparsely in 6-well plates for Puromycin (Sigma-Aldrich, P8833) selection of single cell clones with successful genomic integration of the LV-Dlxi1/2b-mScarlet construct. Selected clones were used to make hSO organoids according to the previously described protocol and used for further experiment. On the day of analysis, 3 spheres per line were placed in an Eppendorf tube and dissociated in 300μL of Accutase at 37°C for 30mins. After incubation, 1200μL of neurobasal medium was added to each tube to inactivate the Accutase. The cells were then centrifuged at 200g for 4 mins and resuspended in DPBS (Thermo Fisher Scientific, 14190144) and strained using a 40μM cell strainer (Biologix Research Company, 15-1040). Using the Annexin V kit (Invitrogen, V13246) the cells were stained with Annexin V, Alexa Fluor-647 (Invitrogen, A23204) in RT for 15 mins. The analysis of Annexin V signals in Dlxi1/2b::mScarlet cells was done using FlowJo (v.10.8.1).

### Real time quantitative PCR (qPCR)

mRNA was isolated using the RNeasy Mini Kit and RNase-Free DNase set (Qiagen, 74136), and template cDNA was prepared by reverse transcription using the PowerUp SYBR Green master mix for qRT-PCR (Life Technologies, A25742). Real time qPCR was performed on a CFX384 Real time system (BioRad). Data was processed using the BioRad CFX Maestro (BioRad). Primers used are listed in Table S2.

### Western blot assays

Whole hSOs were rapidly lysed by gentle agitation on ice using the RIPA buffer system with protease and a phosphatase inhibitor cocktail (Santacruz, sc-24948A) for HIF1α, RAMP1 and RAMP2 analyses. Whole cell lysates were incubated at 4°C for 1 hr, centrifuged at 14,000 ⅹ g for 15 mins. After, the supernatant was collected, and the protein concentration of the supernatant was measured using the Pierce™ BCA Protein Assay Kits (Thermo Fisher Scientific, 23225). The lysate was denatured in Bolt™ LDS Sample Buffer (Invitrogen, B0007) at 95°C for 5 mins. The samples (at 7 μg) were loaded on a Bolt™ 4-12% Bis-Tris Protein Gels (Invitrogen, NW04120BOX) and then transferred to a PVDF membrane using iBlot™ 2 Transfer Stacks (Invitrogen, IB24002) and an iBlot2 (Method: P3) (Thermo Fisher Scientific, IB24002). After the samples were blocked with 5% skim milk (BD, 232100) in TBST (Tris-buffered saline (Boston BioProducts, BM-301X) containing 0.1% Tween 20 (Sigma-Aldrich, P1379 TBST)) for 1 hr, they were incubated in the blocking solution (TBST containing 5% BSA (Gendepot, A0100-005)) with following primary antibodies: β-actin (anti-mouse, Clone 13E5, 4970S), HIF1α (anti-rabbit, Clone D2U3T, 14179S), RAMP1 (rabbit, 1:500, Abcam, ab156575), RAMP2 (mouse, 1:500, Santacruz, sc-365240), α-tubulin (anti-rat, 1:1000; Abcam, ab6160) at 4°C overnight. After, the blot was washed with 5% TBST, the membranes were incubated in the blocking solution with following horseradish peroxidase-conjugated secondary antibodies: Anti-rat IgG (1:2000; Cell Signaling, #7077), Anti-mouse IgG (1:2000; Cell Signaling, #7076), Anti-rabbit IgG (1:2000; Cell Signaling, #7074) at RT for 1 hr. The SuperSignal™ West Femto Maximum Sensitivity Substrate (Thermo Fisher Scientific, 34095) was used to develop the membranes. The iBright 1500 (Thermo Fisher Scientific, A44114) was used to detect and analyze protein bands. The intensity of each band was quantified with ImageJ (1.53t) software (NIMH, Bethesda, MD) with normalization to background β-actin or α-tubulin.

### Enzyme Immunoassay

Adrenomedullin (ADM) levels were quantified using the competitive enzyme immunoassay for human ADM (Phoenix Pharmaceuticals, EK-010-01) and assays was performed according to the manufacturer’s recommendation. Calculations of sample concentration were performed based on peptide standard solution in the range of 0-100ng/mL. The concentrations of the samples were log_10_ transformed, and concentrations of the samples were interpolated from the sigmoidal curve. Range of reliable detection was between 0.01100 ng/mL. Samples with concentration <0.01ng/mL were quantified as 0.

The quantification of the pAKT:AKT(pS473) ratio and pERK:ERK (pT202/pY204, pT185/pY187) ratio was done using the AKT (Total/Phospho) InstantOne ELISA Kit (Thermo Fisher Scientific, 85-86046-11) for pAKT:AKT, and the ERK1/2 (Total/Phospho) InstantOne ELISA Kit (Thermo Fisher Scientific, 8586013-11) for pERK:ERK. Both assays were performed according to manufacturers’ protocol. For both assays, samples were lysed in a 200μl lysis buffer provided by the manufacturer. Samples were incubated for 1h, and 50μl of lysate per well was used each of the assays. Quantification of cAMP levels was performed using the cAMP Assay Kit (Competitive ELISA) (Abcam, ab65355) according to manufacturers’ protocol. For the analysis, samples were lysed in 250μl lysis buffer and 100μl was used for the assay. The concentration of cAMP was calculated based on the standard curve obtained from kit standards. The quantification of PKA was done using the PKA (Protein Kinase A) Colorimetric Activity Kit (Thermo Fisher Scientific, EIA PKA) according to manufacturers’ protocol. For the assay, samples were lysed in 200μl lysis buffer containing protease inhibitor cocktail (1μl/mL, Sigma-Aldrich, P1860), phenylmethylsulfonyl fluoride (1mM, Sigma-Aldrich, 10837091001), and Activated Sodium Orthovanadate (10mM, Calibiochem, 567540). Samples were incubated for 1h under shaking (250rpm), and centrifuged at 10,000rpm for 10 min at 4°C. Samples were then diluted 1x with kit dilution buffer and 40μl of diluted sample was used for analysis. The quantification of the pCREB:CREB ratio was done using the phosphoELISA CREB (pS133) (Thermo Scientific, KHO0241) and phosphoELISA CREB (Total) (Thermo Scientific, KHO0231) according to manufacturer’s protocol. For both assays, samples were lysed in 100μl RIPA lysis buffer containing phosphatase inhibitor cocktail (Santa Cruz Biotechnology, sc-45065) and a 1:50 dilution of the lysates in standard diluent buffer were used for the assay. The concentration of pCREB and CREB (total) were calculated based on the standard curve obtained from the kit standards using the Tecan Infinite M1000 Pro and i-control 1.10 software.

### Exogenous ADM peptide denaturation and LC-MS analysis

The synthetic peptide Adrenomedullin (ADM) (2 nmol, Anaspec, AS-60447) was incubated with either ddH_2_0 (for control condition) or Dithiothreitol (DTT) (5mM, Sigma Aldrich, D5545), at 56 °C for 30 min. Following reduction of disulfide bridges, alkylation of the free SH-groups was performed Iodoacetamide (IAM) (20mM, Sigma-Aldrich, I1149) at 23 °C for 45 min in the dark. The unreacted IAM was quenched with 5mM final DTT. The solution was desalted and concentrated using Amicon Ultra (3kD, EMD Millipore, UFC500324), and the volume was adjusted to the starting volume with UltraPure DNase/RNase Free Distilled Water (Thermo Fisher Scientific, 10-977-015). LC-MS analysis was performed using Vanquish LC system coupled with an ID-X Tribrid mass spectrometry equipped with a heated electrospray ionization source (HESI-II, Thermo Scientific). For the LC system, a ZORBAX RRHD diphenyl 100 x 2.1 mm column (Agilent, 858750-944) was used. The mobile phases were 0.1% formic acid (Fisher Scientific, A117-50) in ddH_2_O for A and 0.1% formic acid in acetonitrile (Sigma-Aldrich, 34998) for B. The chromatographic gradient was as follows: 0-2min 10% B, linear increase 2-10min up to 60% B, linear increase 10-15min up to 98% B, hold 15-17min at 98% B, linear decrease 17-18min to 10% B, equilibrate 18-21min at 10% B. The mobile phase flow rate was set to 250 μl/min with an injection volume of 3 μL. The autosampler temperature was set to 4 °C, and the column chamber temperature to ambient. Data acquisition on the IDX MS was acquired at 120,000 resolution setting with positive ion voltage of 3.5kV, vaporizer and transfer tube temperature of 325 °C; AGC target set to standard, RF lens set to 45%, sheath gas set to 40 AU, aux gas set to 12 AU, maximum injection time of 100ms, with 5 microscans/acquisition, and scan range of 500-2000 *m/z*. Examination of the LC-MS data was performed using Xcalibur FreeStyle 1.6 (Thermo Scientific). Here, we monitored the formation of carbamidomethyl-cysteines at Cys16-Cys21 (predicted M+1: 6142.9973), or the reduced form of the peptide (predicted M+1: 6028.9544).

### Statistical analyses

Data are presented as mean ± s.e.m. unless otherwise indicated. Distribution of the raw data was tested for normality; statistical analyses were performed as indicated in figure legends. Sample sizes were estimated empirically or based on power calculations. Blinding was used for analyses comparing control and treatment samples.

## List of supplementary material

**Table S1.** List of hiPSC lines and number of cells used for each experiment.

**Table S2.** List of primers used for qPCRs.

**Data File 1.** List of differentially expressed genes by RNA-Sequencing in hSOs exposed to hypoxia.

**Data File 2.** List of GSEA terms from RNA-Sequencing.

## ACKNOWLEDGMENTS

We thank the members of Anca Pasca’s lab and other collaborators for constructive feedback and or editing the manuscript (alphabetical order: Tudor Braicu, Yusuke Hori, Madison House, Geana Marte, Sherry Mestan, Deniz Yagmur Urey, Andrea Navarrette Vargas). We thank Sergiu Pasca for constructive scientific discussion in the early part of developing this project and for allowing us to use the 10X Genomics equipment for library preparation for scRNA-sequencing. We also thank Yuki Miura in Sergiu Pasca’s lab for assisting wih answering questions about the use of equipment during library prep for scRNA-sequencing. Figures were created with Biorender.

## FUNDING

Stanford Maternal & Child Health Research Institute (MCHRI) Postdoctoral Fellowship (to L. L.)

Knut and Alice Wallenberg Foundation Postdoctoral Fellowship (to W.M)

Swedish Research Council IPD (to W.M)

Bill and Melinda Gates Foundation (TO A.M.P.)

## AUTHOR CONTRIBUTIONS

A.P. and W.P.M. contributed equally to the work.

Conceptualization: AMP, AP, WPM, LL, AJW

Methodology: AMP, AP, WPM, DN

Investigation: AMP, AP, WPM, LL, AE, KM, FB, SH, JBC, YD, SP, EG

Visualization: LL, AP, WPM, KM, AE, FB

Funding acquisition: AMP, LL, WPM

Supervision: AP

Writing - original draft: WPM, AP, AMP, KM, AE, AJW, FB

Writing – review and editing: AMP, WPM, AP, LL, AE, KM, FB, JBC, DN, AJW

## DATA AVAILABILITY

Gene expression data are available in the Gene Expression Omnibus (GEO) under accession number GSE183201 (token: odqhewkkrjsdzgd).

---

**Figure S1.**
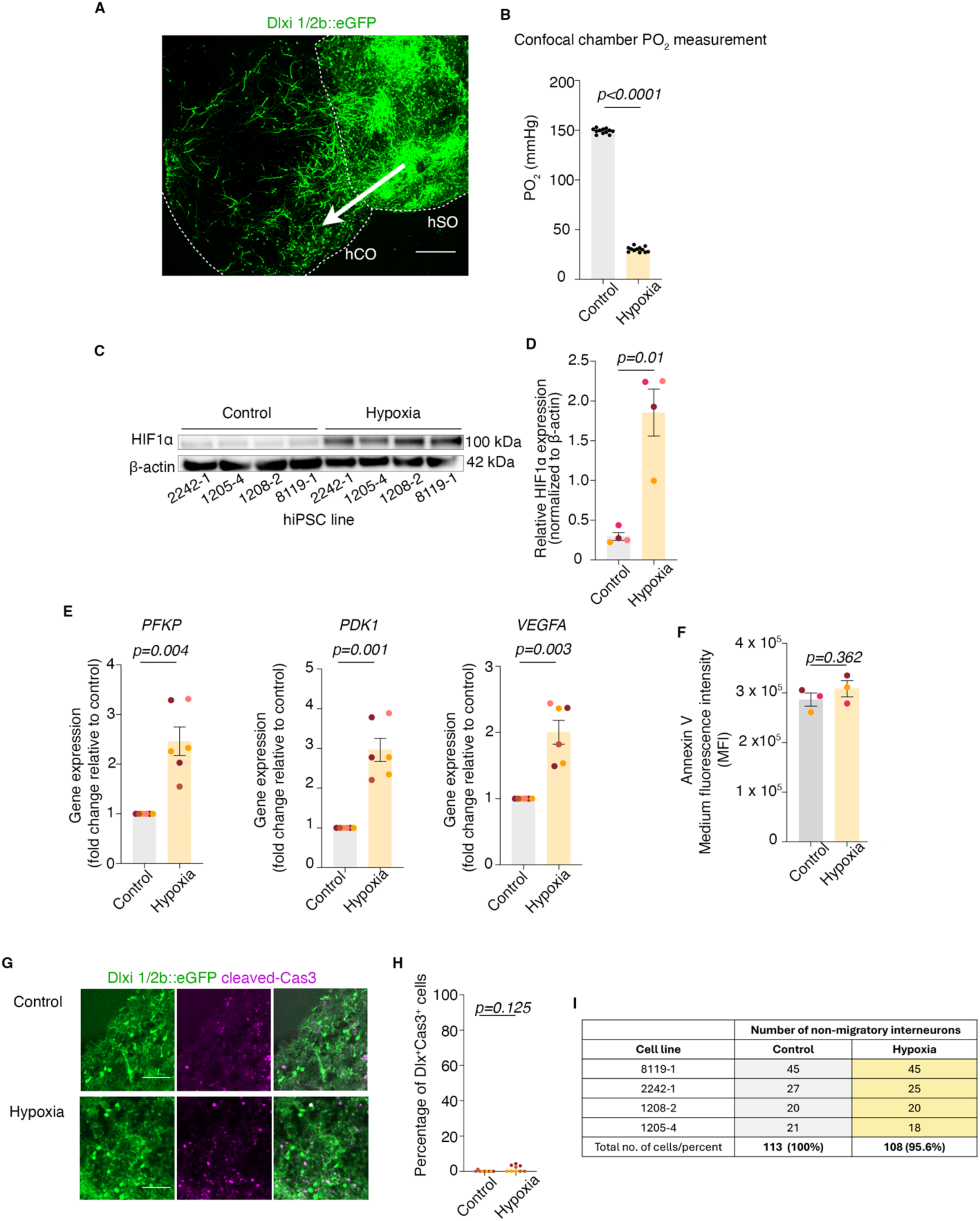
Example of hFA, oxygen level measurements, qPCR changes in expression of hypoxia-responsive genes and cell death analyses. **A.** Example image of hFA showing cortical interneurons migrated from the hSO side to the hCO side; arrow indicates the direction of migration from hSO to hCO; **B.** Oxygen levels (PO_2_ (mmHg)) in cell culture media under control conditions and following exposure to hypoxia for 24 hrs in the confocal microscope environmental chamber (unpaired *t-*test, P<0.0001); **C.** Example of Western Blot showing the stabilization of HIF1α protein in hSOs from 4 hiPSC lines upon exposure to hypoxia, and β-actin expression for normalization; **D.** Quantification of HIF1α protein, normalized to β-actin (two-tailed paired *t*-test; P=0.01); **E.** Transcriptional upregulation of hypoxia-responsive genes in hSO exposed to 24 hrs of hypoxia (<1% O_2_): *PFKP* (two-tailed paired *t*-test, P=0.004), *PDK1* (two-tailed paired *t*-test, P=0.001), *VEGFA* (two-tailed paired *t*-test, P=0.003); **F.** Quantification of Annexin V levels in Dlx^+^ interneurons in hSOs in control and hypoxia conditions (unpaired *t*-test, P=0.362); **G.** Representative image of Dlxi1/2b::eGFP and cleaved-CAS3 positive cells in control and hypoxia conditions; **H.** Quantification (percentage) of Dlxi1/2b::eGFP and cleaved-CAS3 positive cells in control and hypoxia conditions (unpaired *t*-test, P= 0.125); **I.** Table showing the number of non-migratory interneurons from 4 hiPSC lines in control and hypoxia. Bar charts: mean ± s.e.m.; scale bar: 100 μm; Different dot colors represent individual hiPSC lines.

**Figure S2.**
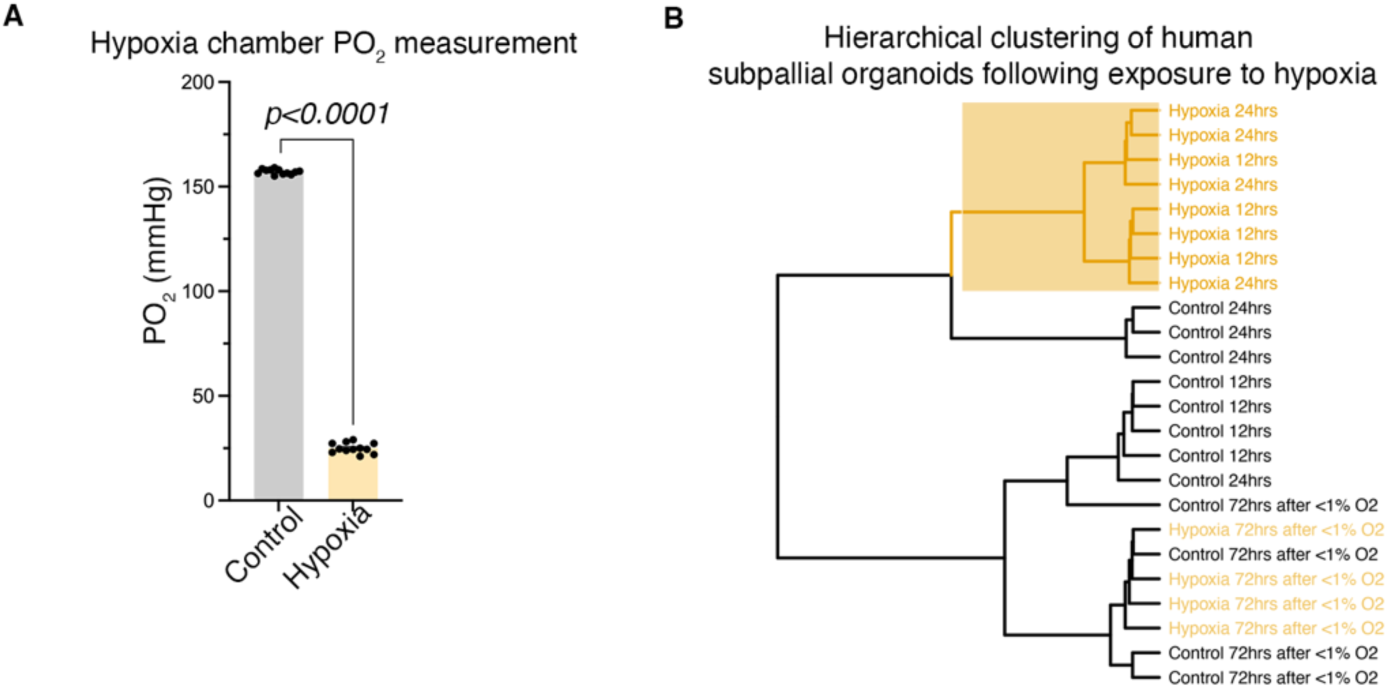
Oxygen level measurements for RNA-Sequencing experiments and dendrogram of sample clustering. **A.** Oxygen levels (PO_2_ (mmHg)) in cell culture media under control conditions and following exposure to hypoxia for 24 hrs in the C-chamber hypoxia sub-chamber (Biospherix) (unpaired *t*-test, P<0.0001); **B.** Hierarchical clustering, using Ward’s criterion, of the first three principal components of the normalized gene expression profiles of all 17,777 genes passing quality control shows clear separation of 12 and 24 hrs hypoxia-exposed samples versus control and from reoxygenated samples. Bar charts: mean±s.e.m.

**Figure S3.**
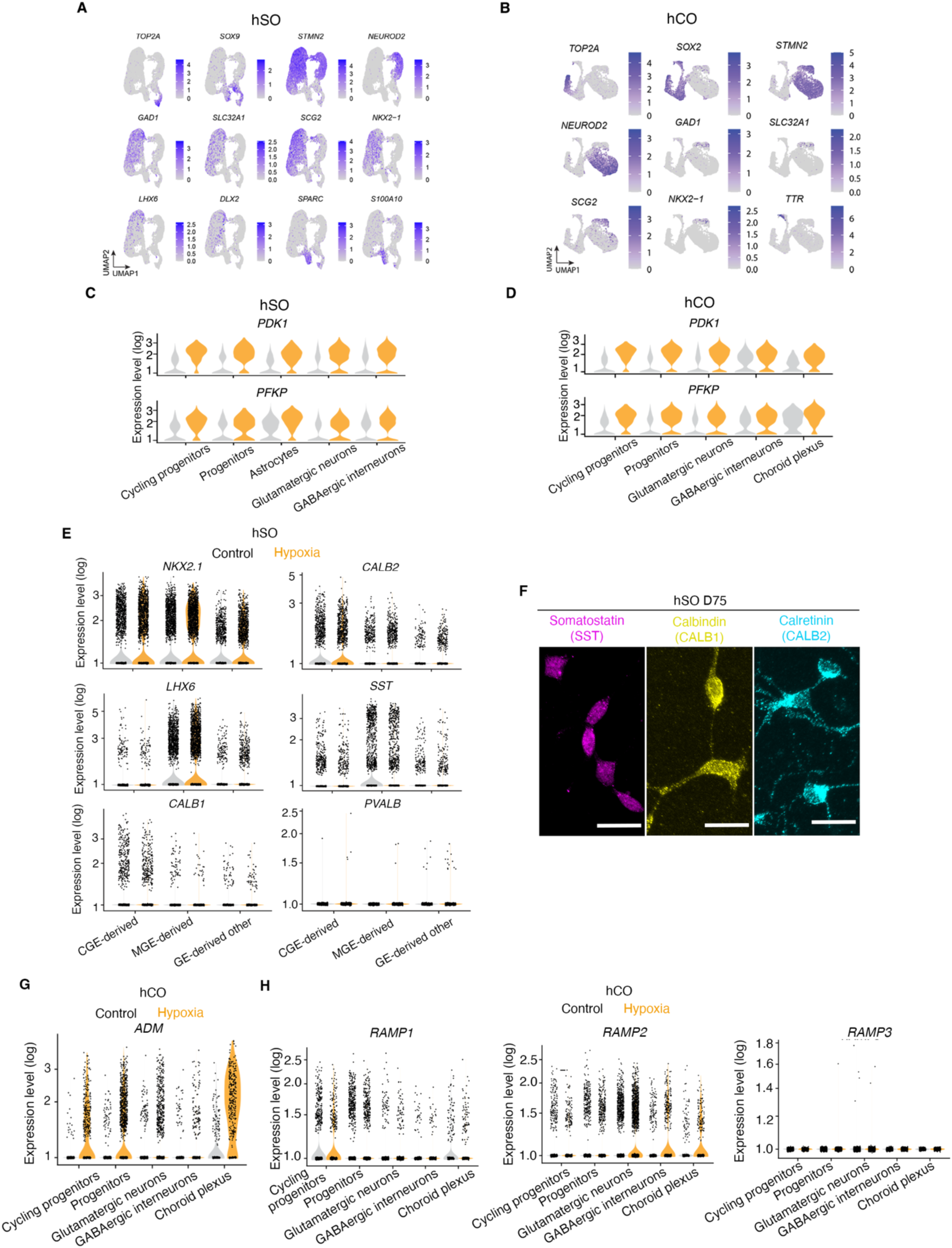
Quality control and robustness analyses for scRNA-sequencing data. **A.** UMAP visualization of main cellular subclusters in hSOs with corresponding gene expression including, astrocytes (*SPARC, S100A10*), progenitors (*SOX9*), cycling progenitors (*SOX9, TOP2A*), glutamatergic neurons (*STMN2, NEUROD2*), interneuron progenitors (*NKX2.1, LHX6, DLX2*), and interneurons (*STMN2, GAD1, SLC32A1, SCG2*); **B.** UMAP visualization of main cellular subclusters in hCO with corresponding gene expression including, choroid plexus (*TTR)*, progenitors (*SOX2*), cycling progenitors (*SOX2, TOP2A*), glutaminergic neurons (*STMN2, NEUROD2*), interneuron progenitors (*NKX2.1*), and interneurons (*STMN2, GAD1, SLC32A1, SCG2*); **C.** Single cell gene expression level (log) of hypoxia-responsive genes (*PDK1* and *PFKP*) across main cellular subclusters in hSO; **D.** Single cell gene expression level (log) of hypoxia-responsive genes (*PDK1* and *PFKP*) across main cell clusters in hCO; **E.** Single cell gene expression level (log) of well-established interneuron markers of MGE and CGE origin (*NKX2.1, CALB2, SST, LHX6*) across interneuron subclusters and additional interneuron markers (*CALB1* and *PVALB*) MGE and CGE interneurons across hSOs; **f.** Single cell gene expression level (log) of well-established interneuron markers of MGE and CGE origin (*NKX2.1, CALB2, SST, LHX6*) across interneuron subclusters and additional interneuron markers (CALB1 and *PVALB*) MGE and CGE interneurons across hCOs; **F.** Example images of interneuron subtype immunostaining in hSOs. Magenta: Somatostatin; Yellow: Calbindin; Cyan: Calretinin. Scale bar: 20 μm; **G.** Single cell gene expression level of ADM in main cell clusters of hCO; **H.** Single cell gene expression level (log) of other *RAMP* family receptors (*RAMP1, RAMP2, RAMP3*) in main cell clusters of hCO. Colors: Gray represents control and yellow represents hypoxia in the graphs.

**Figure S4.**
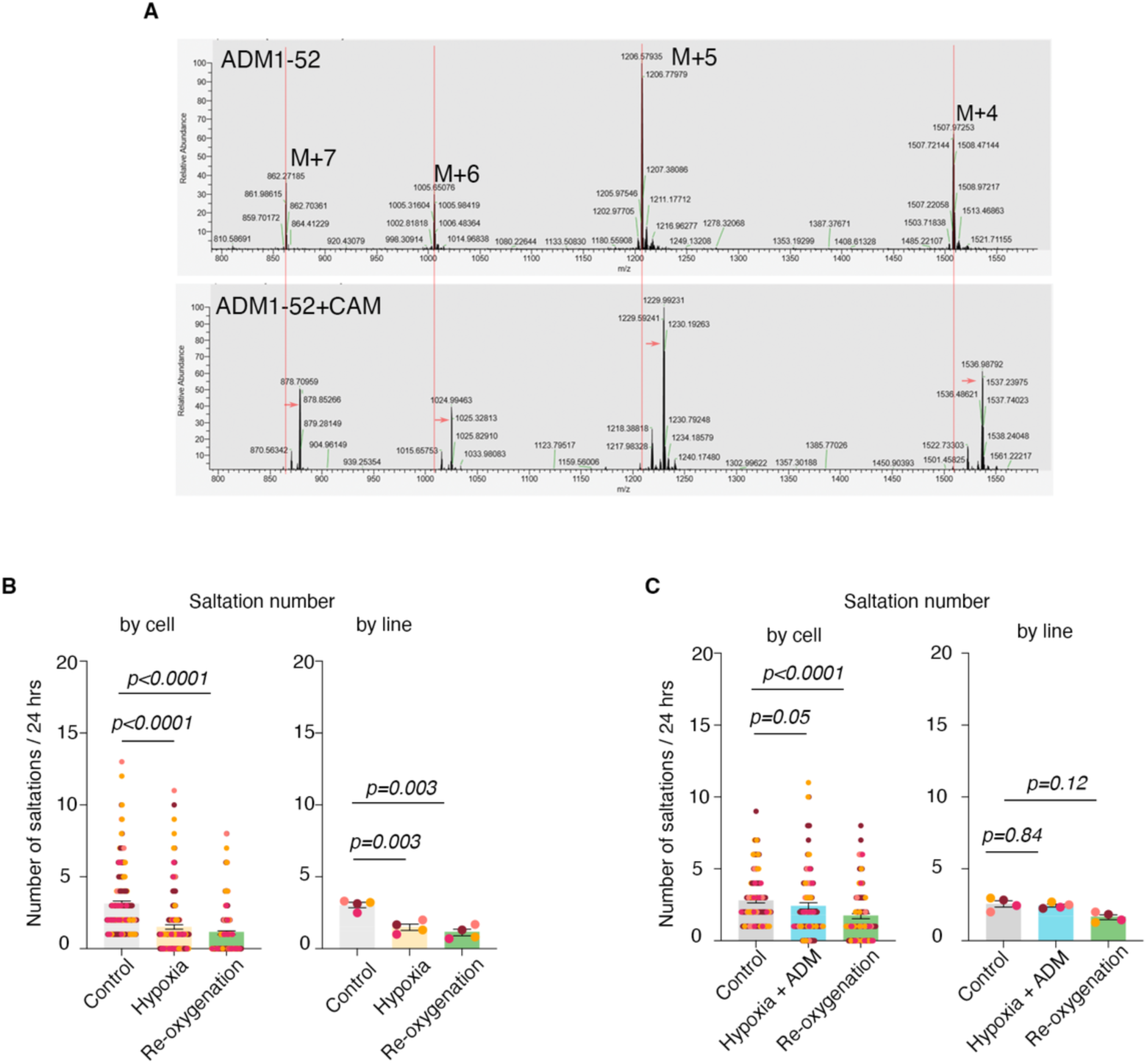
A. LC-MS based analysis of the non-modified ADM (top) and the denatured ADM form (bottom), displays characteristic M+4 through M+7 peaks with two CAM modifications (+114 Da) in the bottom panel represented by the peak shift (arrow); **B. (Left)** Quantification of saltations number/24hr in control versus hypoxia by individual cells (Friedman test, P<0.0001) and control versus reoxygenation by individual cells (Friedman test, P<0.0001); **(Right)** Quantification of saltations/24hr in control versus hypoxia (one-way ANOVA, P=0.003) by hiPSC line and control versus reoxygenation by hiPSC line (one-way ANOVA, P=0.003); **C. (Left)** Quantification of saltations number/24 hrs in control versus hypoxia + ADM condition by individual cells (Friedman test, P=0.05) and in control versus re-oxygenation after hypoxia + ADM by individual cells (Friedman test, P<0.0001); 35% decrease compared to 65% decrease when ADM not added to hypoxia; **(Right)** Quantification of saltations number/24 hrs in control versus hypoxia + ADM condition by hiPSC line (one-way ANOVA, P=0.84) and in control versus re-oxygenation after hypoxia + ADM condition by hiPSC line (one-way ANOVA, P=0.12); Bar charts: mean±s.e.m.; Different dot colors represent individual hiPSC lines.

**Figure S5.**
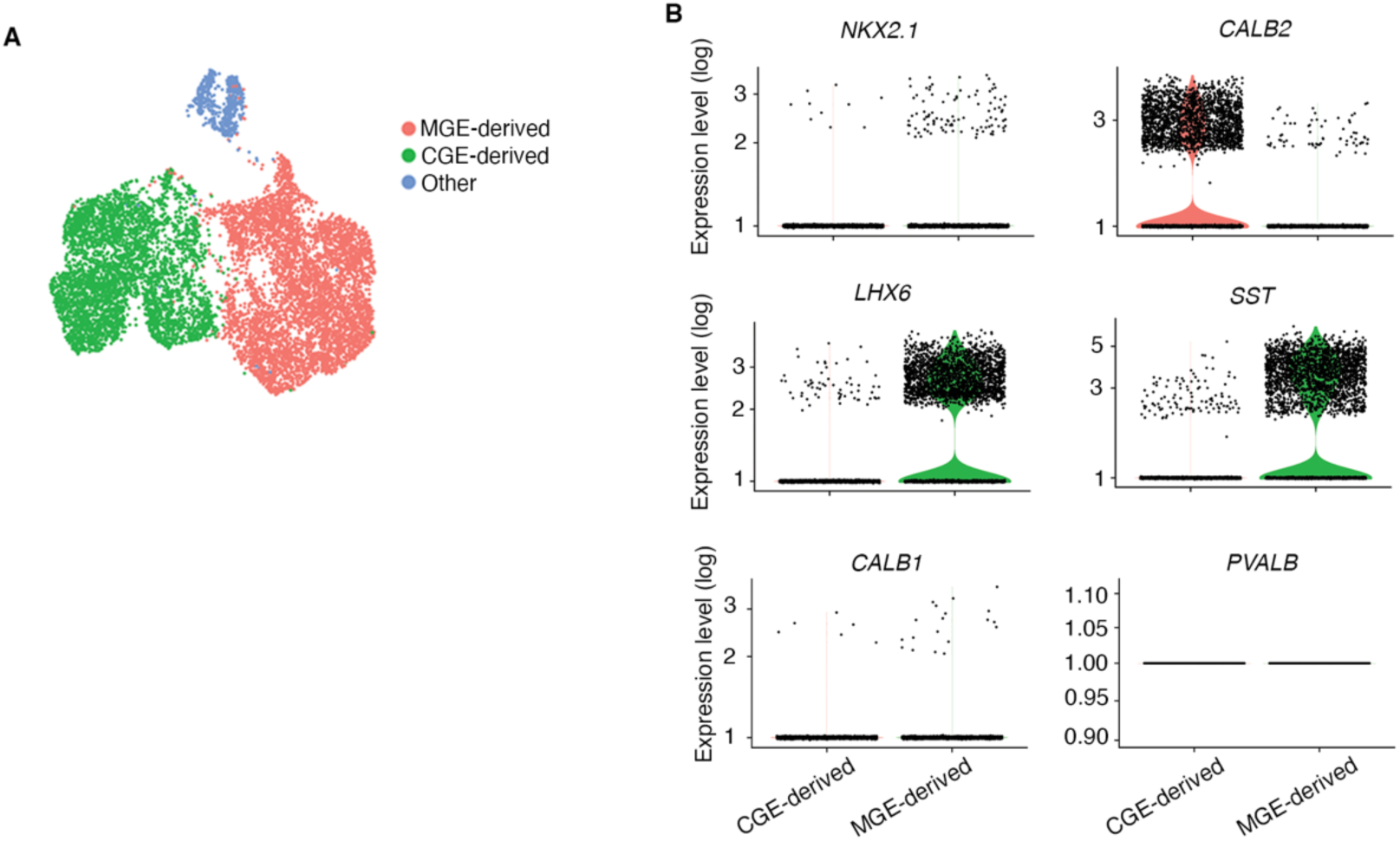
Additional data to support molecular mechanisms of rescue by ADM. **A.** UMAP showing three clusters, including a MGE cluster, a CGE cluster and a third cluster represented by doublets**; B.** Single cell gene expression level (log) of the well-established genes present in the interneurons of MGE and CGE origin (e.g. NKX2.1, CALB2, LHX6, SST) across the interneurons subclusters and Graphical representation of expression of genes associated with GABAergic interneurons subtypes, including *CALB1* and *PVALB*.

**Figure S6.**
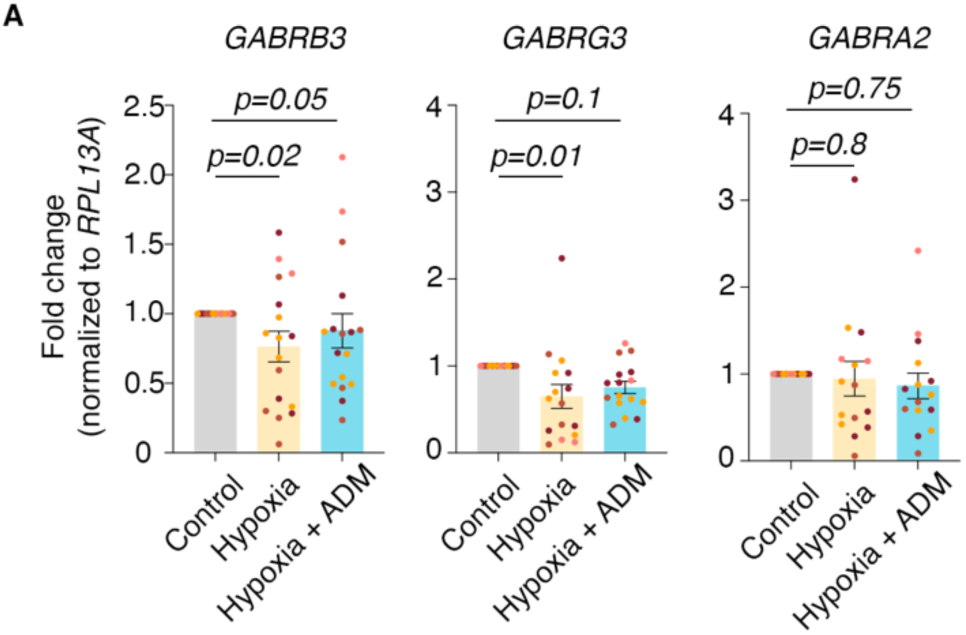
Additional data to support molecular mechanisms of rescue by ADM. A. (Left) Quantification (by q-PCR) of GABRB3 in individual samples in control versus hypoxia conditions (Kruskal-Wallis test, P=0.02) and control versus hypoxia + ADM (Kruskal-Wallis test, P=0.05); **(Center)** Quantification (by q-PCR) of GABRG3 in individual samples in control versus hypoxia conditions (one-way ANOVA test, P=0.01) and control versus hypoxia + ADM (one-way ANOVA test, P=0.1); **(Right)** Quantification (by q-PCR) of GABRA2 in individual samples in control versus hypoxia conditions (one-way ANOVA, P=0.8) and control versus hypoxia + ADM (one-way ANOVA, P=0.75). Bar charts: mean±s.e.m.; Different dot colors represent individual hiPSC lines.

