## Supplementary material for "Adrenomedullin restores the human cortical interneurons migration defects induced by hypoxia": Table S2

**EXTENDED TABLE 2.**

Primers used for qPCR experiments.

| **Gene** | **Forward Primer** | **Reverse Primer** |
| --- | --- | --- |
| RPL13A | CCTGGAGGAGAAGAGGAAAGAGA | TTGAGGACCTCTGTGTATTTGTCAA |
| PFKP | CGCCTACCTCAACGTGGTG | ACCTCCAGAACGAAGGTCCTC |
| PDK1 | CTGTGATACGGATCAGAAACCG | TCCACCAAACAATAAAGAGTGCT |
| ADM | GAACTGCGGATGTCCAG | GTCCTTGTCCTTATCTGTG |
| VEGFA | AGGGCAGAATCATCACGAAGT | AGGGTCTCGATTGGATGGCA |
| CXCR4 | CCCTCCTGCTGACTATTCCC | TAAGGCCAACCATGATGTGC |
| CXCR7 | CAACCTCTTCGGCAGCATTT | AGACGACACGGCGTACCAT |
| CXCL12 | ACTCCAAACTGTGCCCTTCA | CCACTTTAGCTTCGGGTCAAT |
| GABRA1 | GTCACCAGTTTCGGACCCG | AACCGGAGGACTGTCATAGGT |
| GABRA2 | GTTCAAGCTGAATGCCCAAT | ACCTAGAGCCATCAGGAGCA |
| GABRB2 | GCAGAGTGTCAATGACCCTAGT | TGGCAATGTCAATGTTCATCCC |
| GABRG2 | CACAGAAAATGACGGTGTGG | TCACCCTCAGGAACTTTTGG |
| GABRB3 | CAAGCTGTTGAAAGGCTACGA | ACTTCGGAAACCATGTCGATG |
| GABRG3 | AACCAACCACCACGAAGAAGA | CCTCATGTCCAGGAGGGAAT |
