## Supplementary material for "Adrenomedullin restores the human cortical interneurons migration defects induced by hypoxia": Data File S1

| EnsemblID | GeneSymbol | GeneType | log2FoldChange | padj | pvalue | baseMean |
| --- | --- | --- | --- | --- | --- | --- |
| ENSG00000139832 | RAB20 | protein_coding | 2.901248614 | 3.58E-08 | 2.63E-10 | 9.035593747 |
| ENSG00000287560 | AL731533.3 | lncRNA | 2.899471327 | 2.17E-11 | 7.22E-14 | 28.11112511 |
| ENSG00000187017 | ESPN | protein_coding | 2.566920965 | 1.33E-10 | 5.75E-13 | 10.54761269 |
| ENSG00000129521 | EGLN3 | protein_coding | 2.370636304 | 4.26E-14 | 8.23E-17 | 119.8264534 |
| ENSG00000114268 | PFKFB4 | protein_coding | 2.274434387 | 1.84E-19 | 7.09E-23 | 172.6965662 |
| ENSG00000251493 | FOXD1 | protein_coding | 2.25608075 | 0.000320611 | 6.39E-06 | 15.4375431 |
| ENSG00000261051 | AC107021.2 | lncRNA | 2.249336047 | 8.42E-07 | 7.61E-09 | 19.40887726 |
| ENSG00000148926 | ADM | protein_coding | 2.246564807 | 8.97E-06 | 1.00E-07 | 160.240717 |
| ENSG00000146674 | IGFBP3 | protein_coding | 2.205938425 | 0.000122315 | 2.05E-06 | 25.05539141 |
| ENSG00000116016 | EPAS1 | protein_coding | 2.150018872 | 0.000150966 | 2.69E-06 | 9.285674822 |
| ENSG00000114023 | FAM162A | protein_coding | 2.042977068 | 5.91E-18 | 4.56E-21 | 158.1521575 |
| ENSG00000123095 | BHLHE41 | protein_coding | 2.007169904 | 5.17E-08 | 3.87E-10 | 67.68741005 |
| ENSG00000181458 | TMEM45A | protein_coding | 1.976894276 | 2.84E-15 | 4.16E-18 | 27.0594285 |
| ENSG00000104419 | NDRG1 | protein_coding | 1.970442241 | 4.81E-09 | 2.90E-11 | 54.39515885 |
| ENSG00000152952 | PLOD2 | protein_coding | 1.96884566 | 3.32E-15 | 5.13E-18 | 169.9108321 |
| ENSG00000240032 | LNCSRLR | lncRNA | 1.937966986 | 6.27E-06 | 6.69E-08 | 10.63859767 |
| ENSG00000114480 | GBE1 | protein_coding | 1.925203698 | 4.67E-09 | 2.78E-11 | 47.92777741 |
| ENSG00000113083 | LOX | protein_coding | 1.89851873 | 2.15E-13 | 4.64E-16 | 54.04778435 |
| ENSG00000113739 | STC2 | protein_coding | 1.889977341 | 4.30E-22 | 6.64E-26 | 240.8319722 |
| ENSG00000159208 | CIART | protein_coding | 1.853264144 | 4.35E-19 | 2.01E-22 | 60.46813927 |
| ENSG00000105643 | ARRDC2 | protein_coding | 1.846364691 | 2.29E-08 | 1.59E-10 | 10.0829822 |
| ENSG00000279689 | AC022400.9 | TEC | 1.809476039 | 1.67E-07 | 1.39E-09 | 16.74090628 |
| ENSG00000260231 | KDM7A-DT | lncRNA | 1.780465606 | 1.21E-12 | 3.07E-15 | 37.65484129 |
| ENSG00000061656 | SPAG4 | protein_coding | 1.773387995 | 8.41E-08 | 6.75E-10 | 10.15038024 |
| ENSG00000159167 | STC1 | protein_coding | 1.769971962 | 1.71E-10 | 7.65E-13 | 111.2352274 |
| ENSG00000152256 | PDK1 | protein_coding | 1.768777428 | 3.17E-18 | 1.96E-21 | 284.5711845 |
| ENSG00000141526 | SLC16A3 | protein_coding | 1.75212781 | 4.13E-10 | 2.01E-12 | 104.5221512 |
| ENSG00000162772 | ATF3 | protein_coding | 1.747219409 | 3.71E-06 | 3.63E-08 | 28.72586954 |
| ENSG00000075426 | FOSL2 | protein_coding | 1.711266261 | 5.02E-11 | 1.94E-13 | 41.4910812 |
| ENSG00000170525 | PFKFB3 | protein_coding | 1.709246947 | 3.35E-12 | 9.57E-15 | 306.4725882 |
| ENSG00000117394 | SLC2A1 | protein_coding | 1.70857759 | 4.91E-11 | 1.86E-13 | 521.6760366 |
| ENSG00000074800 | ENO1 | protein_coding | 1.683194375 | 4.19E-15 | 6.80E-18 | 2752.213579 |
| ENSG00000135245 | HILPDA | protein_coding | 1.670976763 | 2.02E-18 | 1.09E-21 | 277.8972789 |
| ENSG00000177469 | CAVIN1 | protein_coding | 1.669377075 | 0.036038221 | 0.003692304 | 10.07677265 |
| ENSG00000179403 | VWA1 | protein_coding | 1.62856127 | 1.55E-16 | 1.80E-19 | 15.29387944 |
| ENSG00000128965 | CHAC1 | protein_coding | 1.625684876 | 8.60E-10 | 4.38E-12 | 60.19076831 |
| ENSG00000182379 | NXPH4 | protein_coding | 1.603999661 | 5.82E-09 | 3.68E-11 | 20.18363757 |
| ENSG00000096060 | FKBP5 | protein_coding | 1.599241706 | 0.000321508 | 6.45E-06 | 32.08396657 |
| ENSG00000079739 | PGM1 | protein_coding | 1.588149306 | 1.45E-09 | 8.17E-12 | 39.11295178 |
| ENSG00000067057 | PFKP | protein_coding | 1.586575787 | 1.54E-14 | 2.73E-17 | 191.7869022 |
| ENSG00000159399 | HK2 | protein_coding | 1.578396188 | 5.30E-09 | 3.23E-11 | 801.0974119 |
| ENSG00000122884 | P4HA1 | protein_coding | 1.574832937 | 2.00E-12 | 5.26E-15 | 258.2172105 |
| ENSG00000072682 | P4HA2 | protein_coding | 1.568640337 | 3.00E-08 | 2.16E-10 | 40.33138011 |
| ENSG00000128165 | ADM2 | protein_coding | 1.561483388 | 0.000129257 | 2.21E-06 | 12.20461062 |
| ENSG00000169499 | PLEKHA2 | protein_coding | 1.559123903 | 2.09E-11 | 6.76E-14 | 27.5749119 |
| ENSG00000134107 | BHLHE40 | protein_coding | 1.555855361 | 2.65E-08 | 1.88E-10 | 222.5697042 |
| ENSG00000273001 | AL731533.2 | lncRNA | 1.555320688 | 6.49E-06 | 6.97E-08 | 27.66731694 |
| ENSG00000176171 | BNIP3 | protein_coding | 1.552028162 | 2.60E-13 | 6.03E-16 | 510.0167336 |
| ENSG00000145901 | TNIP1 | protein_coding | 1.53919512 | 3.78E-12 | 1.11E-14 | 103.2217039 |
| ENSG00000161544 | CYGB | protein_coding | 1.538213665 | 4.63E-10 | 2.29E-12 | 15.24227599 |
| ENSG00000162433 | AK4 | protein_coding | 1.512540065 | 9.88E-14 | 2.06E-16 | 162.9262667 |
| ENSG00000115457 | IGFBP2 | protein_coding | 1.510108912 | 6.88E-15 | 1.17E-17 | 731.849513 |
| ENSG00000108001 | EBF3 | protein_coding | 1.50640266 | 0.013874802 | 0.000940053 | 26.35274963 |
| ENSG00000162496 | DHRS3 | protein_coding | 1.505137388 | 0.007157782 | 0.000370268 | 14.4378535 |
| ENSG00000083444 | PLOD1 | protein_coding | 1.49473138 | 3.56E-18 | 2.47E-21 | 93.80587262 |
| ENSG00000186918 | ZNF395 | protein_coding | 1.488075656 | 2.13E-12 | 5.76E-15 | 59.62164675 |
| ENSG00000247095 | MIR210HG | lncRNA | 1.472115544 | 8.80E-10 | 4.55E-12 | 102.7894058 |
| ENSG00000143847 | PPFIA4 | protein_coding | 1.471586049 | 1.24E-09 | 6.70E-12 | 129.8918332 |
| ENSG00000134013 | LOXL2 | protein_coding | 1.457590301 | 0.000592681 | 1.40E-05 | 14.82704483 |
| ENSG00000221869 | CEBPD | protein_coding | 1.451361515 | 0.015558558 | 0.001111154 | 11.11189645 |
| ENSG00000124785 | NRN1 | protein_coding | 1.446781592 | 7.31E-05 | 1.15E-06 | 49.47459388 |
| ENSG00000284138 | ATP6V0CP4 | processed_pseudogene | 1.444181471 | 0.003630506 | 0.000143516 | 13.9717015 |
| ENSG00000171388 | APLN | protein_coding | 1.441073645 | 7.90E-08 | 6.28E-10 | 15.28774365 |
| ENSG00000130766 | SESN2 | protein_coding | 1.432973297 | 3.03E-11 | 1.03E-13 | 83.34525427 |
| ENSG00000151012 | SLC7A11 | protein_coding | 1.424659748 | 1.65E-05 | 2.01E-07 | 52.26048 |
| ENSG00000279692 | AC110285.6 | TEC | 1.419420509 | 0.010419032 | 0.000634699 | 15.96937478 |
| ENSG00000095739 | BAMBI | protein_coding | 1.403465384 | 0.018340016 | 0.001384847 | 12.3502762 |
| ENSG00000220008 | LINGO3 | protein_coding | 1.4017981 | 0.000844316 | 2.10E-05 | 10.00333197 |
| ENSG00000128228 | SDF2L1 | protein_coding | 1.390145898 | 0.00411463 | 0.000172744 | 20.84095382 |
| ENSG00000119938 | PPP1R3C | protein_coding | 1.385742385 | 1.04E-05 | 1.17E-07 | 97.32943805 |
| ENSG00000168209 | DDIT4 | protein_coding | 1.379972344 | 1.46E-10 | 6.41E-13 | 2557.880707 |
| ENSG00000214193 | SH3D21 | protein_coding | 1.379796669 | 2.61E-12 | 7.27E-15 | 80.18205366 |
| ENSG00000101255 | TRIB3 | protein_coding | 1.376632553 | 7.05E-07 | 6.31E-09 | 36.68780737 |
| ENSG00000173281 | PPP1R3B | protein_coding | 1.372577341 | 0.000158174 | 2.83E-06 | 15.55840536 |
| ENSG00000121966 | CXCR4 | protein_coding | 1.357842138 | 1.51E-07 | 1.25E-09 | 417.3630505 |
| ENSG00000100889 | PCK2 | protein_coding | 1.35540452 | 6.21E-06 | 6.57E-08 | 22.07254802 |
| ENSG00000109991 | P2RX3 | protein_coding | 1.352428885 | 0.016882515 | 0.001235687 | 26.99478222 |
| ENSG00000101665 | SMAD7 | protein_coding | 1.352197205 | 5.76E-08 | 4.36E-10 | 39.13003373 |
| ENSG00000182199 | SHMT2 | protein_coding | 1.349417407 | 3.41E-08 | 2.48E-10 | 327.5277462 |
| ENSG00000069667 | RORA | protein_coding | 1.344012683 | 1.07E-11 | 3.38E-14 | 87.58594049 |
| ENSG00000116962 | NID1 | protein_coding | 1.339450517 | 0.030699299 | 0.002924871 | 17.66078099 |
| ENSG00000175197 | DDIT3 | protein_coding | 1.334021469 | 0.00020404 | 3.77E-06 | 59.07072499 |
| ENSG00000060718 | COL11A1 | protein_coding | 1.319477987 | 0.000147878 | 2.60E-06 | 64.09559247 |
| ENSG00000186352 | ANKRD37 | protein_coding | 1.315132572 | 3.22E-06 | 3.06E-08 | 48.12064816 |
| ENSG00000129474 | AJUBA | protein_coding | 1.312913068 | 0.002298148 | 7.82E-05 | 32.06699467 |
| ENSG00000104812 | GYS1 | protein_coding | 1.309892612 | 5.67E-21 | 1.73E-24 | 89.23387114 |
| ENSG00000196968 | FUT11 | protein_coding | 1.30974233 | 2.45E-13 | 5.48E-16 | 179.8858171 |
| ENSG00000172216 | CEBPB | protein_coding | 1.308110968 | 0.000113587 | 1.87E-06 | 87.40411007 |
| ENSG00000197747 | S100A10 | protein_coding | 1.306232555 | 0.01772732 | 0.001320789 | 18.5670255 |
| ENSG00000138119 | MYOF | protein_coding | 1.304530159 | 0.027135033 | 0.002459584 | 27.14478996 |
| ENSG00000051108 | HERPUD1 | protein_coding | 1.299444303 | 5.11E-07 | 4.54E-09 | 166.0984338 |
| ENSG00000105220 | GPI | protein_coding | 1.296919254 | 1.27E-15 | 1.77E-18 | 588.338488 |
| ENSG00000173376 | NDNF | protein_coding | 1.279146214 | 0.000247073 | 4.73E-06 | 30.59426437 |
| ENSG00000144642 | RBMS3 | protein_coding | 1.27536783 | 0.013945835 | 0.000948601 | 9.506898618 |
| ENSG00000142694 | EVA1B | protein_coding | 1.275090811 | 0.027426947 | 0.002498749 | 9.765295591 |
| ENSG00000102144 | PGK1 | protein_coding | 1.267545406 | 1.13E-10 | 4.63E-13 | 810.2196342 |
| ENSG00000247809 | NR2F2-AS1 | lncRNA | 1.241268705 | 0.029834957 | 0.002801392 | 10.80510424 |
| ENSG00000060982 | BCAT1 | protein_coding | 1.23605725 | 2.80E-09 | 1.62E-11 | 374.4461232 |
| ENSG00000140939 | NOL3 | protein_coding | 1.226720692 | 1.45E-09 | 8.18E-12 | 43.72289536 |
| ENSG00000101180 | HRH3 | protein_coding | 1.220001574 | 1.16E-07 | 9.37E-10 | 32.17075076 |
| ENSG00000100219 | XBP1 | protein_coding | 1.211872857 | 0.000350824 | 7.12E-06 | 230.1116078 |
| ENSG00000119950 | MXI1 | protein_coding | 1.205744332 | 4.11E-10 | 1.94E-12 | 100.2756867 |
| ENSG00000087250 | MT3 | protein_coding | 1.199903836 | 2.26E-05 | 2.87E-07 | 25.52068345 |
| ENSG00000272870 | SAP30-DT | lncRNA | 1.199182496 | 1.67E-06 | 1.57E-08 | 18.55698475 |
| ENSG00000146094 | DOK3 | protein_coding | 1.19424739 | 0.003039654 | 0.000113588 | 11.05392823 |
| ENSG00000213700 | RPL17P50 | processed_pseudogene | 1.1905656 | 4.51E-06 | 4.60E-08 | 35.81824146 |
| ENSG00000155660 | PDIA4 | protein_coding | 1.186938695 | 0.0036342 | 0.00014398 | 214.852665 |
| ENSG00000101670 | LIPG | protein_coding | 1.186407365 | 2.40E-05 | 3.05E-07 | 56.98121251 |
| ENSG00000059804 | SLC2A3 | protein_coding | 1.18487945 | 6.95E-08 | 5.37E-10 | 566.4301224 |
| ENSG00000273604 | EPOP | protein_coding | 1.179195625 | 5.81E-09 | 3.59E-11 | 28.36119533 |
| ENSG00000116014 | KISS1R | protein_coding | 1.178893209 | 0.002215407 | 7.41E-05 | 9.11895359 |
| ENSG00000280670 | CCDC163 | protein_coding | 1.17736009 | 0.009423812 | 0.000557711 | 12.05329684 |
| ENSG00000145911 | N4BP3 | protein_coding | 1.17670074 | 1.37E-05 | 1.60E-07 | 31.77790673 |
| ENSG00000124006 | OBSL1 | protein_coding | 1.175654771 | 4.11E-10 | 1.97E-12 | 420.9836696 |
| ENSG00000169299 | PGM2 | protein_coding | 1.171065496 | 1.45E-07 | 1.18E-09 | 17.91198412 |
| ENSG00000099860 | GADD45B | protein_coding | 1.168594181 | 0.020226838 | 0.001597595 | 41.10005736 |
| ENSG00000135766 | EGLN1 | protein_coding | 1.168301009 | 1.11E-09 | 5.82E-12 | 29.86547778 |
| ENSG00000188064 | WNT7B | protein_coding | 1.162878893 | 0.00998461 | 0.000601802 | 11.86406721 |
| ENSG00000155090 | KLF10 | protein_coding | 1.156134302 | 5.67E-21 | 1.75E-24 | 146.9323476 |
| ENSG00000228232 | GAPDHP1 | processed_pseudogene | 1.154494754 | 0.036167383 | 0.003713914 | 9.32648113 |
| ENSG00000125629 | INSIG2 | protein_coding | 1.151215479 | 8.26E-13 | 1.98E-15 | 102.7447238 |
| ENSG00000215474 | SKOR2 | protein_coding | 1.149381597 | 0.042106702 | 0.004700929 | 12.08704814 |
| ENSG00000082684 | SEMA5B | protein_coding | 1.146602076 | 0.001629482 | 4.97E-05 | 103.0648056 |
| ENSG00000188483 | IER5L | protein_coding | 1.138184823 | 1.41E-06 | 1.29E-08 | 97.37478761 |
| ENSG00000167797 | CDK2AP2 | protein_coding | 1.13290361 | 0.000144062 | 2.51E-06 | 88.76360986 |
| ENSG00000067992 | PDK3 | protein_coding | 1.131497722 | 3.39E-11 | 1.18E-13 | 115.1128519 |
| ENSG00000228451 | SDAD1P1 | processed_pseudogene | 1.116838469 | 2.63E-05 | 3.37E-07 | 20.43747752 |
| ENSG00000128590 | DNAJB9 | protein_coding | 1.116824319 | 0.00028517 | 5.57E-06 | 188.2484365 |
| ENSG00000102158 | MAGT1 | protein_coding | 1.111662841 | 0.002035845 | 6.59E-05 | 46.54168667 |
| ENSG00000204592 | HLA-E | protein_coding | 1.111001594 | 0.004423173 | 0.000190901 | 37.62024888 |
| ENSG00000269968 | AC006064.4 | lncRNA | 1.108491627 | 0.024859167 | 0.00217844 | 14.22022644 |
| ENSG00000123689 | G0S2 | protein_coding | 1.107916552 | 0.00079153 | 1.95E-05 | 21.04160237 |
| ENSG00000239911 | PRKAG2-AS1 | lncRNA | 1.105610932 | 0.012335137 | 0.000801106 | 9.277706128 |
| ENSG00000121440 | PDZRN3 | protein_coding | 1.095748378 | 0.008668123 | 0.000491229 | 28.94097236 |
| ENSG00000104313 | EYA1 | protein_coding | 1.091388578 | 0.042384185 | 0.004754804 | 9.811874725 |
| ENSG00000111218 | PRMT8 | protein_coding | 1.091040645 | 0.000194806 | 3.56E-06 | 27.29989524 |
| ENSG00000164649 | CDCA7L | protein_coding | 1.088333071 | 0.000471126 | 1.04E-05 | 20.16881217 |
| ENSG00000166123 | GPT2 | protein_coding | 1.074693263 | 9.17E-18 | 7.78E-21 | 115.1141729 |
| ENSG00000170412 | GPRC5C | protein_coding | 1.074638973 | 0.008885975 | 0.000510071 | 27.38039138 |
| ENSG00000111669 | TPI1 | protein_coding | 1.071918751 | 6.41E-11 | 2.53E-13 | 1632.772301 |
| ENSG00000115461 | IGFBP5 | protein_coding | 1.071707689 | 0.001122045 | 3.00E-05 | 655.0595981 |
| ENSG00000102580 | DNAJC3 | protein_coding | 1.066745169 | 0.00137734 | 4.00E-05 | 83.86490848 |
| ENSG00000110328 | GALNT18 | protein_coding | 1.065817298 | 3.58E-07 | 3.12E-09 | 16.18956611 |
| ENSG00000204525 | HLA-C | protein_coding | 1.059408804 | 0.047493861 | 0.00566309 | 210.0264828 |
| ENSG00000130829 | DUSP9 | protein_coding | 1.058354974 | 0.004350839 | 0.000185428 | 8.948741087 |
| ENSG00000109107 | ALDOC | protein_coding | 1.057278773 | 3.82E-07 | 3.37E-09 | 235.9514524 |
| ENSG00000182704 | TSKU | protein_coding | 1.056202658 | 3.58E-09 | 2.10E-11 | 37.83960892 |
| ENSG00000115548 | KDM3A | protein_coding | 1.050961916 | 1.21E-12 | 3.08E-15 | 422.7029698 |
| ENSG00000287594 | AL356123.2 | lncRNA | 1.046070565 | 0.032964839 | 0.003237944 | 8.619178097 |
| ENSG00000088340 | FER1L4 | unitary_pseudogene | 1.04463867 | 0.004607093 | 0.000202396 | 15.90250243 |
| ENSG00000065911 | MTHFD2 | protein_coding | 1.044456132 | 1.18E-05 | 1.36E-07 | 103.6721337 |
| ENSG00000253554 | LINC01414 | lncRNA | 1.034266365 | 0.034572688 | 0.003462074 | 10.32808086 |
| ENSG00000079308 | TNS1 | protein_coding | 1.032391062 | 5.21E-06 | 5.47E-08 | 75.73864782 |
| ENSG00000135069 | PSAT1 | protein_coding | 1.031996013 | 2.66E-05 | 3.45E-07 | 146.4740672 |
| ENSG00000243364 | EFNA4 | protein_coding | 1.030329935 | 0.006872472 | 0.000348611 | 9.533928373 |
| ENSG00000198363 | ASPH | protein_coding | 1.029836853 | 2.69E-16 | 3.33E-19 | 108.0192014 |
| ENSG00000115738 | ID2 | protein_coding | 1.026514929 | 0.002416734 | 8.28E-05 | 60.83166437 |
| ENSG00000198855 | FICD | protein_coding | 1.026087937 | 0.005290171 | 0.000240982 | 13.71804691 |
| ENSG00000105137 | SYDE1 | protein_coding | 1.023349336 | 0.004094095 | 0.000170693 | 15.2638085 |
| ENSG00000115414 | FN1 | protein_coding | 1.022853634 | 0.023256077 | 0.001972934 | 97.26365198 |
| ENSG00000164105 | SAP30 | protein_coding | 1.022554945 | 1.73E-09 | 9.89E-12 | 59.99988728 |
| ENSG00000145050 | MANF | protein_coding | 1.019448375 | 0.001132048 | 3.04E-05 | 128.8787628 |
| ENSG00000101134 | DOK5 | protein_coding | 1.015077651 | 0.012966415 | 0.000860696 | 12.01639805 |
| ENSG00000185624 | P4HB | protein_coding | 1.007549179 | 0.000306204 | 6.08E-06 | 543.9051112 |
| ENSG00000212743 | LINC02656 | lncRNA | 1.002328081 | 0.034737178 | 0.003486591 | 14.44832992 |
| ENSG00000076554 | TPD52 | protein_coding | 1.000173253 | 0.002979518 | 0.000109961 | 17.729984 |
| ENSG00000171314 | PGAM1 | protein_coding | 0.987514684 | 4.85E-10 | 2.44E-12 | 419.1564285 |
| ENSG00000067225 | PKM | protein_coding | 0.985677357 | 1.00E-11 | 3.09E-14 | 1195.779722 |
| ENSG00000089116 | LHX5 | protein_coding | 0.984008192 | 0.00913543 | 0.000533229 | 29.7885814 |
| ENSG00000167601 | AXL | protein_coding | 0.982220586 | 0.006397718 | 0.00031218 | 28.02258212 |
| ENSG00000150961 | SEC24D | protein_coding | 0.982059429 | 0.008151268 | 0.000451869 | 38.54364753 |
| ENSG00000078114 | NEBL | protein_coding | 0.977235691 | 0.009423812 | 0.000558065 | 162.2526763 |
| ENSG00000165349 | SLC7A3 | protein_coding | 0.973754653 | 0.002975811 | 0.000109594 | 34.88476725 |
| ENSG00000104765 | BNIP3L | protein_coding | 0.973627941 | 1.16E-10 | 4.92E-13 | 487.0234231 |
| ENSG00000069998 | HDHD5 | protein_coding | 0.971865089 | 1.36E-09 | 7.43E-12 | 51.69336703 |
| ENSG00000167771 | RCOR2 | protein_coding | 0.971836646 | 4.39E-14 | 8.80E-17 | 316.3053588 |
| ENSG00000149925 | ALDOA | protein_coding | 0.971598437 | 6.08E-05 | 8.96E-07 | 22.16815644 |
| ENSG00000187840 | EIF4EBP1 | protein_coding | 0.971464089 | 0.000365055 | 7.58E-06 | 40.57944762 |
| ENSG00000177606 | JUN | protein_coding | 0.97085139 | 0.0002956 | 5.82E-06 | 282.7235497 |
| ENSG00000184441 | AP001062.1 | lncRNA | 0.967415992 | 0.042392658 | 0.004771897 | 11.5342334 |
| ENSG00000111981 | ULBP1 | protein_coding | 0.966138336 | 0.005023201 | 0.00022377 | 10.37458752 |
| ENSG00000154721 | JAM2 | protein_coding | 0.962613216 | 0.002980012 | 0.000110328 | 65.37208265 |
| ENSG00000260400 | AL513534.3 | lncRNA | 0.962232056 | 0.000150966 | 2.69E-06 | 78.7410375 |
| ENSG00000182158 | CREB3L2 | protein_coding | 0.959737663 | 0.002909202 | 0.000106018 | 37.1672561 |
| ENSG00000156515 | HK1 | protein_coding | 0.95955903 | 1.20E-09 | 6.37E-12 | 291.6217395 |
| ENSG00000187498 | COL4A1 | protein_coding | 0.958997251 | 0.047012857 | 0.005553557 | 99.49858176 |
| ENSG00000133639 | BTG1 | protein_coding | 0.953305487 | 7.03E-09 | 4.56E-11 | 675.8130629 |
| ENSG00000147872 | PLIN2 | protein_coding | 0.95128315 | 0.001827432 | 5.74E-05 | 16.04527025 |
| ENSG00000138640 | FAM13A | protein_coding | 0.944336239 | 4.64E-05 | 6.41E-07 | 51.91913578 |
| ENSG00000132199 | ENOSF1 | protein_coding | 0.944181123 | 0.004555366 | 0.000198384 | 19.2077654 |
| ENSG00000006459 | KDM7A | protein_coding | 0.943134154 | 7.70E-11 | 3.09E-13 | 65.83927693 |
| ENSG00000089159 | PXN | protein_coding | 0.939173716 | 5.43E-05 | 7.89E-07 | 53.0840894 |
| ENSG00000125945 | ZNF436 | protein_coding | 0.938379545 | 4.84E-11 | 1.79E-13 | 143.441372 |
| ENSG00000026025 | VIM | protein_coding | 0.932990509 | 0.001471372 | 4.38E-05 | 3874.138609 |
| ENSG00000092621 | PHGDH | protein_coding | 0.93136028 | 1.65E-06 | 1.55E-08 | 82.22017 |
| ENSG00000100968 | NFATC4 | protein_coding | 0.929325699 | 0.000250326 | 4.81E-06 | 41.960751 |
| ENSG00000117298 | ECE1 | protein_coding | 0.926569545 | 0.001827432 | 5.74E-05 | 53.62261125 |
| ENSG00000213553 | RPLP0P6 | processed_pseudogene | 0.922707095 | 0.034923796 | 0.003513412 | 12.83873558 |
| ENSG00000254815 | LMNTD2-AS1 | lncRNA | 0.918038977 | 0.002733703 | 9.71E-05 | 64.96960079 |
| ENSG00000080200 | CRYBG3 | protein_coding | 0.915447216 | 0.000770101 | 1.88E-05 | 39.01990366 |
| ENSG00000111640 | GAPDH | protein_coding | 0.914740932 | 7.53E-12 | 2.27E-14 | 6610.841502 |
| ENSG00000197355 | UAP1L1 | protein_coding | 0.91248731 | 0.017861029 | 0.00133351 | 24.67448383 |
| ENSG00000100058 | CRYBB2P1 | unprocessed_pseudogene | 0.910824697 | 4.61E-06 | 4.74E-08 | 31.53033249 |
| ENSG00000162373 | BEND5 | protein_coding | 0.910678414 | 3.75E-06 | 3.70E-08 | 32.08237364 |
| ENSG00000146278 | PNRC1 | protein_coding | 0.909636363 | 2.66E-10 | 1.23E-12 | 710.2209422 |
| ENSG00000178425 | NT5DC1 | protein_coding | 0.909202386 | 0.011367567 | 0.000716178 | 10.80805957 |
| ENSG00000197930 | ERO1A | protein_coding | 0.90245154 | 8.86E-16 | 1.16E-18 | 179.6046575 |
| ENSG00000144579 | CTDSP1 | protein_coding | 0.900314515 | 0.003057963 | 0.000114508 | 39.94347684 |
| ENSG00000110492 | MDK | protein_coding | 0.899563892 | 0.004074246 | 0.000168992 | 159.4509994 |
| ENSG00000128285 | MCHR1 | protein_coding | 0.896957583 | 0.029244127 | 0.002716285 | 17.65511354 |
| ENSG00000001617 | SEMA3F | protein_coding | 0.895614556 | 5.19E-05 | 7.34E-07 | 20.37556398 |
| ENSG00000214063 | TSPAN4 | protein_coding | 0.892070212 | 2.64E-05 | 3.40E-07 | 22.68112051 |
| ENSG00000134333 | LDHA | protein_coding | 0.891552808 | 0.021799016 | 0.001798303 | 237.0467607 |
| ENSG00000135116 | HRK | protein_coding | 0.889963289 | 0.032086967 | 0.003106131 | 14.65177112 |
| ENSG00000166450 | PRTG | protein_coding | 0.886166016 | 0.00402151 | 0.000165493 | 89.61901629 |
| ENSG00000026559 | KCNG1 | protein_coding | 0.885305439 | 0.02091437 | 0.001695498 | 16.33026086 |
| ENSG00000105281 | SLC1A5 | protein_coding | 0.883519932 | 0.009866052 | 0.00059111 | 64.69093463 |
| ENSG00000164096 | C4orf3 | protein_coding | 0.882884185 | 5.82E-09 | 3.66E-11 | 213.7721737 |
| ENSG00000166130 | IKBIP | protein_coding | 0.882450357 | 0.000734494 | 1.79E-05 | 15.96340118 |
| ENSG00000135362 | PRR5L | protein_coding | 0.882006518 | 0.006565635 | 0.000323173 | 16.87129527 |
| ENSG00000101400 | SNTA1 | protein_coding | 0.879801999 | 0.003760223 | 0.000151547 | 9.786531084 |
| ENSG00000204767 | INSYN2B | protein_coding | 0.878179907 | 0.006839845 | 0.000346428 | 15.70876361 |
| ENSG00000259820 | AC083843.3 | lncRNA | 0.877393871 | 0.000397374 | 8.44E-06 | 31.45014997 |
| ENSG00000128050 | PAICS | protein_coding | 0.875595003 | 1.27E-08 | 8.43E-11 | 232.9030491 |
| ENSG00000118707 | TGIF2 | protein_coding | 0.868482284 | 0.006705747 | 0.000335729 | 41.23087342 |
| ENSG00000112559 | MDFI | protein_coding | 0.868100564 | 0.027218107 | 0.002469215 | 19.49698796 |
| ENSG00000172638 | EFEMP2 | protein_coding | 0.867069941 | 0.006839845 | 0.000346354 | 22.6815069 |
| ENSG00000177106 | EPS8L2 | protein_coding | 0.86631119 | 0.004607093 | 0.000202296 | 10.87442514 |
| ENSG00000167397 | VKORC1 | protein_coding | 0.864836082 | 0.001674544 | 5.12E-05 | 13.34078617 |
| ENSG00000198856 | OSTC | protein_coding | 0.864791329 | 0.001436261 | 4.25E-05 | 59.38713857 |
| ENSG00000230330 | HMGN2P3 | processed_pseudogene | 0.863674747 | 0.040168351 | 0.004372867 | 16.52302825 |
| ENSG00000127863 | TNFRSF19 | protein_coding | 0.860898518 | 0.024859167 | 0.002176983 | 19.70911668 |
| ENSG00000112715 | VEGFA | protein_coding | 0.860566303 | 0.000112307 | 1.84E-06 | 955.5869294 |
| ENSG00000166171 | DPCD | protein_coding | 0.859225519 | 1.10E-05 | 1.25E-07 | 27.46788936 |
| ENSG00000134109 | EDEM1 | protein_coding | 0.856371174 | 4.40E-05 | 6.05E-07 | 68.76669739 |
| ENSG00000099849 | RASSF7 | protein_coding | 0.854411179 | 0.005498217 | 0.000253029 | 34.94655251 |
| ENSG00000107731 | UNC5B | protein_coding | 0.854162632 | 0.014300698 | 0.000983425 | 37.69031299 |
| ENSG00000225697 | SLC26A6 | protein_coding | 0.854007374 | 0.000212887 | 3.96E-06 | 28.06574291 |
| ENSG00000105642 | KCNN1 | protein_coding | 0.852785208 | 6.31E-05 | 9.45E-07 | 31.44242815 |
| ENSG00000172456 | FGGY | protein_coding | 0.851974905 | 0.015256043 | 0.001084838 | 14.42246248 |
| ENSG00000259865 | AL390728.6 | lncRNA | 0.844799935 | 0.024254336 | 0.002102583 | 61.04906557 |
| ENSG00000181201 | H2BU2P | unitary_pseudogene | 0.840782948 | 0.002772036 | 9.89E-05 | 56.96618318 |
| ENSG00000116685 | KIAA2013 | protein_coding | 0.837207475 | 2.08E-08 | 1.43E-10 | 86.2067261 |
| ENSG00000100427 | MLC1 | protein_coding | 0.835156839 | 0.036038221 | 0.003687614 | 29.77502666 |
| ENSG00000143870 | PDIA6 | protein_coding | 0.833603369 | 0.001891697 | 6.00E-05 | 213.4528683 |
| ENSG00000181826 | RELL1 | protein_coding | 0.832199826 | 0.017591752 | 0.001307972 | 23.19039529 |
| ENSG00000105202 | FBL | protein_coding | 0.830353688 | 4.28E-05 | 5.81E-07 | 115.8078053 |
| ENSG00000215244 | LINC02649 | lncRNA | 0.826732804 | 0.039966333 | 0.004338532 | 10.18260562 |
| ENSG00000169136 | ATF5 | protein_coding | 0.825686566 | 0.001471372 | 4.39E-05 | 89.2390411 |
| ENSG00000130702 | LAMA5 | protein_coding | 0.822836862 | 6.79E-05 | 1.04E-06 | 97.53455334 |
| ENSG00000166598 | HSP90B1 | protein_coding | 0.818514212 | 0.025486388 | 0.002245211 | 651.0440225 |
| ENSG00000173530 | TNFRSF10D | protein_coding | 0.817184612 | 0.032666478 | 0.003177869 | 11.89271765 |
| ENSG00000141682 | PMAIP1 | protein_coding | 0.808690633 | 0.027331361 | 0.002481947 | 20.97746125 |
| ENSG00000148700 | ADD3 | protein_coding | 0.808494856 | 0.00718316 | 0.000372329 | 44.00455345 |
| ENSG00000103257 | SLC7A5 | protein_coding | 0.806706528 | 4.77E-05 | 6.63E-07 | 174.769364 |
| ENSG00000151092 | NGLY1 | protein_coding | 0.802571525 | 4.94E-06 | 5.15E-08 | 27.42114777 |
| ENSG00000066468 | FGFR2 | protein_coding | 0.802294286 | 0.024181148 | 0.002083552 | 53.20026315 |
| ENSG00000157613 | CREB3L1 | protein_coding | 0.80053185 | 0.035284705 | 0.00356879 | 9.170525981 |
| ENSG00000261829 | AC009407.1 | lncRNA | 0.796656495 | 0.043302174 | 0.004944712 | 19.15146722 |
| ENSG00000064651 | SLC12A2 | protein_coding | 0.795060037 | 1.75E-05 | 2.16E-07 | 61.00062301 |
| ENSG00000143499 | SMYD2 | protein_coding | 0.793450654 | 0.000869194 | 2.19E-05 | 19.73423417 |
| ENSG00000107077 | KDM4C | protein_coding | 0.791002094 | 1.40E-08 | 9.39E-11 | 58.97131133 |
| ENSG00000069399 | BCL3 | protein_coding | 0.790416574 | 0.032927908 | 0.003228725 | 8.653538209 |
| ENSG00000132432 | SEC61G | protein_coding | 0.789995314 | 5.91E-08 | 4.52E-10 | 135.6031329 |
| ENSG00000272842 | AL391834.1 | lncRNA | 0.787659179 | 0.049367096 | 0.005999078 | 23.71617589 |
| ENSG00000102230 | PCYT1B | protein_coding | 0.785803965 | 0.001222338 | 3.32E-05 | 32.97661777 |
| ENSG00000149428 | HYOU1 | protein_coding | 0.782967095 | 0.014005601 | 0.000953748 | 396.3537328 |
| ENSG00000109062 | SLC9A3R1 | protein_coding | 0.781256009 | 0.00127038 | 3.55E-05 | 25.94813389 |
| ENSG00000176720 | BOK | protein_coding | 0.779644927 | 0.043535666 | 0.004988182 | 13.16938564 |
| ENSG00000168214 | RBPJ | protein_coding | 0.777919837 | 7.91E-09 | 5.19E-11 | 337.4209438 |
| ENSG00000185633 | NDUFA4L2 | protein_coding | 0.777433499 | 0.006705747 | 0.000335922 | 17.67992347 |
| ENSG00000123131 | PRDX4 | protein_coding | 0.773544208 | 0.00179087 | 5.57E-05 | 59.12029519 |
| ENSG00000157240 | FZD1 | protein_coding | 0.772304221 | 0.022587282 | 0.001891116 | 113.1855034 |
| ENSG00000236404 | VLDLR-AS1 | lncRNA | 0.771004305 | 0.002417365 | 8.31E-05 | 12.05032036 |
| ENSG00000064042 | LIMCH1 | protein_coding | 0.768697428 | 0.032885616 | 0.0032195 | 138.7957342 |
| ENSG00000197837 | H4-16 | protein_coding | 0.766807158 | 0.002024986 | 6.53E-05 | 67.0687408 |
| ENSG00000257365 | FNTB | protein_coding | 0.766764838 | 0.004603028 | 0.000201507 | 8.855388249 |
| ENSG00000227063 | RPL41P1 | processed_pseudogene | 0.765743324 | 0.008128819 | 0.000449097 | 292.7669064 |
| ENSG00000183098 | GPC6 | protein_coding | 0.765089373 | 0.001767155 | 5.47E-05 | 86.75495236 |
| ENSG00000164442 | CITED2 | protein_coding | 0.764102057 | 0.000483977 | 1.08E-05 | 335.9588039 |
| ENSG00000075420 | FNDC3B | protein_coding | 0.762353037 | 0.036418898 | 0.003760318 | 106.1051 |
| ENSG00000140534 | TICRR | protein_coding | 0.759777346 | 0.032067679 | 0.003094858 | 19.33795161 |
| ENSG00000100100 | PIK3IP1 | protein_coding | 0.759541502 | 0.0013417 | 3.79E-05 | 23.78313797 |
| ENSG00000175105 | ZNF654 | protein_coding | 0.758548499 | 2.28E-10 | 1.04E-12 | 53.07668821 |
| ENSG00000026297 | RNASET2 | protein_coding | 0.758492158 | 0.004178631 | 0.000176476 | 16.16787141 |
| ENSG00000109519 | GRPEL1 | protein_coding | 0.755185492 | 1.68E-07 | 1.41E-09 | 41.48472798 |
| ENSG00000181788 | SIAH2 | protein_coding | 0.753558857 | 0.001281489 | 3.61E-05 | 55.15970418 |
| ENSG00000049449 | RCN1 | protein_coding | 0.752704589 | 0.020923148 | 0.001697825 | 97.67346065 |
| ENSG00000120896 | SORBS3 | protein_coding | 0.751961125 | 0.042124567 | 0.004722427 | 26.42543651 |
| ENSG00000152284 | TCF7L1 | protein_coding | 0.748131617 | 0.0296098 | 0.002757058 | 36.06716391 |
| ENSG00000272606 | AC015982.2 | lncRNA | 0.748013125 | 0.033568659 | 0.003335613 | 25.05023595 |
| ENSG00000130707 | ASS1 | protein_coding | 0.74439341 | 0.007855105 | 0.000424535 | 21.69150942 |
| ENSG00000164111 | ANXA5 | protein_coding | 0.744153025 | 0.047266491 | 0.005598116 | 64.65459211 |
| ENSG00000139641 | ESYT1 | protein_coding | 0.743891107 | 0.001354412 | 3.85E-05 | 60.40123465 |
| ENSG00000240972 | MIF | protein_coding | 0.743656708 | 0.00786079 | 0.000425449 | 34.20375714 |
| ENSG00000275074 | NUDT18 | protein_coding | 0.742315579 | 0.013163023 | 0.000881126 | 14.9225951 |
| ENSG00000204856 | FAM216A | protein_coding | 0.741911547 | 0.001003122 | 2.64E-05 | 30.60040635 |
| ENSG00000087903 | RFX2 | protein_coding | 0.741410411 | 0.047066432 | 0.005568696 | 26.27470109 |
| ENSG00000131652 | THOC6 | protein_coding | 0.739993282 | 0.000360462 | 7.40E-06 | 32.9460179 |
| ENSG00000166046 | TCP11L2 | protein_coding | 0.738620763 | 3.71E-06 | 3.64E-08 | 25.4601816 |
| ENSG00000168894 | RNF181 | protein_coding | 0.734205145 | 9.29E-05 | 1.50E-06 | 166.3262591 |
| ENSG00000236552 | RPL13AP5 | processed_pseudogene | 0.734058811 | 0.046862513 | 0.005530498 | 28.35145166 |
| ENSG00000152518 | ZFP36L2 | protein_coding | 0.733653022 | 0.006705747 | 0.000336414 | 171.4628818 |
| ENSG00000182224 | CYB5D1 | protein_coding | 0.731401523 | 0.026050511 | 0.002318606 | 12.68807393 |
| ENSG00000183258 | DDX41 | protein_coding | 0.73074729 | 1.09E-16 | 1.10E-19 | 181.1756561 |
| ENSG00000142798 | HSPG2 | protein_coding | 0.728861814 | 0.040713872 | 0.004481004 | 45.47321284 |
| ENSG00000115318 | LOXL3 | protein_coding | 0.725963176 | 8.59E-05 | 1.37E-06 | 43.64961272 |
| ENSG00000170779 | CDCA4 | protein_coding | 0.725682944 | 0.031651783 | 0.00304176 | 21.09104957 |
| ENSG00000114315 | HES1 | protein_coding | 0.72419873 | 0.018938876 | 0.001452 | 147.6343591 |
| ENSG00000141580 | WDR45B | protein_coding | 0.723388598 | 0.000122315 | 2.05E-06 | 141.4596073 |
| ENSG00000177453 | NIM1K | protein_coding | 0.721774098 | 0.005042355 | 0.000225411 | 13.3043006 |
| ENSG00000109501 | WFS1 | protein_coding | 0.721332651 | 0.049925734 | 0.006103619 | 30.37583976 |
| ENSG00000106638 | TBL2 | protein_coding | 0.721060256 | 2.78E-05 | 3.63E-07 | 59.88855667 |
| ENSG00000166794 | PPIB | protein_coding | 0.719326738 | 0.0036342 | 0.000144223 | 252.1307953 |
| ENSG00000125844 | RRBP1 | protein_coding | 0.718187661 | 0.02165112 | 0.001771959 | 45.51509414 |
| ENSG00000135070 | ISCA1 | protein_coding | 0.717550698 | 2.16E-07 | 1.83E-09 | 73.62527477 |
| ENSG00000205485 | AC004980.1 | unprocessed_pseudogene | 0.715642614 | 0.009959061 | 0.000598989 | 9.281321178 |
| ENSG00000037637 | FBXO42 | protein_coding | 0.714462998 | 1.16E-10 | 4.87E-13 | 96.84884636 |
| ENSG00000099991 | CABIN1 | protein_coding | 0.712464491 | 2.48E-08 | 1.75E-10 | 195.2797802 |
| ENSG00000167004 | PDIA3 | protein_coding | 0.710409263 | 0.001432815 | 4.23E-05 | 248.3864589 |
| ENSG00000113407 | TARS1 | protein_coding | 0.709604397 | 6.83E-05 | 1.06E-06 | 151.0535115 |
| ENSG00000120254 | MTHFD1L | protein_coding | 0.709512438 | 0.000699347 | 1.69E-05 | 42.94802125 |
| ENSG00000105722 | ERF | protein_coding | 0.708757526 | 0.029766993 | 0.002785485 | 57.76721075 |
| ENSG00000110619 | CARS1 | protein_coding | 0.708401694 | 0.000349286 | 7.06E-06 | 92.29540616 |
| ENSG00000105835 | NAMPT | protein_coding | 0.706721584 | 3.05E-05 | 4.05E-07 | 88.64818538 |
| ENSG00000153113 | CAST | protein_coding | 0.705349647 | 0.028395745 | 0.00262209 | 43.26975866 |
| ENSG00000100908 | EMC9 | protein_coding | 0.702201182 | 0.000586566 | 1.37E-05 | 67.51957784 |
| ENSG00000255717 | SNHG1 | lncRNA | 0.701826056 | 0.000114072 | 1.88E-06 | 255.2820205 |
| ENSG00000183010 | PYCR1 | protein_coding | 0.700325976 | 0.002736679 | 9.74E-05 | 53.64029459 |
| ENSG00000168003 | SLC3A2 | protein_coding | 0.699961925 | 0.006611097 | 0.000328717 | 185.171153 |
| ENSG00000043514 | TRIT1 | protein_coding | 0.69978641 | 2.84E-05 | 3.75E-07 | 54.31139488 |
| ENSG00000134198 | TSPAN2 | protein_coding | 0.699665991 | 0.038506644 | 0.004084839 | 16.97387535 |
| ENSG00000160789 | LMNA | protein_coding | 0.698996433 | 0.000586566 | 1.38E-05 | 54.55687836 |
| ENSG00000157483 | MYO1E | protein_coding | 0.69877339 | 0.007204443 | 0.000375463 | 31.64588123 |
| ENSG00000103507 | BCKDK | protein_coding | 0.691827507 | 4.96E-08 | 3.67E-10 | 66.62730661 |
| ENSG00000179361 | ARID3B | protein_coding | 0.691063938 | 1.20E-05 | 1.39E-07 | 55.4526045 |
| ENSG00000116761 | CTH | protein_coding | 0.688153086 | 0.045140183 | 0.005238241 | 11.87620942 |
| ENSG00000196781 | TLE1 | protein_coding | 0.687920033 | 0.003400439 | 0.000132527 | 51.28108396 |
| ENSG00000189337 | KAZN | protein_coding | 0.686885255 | 0.001495628 | 4.47E-05 | 85.03881929 |
| ENSG00000165195 | PIGA | protein_coding | 0.686353992 | 0.001569041 | 4.75E-05 | 38.73462447 |
| ENSG00000004455 | AK2 | protein_coding | 0.684494444 | 9.89E-07 | 9.01E-09 | 89.61576475 |
| ENSG00000213859 | KCTD11 | protein_coding | 0.683267367 | 0.000122315 | 2.06E-06 | 52.08371274 |
| ENSG00000174428 | GTF2IRD2B | protein_coding | 0.682931521 | 0.004555366 | 0.000198717 | 18.30722833 |
| ENSG00000176890 | TYMS | protein_coding | 0.680611386 | 0.021740582 | 0.001785977 | 66.24665784 |
| ENSG00000095380 | NANS | protein_coding | 0.68049048 | 0.049434897 | 0.006022874 | 23.66603975 |
| ENSG00000108406 | DHX40 | protein_coding | 0.679269026 | 7.24E-06 | 7.88E-08 | 129.1427831 |
| ENSG00000196782 | MAML3 | protein_coding | 0.678493696 | 0.000497432 | 1.12E-05 | 86.54874346 |
| ENSG00000206557 | TRIM71 | protein_coding | 0.677073391 | 0.011481644 | 0.000725138 | 44.81508067 |
| ENSG00000137166 | FOXP4 | protein_coding | 0.676160996 | 0.038663358 | 0.004116489 | 80.46663243 |
| ENSG00000100441 | KHNYN | protein_coding | 0.675856191 | 0.041007754 | 0.004552899 | 67.71808088 |
| ENSG00000125352 | RNF113A | protein_coding | 0.675654195 | 0.006285877 | 0.000303811 | 104.0990674 |
| ENSG00000163629 | PTPN13 | protein_coding | 0.674275546 | 0.007028822 | 0.000359256 | 105.1894973 |
| ENSG00000146197 | SCUBE3 | protein_coding | 0.672851534 | 0.047493861 | 0.005669048 | 44.72376951 |
| ENSG00000188848 | BEND4 | protein_coding | 0.672584022 | 0.013301101 | 0.000892104 | 18.045533 |
| ENSG00000105516 | DBP | protein_coding | 0.67256691 | 0.027031222 | 0.002442651 | 19.58488578 |
| ENSG00000196365 | LONP1 | protein_coding | 0.669744168 | 2.16E-07 | 1.85E-09 | 127.7779367 |
| ENSG00000204128 | C2orf72 | protein_coding | 0.66971991 | 8.81E-05 | 1.42E-06 | 65.7603335 |
| ENSG00000067167 | TRAM1 | protein_coding | 0.663012222 | 0.011264727 | 0.000701001 | 96.98988618 |
| ENSG00000173548 | SNX33 | protein_coding | 0.66222119 | 0.037513814 | 0.003939066 | 23.77947269 |
| ENSG00000134871 | COL4A2 | protein_coding | 0.661856997 | 0.030802106 | 0.002940991 | 242.5242134 |
| ENSG00000183255 | PTTG1IP | protein_coding | 0.65942924 | 0.000141718 | 2.46E-06 | 223.4010042 |
| ENSG00000091127 | PUS7 | protein_coding | 0.658575683 | 0.019040859 | 0.001467169 | 13.44677153 |
| ENSG00000196205 | EEF1A1P5 | processed_pseudogene | 0.656504235 | 0.008343255 | 0.000465089 | 117.767226 |
| ENSG00000114850 | SSR3 | protein_coding | 0.654669586 | 0.000365648 | 7.62E-06 | 182.7070527 |
| ENSG00000196890 | H2BU1 | protein_coding | 0.65449467 | 0.000404188 | 8.71E-06 | 186.9231262 |
| ENSG00000129757 | CDKN1C | protein_coding | 0.652952776 | 0.016498214 | 0.001198643 | 176.8371414 |
| ENSG00000105327 | BBC3 | protein_coding | 0.652250629 | 0.020673696 | 0.001658429 | 40.30354115 |
| ENSG00000167074 | TEF | protein_coding | 0.651994198 | 0.008843419 | 0.000504975 | 19.59729046 |
| ENSG00000220749 | RPL21P28 | processed_pseudogene | 0.651575774 | 0.017489394 | 0.001295658 | 61.25329052 |
| ENSG00000127663 | KDM4B | protein_coding | 0.650307825 | 6.67E-09 | 4.27E-11 | 211.4056443 |
| ENSG00000182253 | SYNM | protein_coding | 0.650156832 | 0.010815587 | 0.000665536 | 24.42408152 |
| ENSG00000068028 | RASSF1 | protein_coding | 0.648550589 | 0.011179955 | 0.000694 | 37.67929313 |
| ENSG00000146021 | KLHL3 | protein_coding | 0.648535237 | 0.000189944 | 3.45E-06 | 29.92462965 |
| ENSG00000038002 | AGA | protein_coding | 0.647103919 | 0.039826872 | 0.004312174 | 11.90790519 |
| ENSG00000177150 | FAM210A | protein_coding | 0.645505539 | 0.000140399 | 2.43E-06 | 37.49767769 |
| ENSG00000114861 | FOXP1 | protein_coding | 0.641091909 | 0.014769255 | 0.001031978 | 38.55637854 |
| ENSG00000166199 | ALKBH3 | protein_coding | 0.639995917 | 0.009330053 | 0.000548191 | 16.32089867 |
| ENSG00000115866 | DARS1 | protein_coding | 0.639902565 | 0.000251153 | 4.89E-06 | 76.43612493 |
| ENSG00000054611 | TBC1D22A | protein_coding | 0.638969025 | 0.000197071 | 3.62E-06 | 26.45281662 |
| ENSG00000132326 | PER2 | protein_coding | 0.638745713 | 7.99E-06 | 8.89E-08 | 42.64755748 |
| ENSG00000171055 | FEZ2 | protein_coding | 0.638549365 | 0.002879339 | 0.000103619 | 34.74545045 |
| ENSG00000116990 | MYCL | protein_coding | 0.638521202 | 0.002231851 | 7.48E-05 | 70.51467241 |
| ENSG00000172071 | EIF2AK3 | protein_coding | 0.637778212 | 0.008081108 | 0.000443612 | 55.71154734 |
| ENSG00000188856 | RPSAP47 | processed_pseudogene | 0.636983652 | 0.020541385 | 0.001631955 | 113.3754612 |
| ENSG00000203644 | AC083799.1 | lncRNA | 0.635945071 | 0.037140685 | 0.003871211 | 22.76426028 |
| ENSG00000005448 | WDR54 | protein_coding | 0.634068281 | 2.84E-05 | 3.73E-07 | 140.124246 |
| ENSG00000241852 | C8orf58 | protein_coding | 0.631810264 | 0.02796861 | 0.002561054 | 19.7390394 |
| ENSG00000178802 | MPI | protein_coding | 0.63014533 | 3.36E-06 | 3.22E-08 | 77.20114615 |
| ENSG00000136010 | ALDH1L2 | protein_coding | 0.629320133 | 0.002274466 | 7.67E-05 | 31.7575003 |
| ENSG00000044115 | CTNNA1 | protein_coding | 0.629038403 | 0.037266841 | 0.003901624 | 256.0720995 |
| ENSG00000130821 | SLC6A8 | protein_coding | 0.624987796 | 0.001278181 | 3.58E-05 | 279.645281 |
| ENSG00000106105 | GARS1 | protein_coding | 0.62369502 | 0.000667796 | 1.60E-05 | 159.8863298 |
| ENSG00000112081 | SRSF3 | protein_coding | 0.621624197 | 3.79E-11 | 1.34E-13 | 388.0767347 |
| ENSG00000179094 | PER1 | protein_coding | 0.621541848 | 0.037810743 | 0.003990693 | 104.693986 |
| ENSG00000114439 | BBX | protein_coding | 0.620850679 | 0.004415453 | 0.000189886 | 142.4162784 |
| ENSG00000151322 | NPAS3 | protein_coding | 0.618154644 | 0.018787536 | 0.001436106 | 108.2294934 |
| ENSG00000212864 | RNF208 | protein_coding | 0.617830689 | 7.29E-06 | 8.00E-08 | 107.7740444 |
| ENSG00000089876 | DHX32 | protein_coding | 0.617504719 | 0.001943202 | 6.23E-05 | 27.48691926 |
| ENSG00000213672 | NCKIPSD | protein_coding | 0.616965726 | 0.000831395 | 2.06E-05 | 43.61577302 |
| ENSG00000133056 | PIK3C2B | protein_coding | 0.615552861 | 1.59E-05 | 1.92E-07 | 141.3477127 |
| ENSG00000151014 | NOCT | protein_coding | 0.614689183 | 0.003643761 | 0.000145165 | 24.83005379 |
| ENSG00000135298 | ADGRB3 | protein_coding | 0.614584253 | 1.59E-06 | 1.48E-08 | 110.0784422 |
| ENSG00000176700 | SCAND2P | unprocessed_pseudogene | 0.613068373 | 0.003907095 | 0.000159276 | 44.44490929 |
| ENSG00000131873 | CHSY1 | protein_coding | 0.613035376 | 0.000770101 | 1.88E-05 | 63.35933695 |
| ENSG00000126804 | ZBTB1 | protein_coding | 0.61290762 | 6.58E-06 | 7.11E-08 | 53.34173066 |
| ENSG00000048028 | USP28 | protein_coding | 0.612682368 | 0.000659219 | 1.58E-05 | 38.1654468 |
| ENSG00000139190 | VAMP1 | protein_coding | 0.609744771 | 0.005705823 | 0.000269608 | 27.4366551 |
| ENSG00000082641 | NFE2L1 | protein_coding | 0.609494989 | 0.012749417 | 0.000840819 | 423.5991008 |
| ENSG00000117305 | HMGCL | protein_coding | 0.606064139 | 0.000978547 | 2.55E-05 | 26.86082591 |
| ENSG00000087074 | PPP1R15A | protein_coding | 0.605402727 | 0.003516205 | 0.00013867 | 282.9414886 |
| ENSG00000124570 | SERPINB6 | protein_coding | 0.605139805 | 0.037266841 | 0.003901512 | 28.43752215 |
| ENSG00000178235 | SLITRK1 | protein_coding | -0.60020802 | 0.032754234 | 0.003193993 | 177.9436995 |
| ENSG00000108960 | MMD | protein_coding | -0.600404338 | 0.004408532 | 0.000189248 | 95.10134694 |
| ENSG00000177511 | ST8SIA3 | protein_coding | -0.602096822 | 0.015028684 | 0.00106092 | 102.0584377 |
| ENSG00000145476 | CYP4V2 | protein_coding | -0.60215286 | 0.013887163 | 0.000943538 | 27.61489242 |
| ENSG00000115266 | APC2 | protein_coding | -0.603855591 | 0.018340016 | 0.001384702 | 379.9248282 |
| ENSG00000127124 | HIVEP3 | protein_coding | -0.60417727 | 0.020544545 | 0.001638551 | 28.20092812 |
| ENSG00000127603 | MACF1 | protein_coding | -0.604291796 | 0.003201872 | 0.000121976 | 809.4831933 |
| ENSG00000109956 | B3GAT1 | protein_coding | -0.606370927 | 0.009556641 | 0.000567407 | 97.58727469 |
| ENSG00000094916 | CBX5 | protein_coding | -0.607822095 | 0.000380444 | 8.04E-06 | 1359.332061 |
| ENSG00000176887 | SOX11 | protein_coding | -0.608355146 | 0.002410744 | 8.25E-05 | 3512.0151 |
| ENSG00000126217 | MCF2L | protein_coding | -0.610042004 | 0.019681616 | 0.001542375 | 151.0031238 |
| ENSG00000126950 | TMEM35A | protein_coding | -0.611090175 | 0.027058926 | 0.002448507 | 87.20755163 |
| ENSG00000114999 | TTL | protein_coding | -0.611274952 | 5.35E-05 | 7.73E-07 | 78.39254892 |
| ENSG00000168913 | ENHO | protein_coding | -0.612040792 | 0.008128819 | 0.00044937 | 58.35456545 |
| ENSG00000197694 | SPTAN1 | protein_coding | -0.612978431 | 0.004949199 | 0.000219718 | 822.1687709 |
| ENSG00000075340 | ADD2 | protein_coding | -0.613017784 | 0.019490744 | 0.001519893 | 211.1907471 |
| ENSG00000166402 | TUB | protein_coding | -0.613208302 | 0.001395632 | 4.07E-05 | 195.4004901 |
| ENSG00000114770 | ABCC5 | protein_coding | -0.61419097 | 0.001395697 | 4.09E-05 | 80.60263986 |
| ENSG00000115525 | ST3GAL5 | protein_coding | -0.615278045 | 0.020368541 | 0.00161036 | 37.50344898 |
| ENSG00000250091 | DNAH10OS | lncRNA | -0.617132785 | 0.013162259 | 0.000880058 | 63.35613777 |
| ENSG00000158882 | TOMM40L | protein_coding | -0.617154874 | 0.005576687 | 0.000260203 | 44.59349408 |
| ENSG00000080824 | HSP90AA1 | protein_coding | -0.617338791 | 6.20E-05 | 9.19E-07 | 1386.179976 |
| ENSG00000108509 | CAMTA2 | protein_coding | -0.617563543 | 0.004423173 | 0.000190735 | 46.04736197 |
| ENSG00000144730 | IL17RD | protein_coding | -0.617811607 | 0.013146999 | 0.000878023 | 98.98586757 |
| ENSG00000197535 | MYO5A | protein_coding | -0.618229524 | 0.02292943 | 0.00193321 | 277.2362263 |
| ENSG00000100604 | CHGA | protein_coding | -0.619315978 | 0.021128257 | 0.001722081 | 77.58017364 |
| ENSG00000106976 | DNM1 | protein_coding | -0.620503033 | 0.002104579 | 6.89E-05 | 156.5827397 |
| ENSG00000178662 | CSRNP3 | protein_coding | -0.621792952 | 0.018938876 | 0.001451359 | 297.5801965 |
| ENSG00000065923 | SLC9A7 | protein_coding | -0.624709689 | 0.01305583 | 0.000869918 | 85.9495341 |
| ENSG00000167378 | IRGQ | protein_coding | -0.625106811 | 0.000380444 | 8.05E-06 | 333.9962569 |
| ENSG00000076641 | PAG1 | protein_coding | -0.627720935 | 0.009423812 | 0.000557723 | 55.94198975 |
| ENSG00000153558 | FBXL2 | protein_coding | -0.627928683 | 0.008885975 | 0.000510436 | 25.18029254 |
| ENSG00000108654 | DDX5 | protein_coding | -0.628886256 | 1.76E-05 | 2.18E-07 | 1415.539926 |
| ENSG00000139970 | RTN1 | protein_coding | -0.632236684 | 0.029666211 | 0.002769146 | 222.2212643 |
| ENSG00000272398 | CD24 | protein_coding | -0.632992602 | 0.027554052 | 0.002514584 | 1389.986136 |
| ENSG00000160796 | NBEAL2 | protein_coding | -0.634063245 | 0.018141271 | 0.001358634 | 18.87772704 |
| ENSG00000169851 | PCDH7 | protein_coding | -0.639042714 | 0.039269377 | 0.004205268 | 44.36594314 |
| ENSG00000163947 | ARHGEF3 | protein_coding | -0.639144688 | 0.047493861 | 0.005651987 | 18.05458879 |
| ENSG00000197147 | LRRC8B | protein_coding | -0.641102004 | 0.008552879 | 0.000480736 | 39.30597808 |
| ENSG00000065413 | ANKRD44 | protein_coding | -0.641926089 | 0.014302432 | 0.000985004 | 29.35420357 |
| ENSG00000068366 | ACSL4 | protein_coding | -0.642716248 | 0.020965788 | 0.001702904 | 95.30445706 |
| ENSG00000112852 | PCDHB2 | protein_coding | -0.64352569 | 0.000857373 | 2.14E-05 | 322.6461858 |
| ENSG00000279495 | AL928654.4 | TEC | -0.644597528 | 0.025608927 | 0.002261937 | 26.36971449 |
| ENSG00000119640 | ACYP1 | protein_coding | -0.644600526 | 0.014636609 | 0.001018189 | 20.83875452 |
| ENSG00000182103 | FAM181B | protein_coding | -0.645202954 | 0.029773685 | 0.002789555 | 49.53474192 |
| ENSG00000197774 | EME2 | protein_coding | -0.647372607 | 0.020733481 | 0.001668027 | 39.82657722 |
| ENSG00000006210 | CX3CL1 | protein_coding | -0.650054049 | 0.014508839 | 0.0010037 | 24.50700269 |
| ENSG00000174599 | TRAM1L1 | protein_coding | -0.650210674 | 0.04476081 | 0.005180393 | 45.78378284 |
| ENSG00000104833 | TUBB4A | protein_coding | -0.65078723 | 0.009113325 | 0.000529828 | 200.0938215 |
| ENSG00000159409 | CELF3 | protein_coding | -0.652984144 | 0.011581057 | 0.00073323 | 331.6807963 |
| ENSG00000164924 | YWHAZ | protein_coding | -0.653986572 | 4.40E-05 | 6.01E-07 | 1151.971643 |
| ENSG00000170837 | GPR27 | protein_coding | -0.65499803 | 0.004282397 | 0.000181189 | 106.0005939 |
| ENSG00000262155 | LINC02175 | lncRNA | -0.656436308 | 0.01514641 | 0.001072364 | 13.67175889 |
| ENSG00000161082 | CELF5 | protein_coding | -0.656489615 | 0.012966415 | 0.00086118 | 138.321731 |
| ENSG00000253305 | PCDHGB6 | protein_coding | -0.657378819 | 0.002470657 | 8.56E-05 | 74.85013022 |
| ENSG00000130294 | KIF1A | protein_coding | -0.658247882 | 0.002663891 | 9.42E-05 | 742.5734404 |
| ENSG00000173898 | SPTBN2 | protein_coding | -0.658585613 | 0.000642881 | 1.52E-05 | 169.3807164 |
| ENSG00000029153 | ARNTL2 | protein_coding | -0.661621807 | 0.045922821 | 0.005357426 | 10.58218794 |
| ENSG00000149218 | ENDOD1 | protein_coding | -0.66266804 | 0.006014126 | 0.000287891 | 19.50964077 |
| ENSG00000183186 | C2CD4C | protein_coding | -0.662731509 | 0.00228044 | 7.73E-05 | 46.82765176 |
| ENSG00000165548 | TMEM63C | protein_coding | -0.663079898 | 0.031017988 | 0.002969604 | 32.00303759 |
| ENSG00000092051 | JPH4 | protein_coding | -0.663099231 | 0.004445465 | 0.000192814 | 190.1803 |
| ENSG00000143494 | VASH2 | protein_coding | -0.663176613 | 0.027011154 | 0.002433757 | 60.35352621 |
| ENSG00000278012 | AL031658.2 | lncRNA | -0.663792979 | 0.044174613 | 0.005092433 | 36.74749451 |
| ENSG00000111670 | GNPTAB | protein_coding | -0.66522092 | 0.002788512 | 9.97E-05 | 73.61354889 |
| ENSG00000100167 | SEPTIN3 | protein_coding | -0.666927193 | 0.001741964 | 5.35E-05 | 717.9564632 |
| ENSG00000164076 | CAMKV | protein_coding | -0.667018982 | 0.024830918 | 0.00217213 | 92.11856786 |
| ENSG00000213190 | MLLT11 | protein_coding | -0.668089823 | 0.00291392 | 0.000106415 | 687.3112567 |
| ENSG00000120694 | HSPH1 | protein_coding | -0.669953396 | 6.83E-05 | 1.06E-06 | 223.5275244 |
| ENSG00000249992 | TMEM158 | protein_coding | -0.67210685 | 0.01029279 | 0.000626024 | 53.31817151 |
| ENSG00000073464 | CLCN4 | protein_coding | -0.672684717 | 0.022473767 | 0.0018705 | 62.15979275 |
| ENSG00000229191 | AL358473.1 | lncRNA | -0.673573247 | 0.009219131 | 0.00054025 | 42.66144558 |
| ENSG00000198668 | CALM1 | protein_coding | -0.673904781 | 0.005325386 | 0.000242997 | 1011.73356 |
| ENSG00000107282 | APBA1 | protein_coding | -0.676034539 | 0.021799016 | 0.001800783 | 68.29288318 |
| ENSG00000169710 | FASN | protein_coding | -0.676303604 | 3.81E-06 | 3.80E-08 | 396.3298803 |
| ENSG00000005379 | TSPOAP1 | protein_coding | -0.677742223 | 0.009349708 | 0.000550259 | 91.26835896 |
| ENSG00000198945 | L3MBTL3 | protein_coding | -0.677849588 | 0.014983512 | 0.001054983 | 23.12881059 |
| ENSG00000267909 | CCDC177 | protein_coding | -0.678934738 | 0.007268417 | 0.000382164 | 46.97674277 |
| ENSG00000178233 | TMEM151B | protein_coding | -0.679258169 | 0.013445083 | 0.000904159 | 98.01613562 |
| ENSG00000131711 | MAP1B | protein_coding | -0.681822321 | 0.000362368 | 7.47E-06 | 3448.831211 |
| ENSG00000101347 | SAMHD1 | protein_coding | -0.681831722 | 0.003157707 | 0.00011995 | 19.41018577 |
| ENSG00000116983 | HPCAL4 | protein_coding | -0.682445255 | 0.037137062 | 0.003867966 | 142.113285 |
| ENSG00000163412 | EIF4E3 | protein_coding | -0.682909145 | 0.004168343 | 0.000175719 | 28.34539503 |
| ENSG00000198720 | ANKRD13B | protein_coding | -0.683404186 | 4.25E-05 | 5.75E-07 | 96.3276708 |
| ENSG00000132024 | CC2D1A | protein_coding | -0.685496273 | 0.004325374 | 0.000184009 | 42.0929022 |
| ENSG00000168824 | NSG1 | protein_coding | -0.685532581 | 0.020544545 | 0.001634338 | 154.9055883 |
| ENSG00000135074 | ADAM19 | protein_coding | -0.686035894 | 0.008087808 | 0.000444605 | 25.77964893 |
| ENSG00000125966 | MMP24 | protein_coding | -0.686762713 | 0.048099875 | 0.005781279 | 73.25582308 |
| ENSG00000167114 | SLC27A4 | protein_coding | -0.687018057 | 0.00232389 | 7.93E-05 | 28.92910635 |
| ENSG00000255690 | TRIL | protein_coding | -0.687461204 | 0.047346616 | 0.005615376 | 24.06435454 |
| ENSG00000266173 | STRADA | protein_coding | -0.687891564 | 0.024254336 | 0.002103468 | 9.344944386 |
| ENSG00000134313 | KIDINS220 | protein_coding | -0.690035506 | 0.005625229 | 0.000264929 | 503.2939553 |
| ENSG00000155893 | PXYLP1 | protein_coding | -0.693644056 | 0.000229032 | 4.37E-06 | 50.4324175 |
| ENSG00000166689 | PLEKHA7 | protein_coding | -0.695473012 | 0.001056801 | 2.81E-05 | 41.43662664 |
| ENSG00000197102 | DYNC1H1 | protein_coding | -0.697485653 | 0.000676409 | 1.63E-05 | 1250.799942 |
| ENSG00000171793 | CTPS1 | protein_coding | -0.700848359 | 4.48E-06 | 4.53E-08 | 61.8340751 |
| ENSG00000186469 | GNG2 | protein_coding | -0.702844591 | 0.003358704 | 0.000130178 | 393.1680806 |
| ENSG00000246922 | UBAP1L | protein_coding | -0.70375404 | 0.025744654 | 0.002275913 | 22.42036269 |
| ENSG00000128512 | DOCK4 | protein_coding | -0.705195665 | 0.005376807 | 0.000246812 | 68.59108496 |
| ENSG00000152092 | ASTN1 | protein_coding | -0.706312012 | 0.006565635 | 0.000322447 | 116.4601611 |
| ENSG00000167065 | DUSP18 | protein_coding | -0.706372231 | 0.018055334 | 0.001350804 | 24.86150444 |
| ENSG00000130477 | UNC13A | protein_coding | -0.706598577 | 0.003663833 | 0.000146248 | 129.2603324 |
| ENSG00000145016 | RUBCN | protein_coding | -0.707862227 | 0.000625864 | 1.48E-05 | 65.72236958 |
| ENSG00000146352 | CLVS2 | protein_coding | -0.708374895 | 0.015592777 | 0.001114802 | 51.71391601 |
| ENSG00000158445 | KCNB1 | protein_coding | -0.708707961 | 0.002466039 | 8.53E-05 | 62.32299097 |
| ENSG00000143061 | IGSF3 | protein_coding | -0.710696153 | 0.000224545 | 4.25E-06 | 103.7785441 |
| ENSG00000084731 | KIF3C | protein_coding | -0.711710517 | 0.000543333 | 1.25E-05 | 239.1334716 |
| ENSG00000162174 | ASRGL1 | protein_coding | -0.713419697 | 0.009751033 | 0.000583466 | 28.99190164 |
| ENSG00000184672 | RALYL | protein_coding | -0.713907859 | 0.041448657 | 0.004608251 | 20.24904603 |
| ENSG00000104435 | STMN2 | protein_coding | -0.71555494 | 0.039780321 | 0.004299361 | 725.534418 |
| ENSG00000242294 | STAG3L5P | unprocessed_pseudogene | -0.715866412 | 0.029687299 | 0.002773443 | 19.35561709 |
| ENSG00000151150 | ANK3 | protein_coding | -0.717234212 | 0.007647558 | 0.000409775 | 503.8038167 |
| ENSG00000151229 | SLC2A13 | protein_coding | -0.719780964 | 0.016096503 | 0.001160758 | 45.25475829 |
| ENSG00000165434 | PGM2L1 | protein_coding | -0.720440233 | 0.01704125 | 0.001252569 | 209.7976945 |
| ENSG00000267632 | AC067852.3 | lncRNA | -0.720586235 | 0.047983567 | 0.005749729 | 19.77873516 |
| ENSG00000274588 | DGKK | protein_coding | -0.721688935 | 0.048317744 | 0.005830809 | 19.22384186 |
| ENSG00000176593 | AC008969.1 | lncRNA | -0.722402912 | 0.008965043 | 0.000517746 | 65.56607549 |
| ENSG00000019505 | SYT13 | protein_coding | -0.724692284 | 0.005852936 | 0.000277915 | 177.8418711 |
| ENSG00000156103 | MMP16 | protein_coding | -0.726077431 | 0.002589526 | 9.06E-05 | 121.6058481 |
| ENSG00000063015 | SEZ6 | protein_coding | -0.727276287 | 0.01029279 | 0.000626214 | 41.63151272 |
| ENSG00000155511 | GRIA1 | protein_coding | -0.728394666 | 0.010858692 | 0.000669865 | 114.3186265 |
| ENSG00000186868 | MAPT | protein_coding | -0.728854247 | 0.033275726 | 0.003298798 | 493.0981229 |
| ENSG00000278864 | AC055811.4 | TEC | -0.729767474 | 0.005625229 | 0.000264579 | 49.56409331 |
| ENSG00000170091 | NSG2 | protein_coding | -0.731168382 | 0.018368194 | 0.001388393 | 381.7153249 |
| ENSG00000102879 | CORO1A | protein_coding | -0.731702147 | 0.031571201 | 0.00302988 | 31.79443241 |
| ENSG00000158528 | PPP1R9A | protein_coding | -0.73174758 | 0.000398214 | 8.55E-06 | 108.2584427 |
| ENSG00000143434 | SEMA6C | protein_coding | -0.732126812 | 0.002906384 | 0.000105691 | 103.590508 |
| ENSG00000141098 | GFOD2 | protein_coding | -0.732281826 | 0.002288577 | 7.77E-05 | 79.25783432 |
| ENSG00000173452 | TMEM196 | protein_coding | -0.733533682 | 0.045770195 | 0.005329019 | 23.17380761 |
| ENSG00000005249 | PRKAR2B | protein_coding | -0.734920277 | 0.022817597 | 0.001921395 | 118.539865 |
| ENSG00000198743 | SLC5A3 | protein_coding | -0.736006529 | 7.88E-08 | 6.21E-10 | 133.5396856 |
| ENSG00000160963 | COL26A1 | protein_coding | -0.737425245 | 0.033314409 | 0.003305205 | 22.05212044 |
| ENSG00000259834 | AL365361.1 | lncRNA | -0.73750518 | 0.007524391 | 0.000398437 | 54.95819038 |
| ENSG00000116544 | DLGAP3 | protein_coding | -0.738651246 | 0.040713872 | 0.004488836 | 28.3141367 |
| ENSG00000148541 | FAM13C | protein_coding | -0.738867925 | 0.013688987 | 0.000923732 | 33.69037195 |
| ENSG00000172020 | GAP43 | protein_coding | -0.739186 | 0.004287017 | 0.000182046 | 179.0728724 |
| ENSG00000143324 | XPR1 | protein_coding | -0.739835169 | 0.007555425 | 0.000402505 | 653.6247262 |
| ENSG00000168970 | JMJD7-PLA2G4B | protein_coding | -0.742286979 | 0.019490744 | 0.001518745 | 12.51574458 |
| ENSG00000279207 | AC015813.6 | TEC | -0.743824599 | 0.012229777 | 0.000792216 | 37.2277543 |
| ENSG00000270885 | RASL10B | protein_coding | -0.746010189 | 0.001889829 | 5.98E-05 | 38.11874114 |
| ENSG00000167037 | SGSM1 | protein_coding | -0.747212127 | 0.03309759 | 0.003263009 | 20.94632617 |
| ENSG00000128482 | RNF112 | protein_coding | -0.7472137 | 0.012925298 | 0.000856231 | 22.53010576 |
| ENSG00000158555 | GDPD5 | protein_coding | -0.748166743 | 0.045004328 | 0.005219001 | 20.04761924 |
| ENSG00000110436 | SLC1A2 | protein_coding | -0.748235009 | 0.024593911 | 0.002145701 | 166.9070307 |
| ENSG00000174460 | ZCCHC12 | protein_coding | -0.749273826 | 0.043590735 | 0.004999114 | 323.9829973 |
| ENSG00000276550 | HERC2P2 | unprocessed_pseudogene | -0.751795809 | 0.000369219 | 7.73E-06 | 83.4983578 |
| ENSG00000077264 | PAK3 | protein_coding | -0.751802729 | 0.015224455 | 0.001081416 | 291.7273579 |
| ENSG00000133083 | DCLK1 | protein_coding | -0.752733174 | 0.001614529 | 4.91E-05 | 501.367532 |
| ENSG00000107954 | NEURL1 | protein_coding | -0.754111649 | 0.043001524 | 0.004891061 | 29.0979845 |
| ENSG00000113657 | DPYSL3 | protein_coding | -0.756384187 | 0.013874802 | 0.000940556 | 1279.969939 |
| ENSG00000204681 | GABBR1 | protein_coding | -0.761840066 | 0.002890916 | 0.000104905 | 232.9165447 |
| ENSG00000100095 | SEZ6L | protein_coding | -0.764838065 | 0.003064663 | 0.00011546 | 259.4281992 |
| ENSG00000138604 | GLCE | protein_coding | -0.766049234 | 0.038506644 | 0.004080456 | 51.63688685 |
| ENSG00000164742 | ADCY1 | protein_coding | -0.766225271 | 0.016853758 | 0.00123098 | 84.28015224 |
| ENSG00000135454 | B4GALNT1 | protein_coding | -0.768680161 | 0.009025518 | 0.000522192 | 112.8877913 |
| ENSG00000102678 | FGF9 | protein_coding | -0.771493079 | 0.048538975 | 0.005861254 | 21.44211981 |
| ENSG00000166257 | SCN3B | protein_coding | -0.773714885 | 0.00514576 | 0.000231251 | 285.0919665 |
| ENSG00000185634 | SHC4 | protein_coding | -0.774414961 | 0.026050511 | 0.002314563 | 10.47449542 |
| ENSG00000182256 | GABRG3 | protein_coding | -0.776157971 | 0.03066509 | 0.002905958 | 30.57226702 |
| ENSG00000253846 | PCDHGA10 | protein_coding | -0.780285226 | 0.042530173 | 0.0047969 | 11.38207847 |
| ENSG00000138083 | SIX3 | protein_coding | -0.782059533 | 0.039826872 | 0.004314168 | 190.7179764 |
| ENSG00000165194 | PCDH19 | protein_coding | -0.783403032 | 0.0075874 | 0.000404794 | 108.210795 |
| ENSG00000136040 | PLXNC1 | protein_coding | -0.784362593 | 0.013301101 | 0.000892423 | 102.7118954 |
| ENSG00000280287 | AC131212.3 | TEC | -0.787715261 | 0.010262674 | 0.000622797 | 16.22414243 |
| ENSG00000177272 | KCNA3 | protein_coding | -0.788569934 | 0.012590785 | 0.000827267 | 40.78778351 |
| ENSG00000131378 | RFTN1 | protein_coding | -0.790746988 | 0.014368824 | 0.000991795 | 26.50750934 |
| ENSG00000273301 | AC016717.2 | lncRNA | -0.79270525 | 0.006301247 | 0.000306013 | 37.57932516 |
| ENSG00000156140 | ADAMTS3 | protein_coding | -0.793303501 | 0.049663343 | 0.006054966 | 11.23975951 |
| ENSG00000221995 | TIAF1 | protein_coding | -0.796929614 | 0.012233242 | 0.000793385 | 15.02551317 |
| ENSG00000175048 | ZDHHC14 | protein_coding | -0.797778385 | 0.00127038 | 3.55E-05 | 11.10359085 |
| ENSG00000018236 | CNTN1 | protein_coding | -0.798477355 | 0.048099875 | 0.00577601 | 214.7164035 |
| ENSG00000151690 | MFSD6 | protein_coding | -0.801252021 | 0.005576687 | 0.000260215 | 18.66256179 |
| ENSG00000198105 | ZNF248 | protein_coding | -0.801810375 | 0.024190309 | 0.002086209 | 43.34651439 |
| ENSG00000270059 | AC121493.1 | lncRNA | -0.802151194 | 0.016085814 | 0.001158745 | 46.02373239 |
| ENSG00000182463 | TSHZ2 | protein_coding | -0.812869988 | 0.001256177 | 3.46E-05 | 85.16556409 |
| ENSG00000054793 | ATP9A | protein_coding | -0.815257876 | 0.000515216 | 1.18E-05 | 446.0085059 |
| ENSG00000066382 | MPPED2 | protein_coding | -0.815816278 | 0.022312728 | 0.001851449 | 55.08277282 |
| ENSG00000198216 | CACNA1E | protein_coding | -0.816044601 | 0.019646352 | 0.001538095 | 99.1406809 |
| ENSG00000162512 | SDC3 | protein_coding | -0.818586011 | 5.19E-05 | 7.41E-07 | 478.7758555 |
| ENSG00000135439 | AGAP2 | protein_coding | -0.820811338 | 0.014607655 | 0.001012791 | 121.5797725 |
| ENSG00000140323 | DISP2 | protein_coding | -0.822013709 | 0.026944465 | 0.002418185 | 81.08286307 |
| ENSG00000064687 | ABCA7 | protein_coding | -0.822798721 | 0.00834766 | 0.000465979 | 32.08449861 |
| ENSG00000260630 | SNAI3-AS1 | lncRNA | -0.826946364 | 0.004074246 | 0.000169195 | 21.33013699 |
| ENSG00000091844 | RGS17 | protein_coding | -0.82755429 | 0.039977334 | 0.004342813 | 35.91106063 |
| ENSG00000144460 | NYAP2 | protein_coding | -0.828990283 | 0.011330863 | 0.000712849 | 12.80634536 |
| ENSG00000061918 | GUCY1B1 | protein_coding | -0.832948307 | 0.030542775 | 0.002886377 | 21.99471339 |
| ENSG00000136352 | NKX2-1 | protein_coding | -0.836939448 | 0.009751033 | 0.000583014 | 374.5962788 |
| ENSG00000145248 | SLC10A4 | protein_coding | -0.839461536 | 0.025442054 | 0.002239341 | 8.792199142 |
| ENSG00000162630 | B3GALT2 | protein_coding | -0.84008503 | 0.021799016 | 0.001797263 | 56.66049754 |
| ENSG00000147231 | RADX | protein_coding | -0.841047505 | 0.001257003 | 3.47E-05 | 67.58091024 |
| ENSG00000265800 | AC022211.4 | lncRNA | -0.851133135 | 0.043001524 | 0.004890448 | 8.708299795 |
| ENSG00000110881 | ASIC1 | protein_coding | -0.851638573 | 0.017282988 | 0.001275675 | 109.7771546 |
| ENSG00000180616 | SSTR2 | protein_coding | -0.853261774 | 0.045782174 | 0.005333948 | 18.06567757 |
| ENSG00000171724 | VAT1L | protein_coding | -0.85521031 | 0.003634288 | 0.000144507 | 118.2306062 |
| ENSG00000184828 | ZBTB7C | protein_coding | -0.855278255 | 0.007188146 | 0.000374059 | 13.17985375 |
| ENSG00000136367 | ZFHX2 | protein_coding | -0.858259141 | 0.000192928 | 3.52E-06 | 76.60479693 |
| ENSG00000271327 | AC010201.2 | lncRNA | -0.858926974 | 0.010905951 | 0.000673791 | 16.94837958 |
| ENSG00000164604 | GPR85 | protein_coding | -0.860981491 | 0.006839845 | 0.000345909 | 62.79968022 |
| ENSG00000179841 | AKAP5 | protein_coding | -0.862015704 | 0.029360201 | 0.002731551 | 11.49032213 |
| ENSG00000117602 | RCAN3 | protein_coding | -0.862211965 | 0.005376807 | 0.000247004 | 68.90814807 |
| ENSG00000185432 | METTL7A | protein_coding | -0.865974078 | 0.037710978 | 0.003971416 | 9.54583169 |
| ENSG00000007237 | GAS7 | protein_coding | -0.866780161 | 0.000351356 | 7.16E-06 | 84.91218951 |
| ENSG00000176749 | CDK5R1 | protein_coding | -0.870538116 | 4.48E-06 | 4.49E-08 | 521.4956178 |
| ENSG00000120875 | DUSP4 | protein_coding | -0.870920547 | 0.002024986 | 6.54E-05 | 333.2537062 |
| ENSG00000125170 | DOK4 | protein_coding | -0.872724339 | 0.004395197 | 0.000187658 | 67.59207812 |
| ENSG00000111846 | GCNT2 | protein_coding | -0.872753532 | 0.000659219 | 1.57E-05 | 21.98190946 |
| ENSG00000121904 | CSMD2 | protein_coding | -0.876924391 | 0.011190421 | 0.000695513 | 66.95082106 |
| ENSG00000139318 | DUSP6 | protein_coding | -0.877492786 | 0.01380319 | 0.00093357 | 149.9077979 |
| ENSG00000160190 | SLC37A1 | protein_coding | -0.879173624 | 0.000810759 | 2.00E-05 | 29.75804389 |
| ENSG00000164484 | TMEM200A | protein_coding | -0.881403972 | 0.011313431 | 0.000706653 | 28.14287901 |
| ENSG00000153234 | NR4A2 | protein_coding | -0.88352572 | 0.036859506 | 0.003832389 | 22.15888064 |
| ENSG00000187134 | AKR1C1 | protein_coding | -0.884699056 | 0.007608135 | 0.000407075 | 31.2072136 |
| ENSG00000176463 | SLCO3A1 | protein_coding | -0.888312582 | 0.005042355 | 0.000225073 | 19.21015929 |
| ENSG00000188766 | SPRED3 | protein_coding | -0.891332053 | 0.000881213 | 2.23E-05 | 60.36851669 |
| ENSG00000253953 | PCDHGB4 | protein_coding | -0.894483265 | 0.031011431 | 0.002966581 | 18.7614861 |
| ENSG00000268670 | AC016586.1 | lncRNA | -0.902946775 | 0.008899159 | 0.00051276 | 13.78605768 |
| ENSG00000156475 | PPP2R2B | protein_coding | -0.903987797 | 0.004074246 | 0.000169236 | 91.76352096 |
| ENSG00000136531 | SCN2A | protein_coding | -0.906456293 | 0.010232681 | 0.000620186 | 104.7215682 |
| ENSG00000198429 | ZNF69 | protein_coding | -0.907123801 | 0.00137734 | 4.00E-05 | 20.65299681 |
| ENSG00000041353 | RAB27B | protein_coding | -0.914153674 | 0.041992078 | 0.004684879 | 22.49178833 |
| ENSG00000170500 | LONRF2 | protein_coding | -0.917122728 | 4.84E-05 | 6.76E-07 | 188.9353366 |
| ENSG00000140557 | ST8SIA2 | protein_coding | -0.91836893 | 0.000659219 | 1.58E-05 | 120.2244133 |
| ENSG00000148082 | SHC3 | protein_coding | -0.91877304 | 5.52E-05 | 8.06E-07 | 150.0905533 |
| ENSG00000128564 | VGF | protein_coding | -0.92190292 | 0.007703207 | 0.000413946 | 385.6095397 |
| ENSG00000113100 | CDH9 | protein_coding | -0.928232374 | 0.04318299 | 0.004924434 | 9.577499864 |
| ENSG00000279700 | AC131212.2 | TEC | -0.929819671 | 0.015863678 | 0.001139069 | 19.78568284 |
| ENSG00000136928 | GABBR2 | protein_coding | -0.930799357 | 0.015815032 | 0.001133134 | 52.49004333 |
| ENSG00000179542 | SLITRK4 | protein_coding | -0.935160898 | 0.020544545 | 0.001638028 | 69.02183306 |
| ENSG00000156687 | UNC5D | protein_coding | -0.936974586 | 0.033199283 | 0.003280967 | 41.54568638 |
| ENSG00000138944 | SHISAL1 | protein_coding | -0.937444469 | 0.004071252 | 0.000168169 | 76.89720442 |
| ENSG00000242265 | PEG10 | protein_coding | -0.937467558 | 0.001767155 | 5.48E-05 | 3938.142468 |
| ENSG00000187601 | MAGEH1 | protein_coding | -0.937564377 | 0.014293141 | 0.000979761 | 262.9721521 |
| ENSG00000152954 | NRSN1 | protein_coding | -0.939606969 | 0.003728258 | 0.000149407 | 26.04003067 |
| ENSG00000157103 | SLC6A1 | protein_coding | -0.943639204 | 0.000729498 | 1.77E-05 | 78.8748418 |
| ENSG00000280057 | AL022069.2 | TEC | -0.943769614 | 0.043509117 | 0.004978421 | 12.33056882 |
| ENSG00000280234 | AC124303.2 | TEC | -0.944458918 | 0.002498871 | 8.69E-05 | 82.68159329 |
| ENSG00000164976 | MYORG | protein_coding | -0.946826048 | 5.19E-05 | 7.42E-07 | 31.77534395 |
| ENSG00000166501 | PRKCB | protein_coding | -0.950539978 | 0.040788803 | 0.004509695 | 109.2392824 |
| ENSG00000198680 | TUSC1 | protein_coding | -0.950864849 | 0.020673696 | 0.001657261 | 32.03309031 |
| ENSG00000165023 | DIRAS2 | protein_coding | -0.951380014 | 0.000917216 | 2.34E-05 | 38.96689684 |
| ENSG00000171587 | DSCAM | protein_coding | -0.953370392 | 0.01253752 | 0.000822799 | 41.97429121 |
| ENSG00000261754 | AC008555.1 | lncRNA | -0.955684564 | 0.000486845 | 1.09E-05 | 53.44045574 |
| ENSG00000070808 | CAMK2A | protein_coding | -0.956175124 | 0.000865064 | 2.17E-05 | 33.50087378 |
| ENSG00000120645 | IQSEC3 | protein_coding | -0.961716304 | 0.007978694 | 0.0004331 | 39.51138898 |
| ENSG00000144583 | MARCHF4 | protein_coding | -0.974964342 | 0.002879339 | 0.000103818 | 57.51830831 |
| ENSG00000198899 | MT-ATP6 | protein_coding | -0.979604151 | 2.47E-14 | 4.58E-17 | 5077.239749 |
| ENSG00000102466 | FGF14 | protein_coding | -0.98457612 | 0.034804375 | 0.003496023 | 157.4832015 |
| ENSG00000151632 | AKR1C2 | protein_coding | -0.985645136 | 0.002088839 | 6.79E-05 | 23.89144827 |
| ENSG00000175040 | CHST2 | protein_coding | -0.998705574 | 0.005238753 | 0.000238235 | 45.17786848 |
| ENSG00000205929 | C21orf62 | protein_coding | -1.003342855 | 0.048676891 | 0.005889182 | 8.354150725 |
| ENSG00000171551 | ECEL1 | protein_coding | -1.005358788 | 0.023256077 | 0.001972833 | 93.99390498 |
| ENSG00000184564 | SLITRK6 | protein_coding | -1.005860308 | 0.042530173 | 0.004793776 | 30.52749537 |
| ENSG00000198763 | MT-ND2 | protein_coding | -1.007693582 | 4.46E-11 | 1.62E-13 | 5041.71861 |
| ENSG00000165521 | EML5 | protein_coding | -1.009132647 | 0.029784331 | 0.002794006 | 29.53145605 |
| ENSG00000212907 | MT-ND4L | protein_coding | -1.015248002 | 1.54E-05 | 1.85E-07 | 855.0577254 |
| ENSG00000169213 | RAB3B | protein_coding | -1.016925391 | 0.001041922 | 2.75E-05 | 157.9710809 |
| ENSG00000168016 | TRANK1 | protein_coding | -1.023977993 | 0.002723017 | 9.65E-05 | 19.13694269 |
| ENSG00000196189 | SEMA4A | protein_coding | -1.02572234 | 0.011427427 | 0.000720831 | 13.59420167 |
| ENSG00000117600 | PLPPR4 | protein_coding | -1.025738487 | 0.025514793 | 0.002249683 | 35.66975273 |
| ENSG00000266777 | SH3GL1P1 | processed_pseudogene | -1.029301026 | 0.008385177 | 0.000470015 | 10.45968276 |
| ENSG00000242419 | PCDHGC4 | protein_coding | -1.03511001 | 0.000150693 | 2.66E-06 | 17.57269171 |
| ENSG00000273274 | ZBTB8B | protein_coding | -1.037983798 | 0.002570801 | 8.97E-05 | 52.11620032 |
| ENSG00000187325 | TAF9B | protein_coding | -1.038755268 | 0.004079599 | 0.000169773 | 33.35078465 |
| ENSG00000184515 | BEX5 | protein_coding | -1.040020303 | 0.009079016 | 0.000527132 | 60.40003189 |
| ENSG00000011677 | GABRA3 | protein_coding | -1.041385361 | 0.002643854 | 9.31E-05 | 55.30131784 |
| ENSG00000187678 | SPRY4 | protein_coding | -1.045602526 | 0.000940794 | 2.40E-05 | 33.77786139 |
| ENSG00000165566 | AMER2 | protein_coding | -1.0463927 | 3.45E-06 | 3.33E-08 | 348.3152638 |
| ENSG00000120738 | EGR1 | protein_coding | -1.055439613 | 0.007188146 | 0.000373557 | 54.98935136 |
| ENSG00000142686 | C1orf216 | protein_coding | -1.059826478 | 0.006581576 | 0.000326233 | 31.90399174 |
| ENSG00000162975 | KCNF1 | protein_coding | -1.060869316 | 0.001196236 | 3.23E-05 | 35.42299484 |
| ENSG00000173210 | ABLIM3 | protein_coding | -1.061367281 | 0.01517587 | 0.001076793 | 32.58836092 |
| ENSG00000155265 | GOLGA7B | protein_coding | -1.064674775 | 0.022587282 | 0.001893795 | 31.95033682 |
| ENSG00000158258 | CLSTN2 | protein_coding | -1.067677406 | 0.005232289 | 0.000237538 | 86.93639194 |
| ENSG00000116852 | KIF21B | protein_coding | -1.07159637 | 0.0001718 | 3.10E-06 | 592.222944 |
| ENSG00000102924 | CBLN1 | protein_coding | -1.080578144 | 0.042392658 | 0.004759781 | 38.22235099 |
| ENSG00000225630 | MTND2P28 | pseudogene | -1.085139015 | 0.012353535 | 0.000805002 | 143.1878844 |
| ENSG00000144596 | GRIP2 | protein_coding | -1.085888805 | 0.02165112 | 0.001773613 | 31.32413394 |
| ENSG00000124313 | IQSEC2 | protein_coding | -1.087141217 | 0.001262819 | 3.50E-05 | 27.37446362 |
| ENSG00000175264 | CHST1 | protein_coding | -1.090492381 | 0.000378622 | 7.95E-06 | 23.66234224 |
| ENSG00000223551 | TMSB4XP4 | processed_pseudogene | -1.098270437 | 0.009128996 | 0.000532149 | 12.85246565 |
| ENSG00000087495 | PHACTR3 | protein_coding | -1.098352431 | 0.011281319 | 0.000703776 | 26.83883124 |
| ENSG00000234345 | AC234782.2 | processed_pseudogene | -1.104389215 | 0.023394085 | 0.001995867 | 19.15037921 |
| ENSG00000273108 | AL121929.3 | lncRNA | -1.108623462 | 0.022422452 | 0.001862767 | 25.79693278 |
| ENSG00000273079 | GRIN2B | protein_coding | -1.115472022 | 0.003967102 | 0.000162335 | 256.6415327 |
| ENSG00000145247 | OCIAD2 | protein_coding | -1.116030122 | 0.019343777 | 0.001497978 | 21.93646009 |
| ENSG00000186642 | PDE2A | protein_coding | -1.122863389 | 0.022816364 | 0.001918393 | 11.71057588 |
| ENSG00000184905 | TCEAL2 | protein_coding | -1.162072472 | 0.008843419 | 0.000505943 | 26.59122597 |
| ENSG00000080644 | CHRNA3 | protein_coding | -1.172415523 | 0.036859506 | 0.003833096 | 9.06341296 |
| ENSG00000074211 | PPP2R2C | protein_coding | -1.203476547 | 0.008039665 | 0.000438854 | 50.77353054 |
| ENSG00000128872 | TMOD2 | protein_coding | -1.208461533 | 1.51E-08 | 1.03E-10 | 265.8903051 |
| ENSG00000152932 | RAB3C | protein_coding | -1.211598248 | 1.92E-05 | 2.40E-07 | 353.0033466 |
| ENSG00000165186 | PTCHD1 | protein_coding | -1.217738476 | 0.013146999 | 0.000877645 | 25.77054048 |
| ENSG00000103044 | HAS3 | protein_coding | -1.236230252 | 0.000221527 | 4.16E-06 | 26.63538831 |
| ENSG00000205856 | C22orf42 | protein_coding | -1.238925539 | 0.040713872 | 0.004486114 | 15.65979869 |
| ENSG00000270953 | AC007938.3 | lncRNA | -1.245500903 | 0.022560863 | 0.001883705 | 396.4396771 |
| ENSG00000198739 | LRRTM3 | protein_coding | -1.253586574 | 0.007118352 | 0.00036603 | 17.27612385 |
| ENSG00000132975 | GPR12 | protein_coding | -1.275352118 | 0.016060206 | 0.00115442 | 54.78641454 |
| ENSG00000274956 | NKAIN3-IT1 | lncRNA | -1.284747906 | 0.006164441 | 0.000296038 | 47.67559988 |
| ENSG00000221630 | MIR1179 | miRNA | -1.310660901 | 0.030037406 | 0.002829342 | 9.099708689 |
| ENSG00000135824 | RGS8 | protein_coding | -1.322336212 | 0.002215407 | 7.38E-05 | 34.42000216 |
| ENSG00000183908 | LRRC55 | protein_coding | -1.338432053 | 0.000122127 | 2.03E-06 | 60.70122131 |
| ENSG00000188517 | COL25A1 | protein_coding | -1.340485402 | 0.008773344 | 0.000499224 | 18.16934139 |
| ENSG00000198883 | PNMA5 | protein_coding | -1.347521594 | 0.019646352 | 0.001536373 | 11.61837614 |
| ENSG00000210117 | MT-TW | Mt_tRNA | -1.446983788 | 0.017591752 | 0.001307943 | 15.72187853 |
| ENSG00000116254 | CHD5 | protein_coding | -1.468744699 | 0.007946962 | 0.000430726 | 38.75447678 |
| ENSG00000198840 | MT-ND3 | protein_coding | -1.495528323 | 5.46E-17 | 5.06E-20 | 2158.272751 |
| ENSG00000198948 | MFAP3L | protein_coding | -1.516941186 | 0.000775047 | 1.90E-05 | 9.700905667 |
| ENSG00000112038 | OPRM1 | protein_coding | -1.544068897 | 0.028075389 | 0.002574168 | 13.13781863 |
| ENSG00000174145 | NWD2 | protein_coding | -1.546376715 | 0.023170282 | 0.001960672 | 8.845361683 |
| ENSG00000198400 | NTRK1 | protein_coding | -1.54977278 | 0.02043442 | 0.001619897 | 25.9926564 |
| ENSG00000162595 | DIRAS3 | protein_coding | -1.552293505 | 0.003805426 | 0.000153663 | 30.34159846 |
| ENSG00000248527 | MTATP6P1 | unprocessed_pseudogene | -1.57616242 | 0.000964872 | 2.50E-05 | 291.6307677 |
| ENSG00000163285 | GABRG1 | protein_coding | -1.603820761 | 0.021612573 | 0.001767118 | 11.03080557 |
| ENSG00000027644 | INSRR | protein_coding | -1.665485595 | 0.011029592 | 0.000682963 | 34.30693968 |
| ENSG00000168959 | GRM5 | protein_coding | -1.689507319 | 0.012229777 | 0.000792206 | 35.346747 |
| ENSG00000255836 | AC131206.1 | processed_pseudogene | -1.690231302 | 0.01271025 | 0.000837079 | 11.05531427 |
| ENSG00000286214 | AUXG01000058.1 | lncRNA | -1.724269907 | 0.007130638 | 0.000367212 | 674.2804126 |
| ENSG00000117152 | RGS4 | protein_coding | -2.004797365 | 6.58E-05 | 9.90E-07 | 16.75346885 |
| ENSG00000210195 | MT-TT | Mt_tRNA | -2.222042025 | 1.50E-05 | 1.78E-07 | 9.639353259 |
| ENSG00000079689 | SCGN | protein_coding | -2.274450893 | 0.017067983 | 0.001257151 | 11.98765259 |
| ENSG00000210196 | MT-TP | Mt_tRNA | -2.391985922 | 1.48E-16 | 1.60E-19 | 61.98408144 |
| ENSG00000198695 | MT-ND6 | protein_coding | -2.542733456 | 2.36E-22 | 1.82E-26 | 369.7526992 |
| ENSG00000210107 | MT-TQ | Mt_tRNA | -3.191895797 | 2.24E-07 | 1.94E-09 | 14.722918 |
